# FAN1 nuclease processes and pauses on disease-associated slipped-DNA repeats: Mechanism against repeat expansions

**DOI:** 10.1101/2021.04.15.439995

**Authors:** Amit Laxmikant Deshmukh, Marie-Christine Caron, Mohiuddin Mohiuddin, Stella Lanni, Gagan B. Panigrahi, Mahreen Khan, Worrawat Engchuan, Natalie Shum, Aisha Faruqui, Peixiang Wang, Ryan K.C. Yuen, Masayuki Nakamori, Kazuhiko Nakatani, Jean-Yves Masson, Christopher E. Pearson

## Abstract

FAN1 nuclease is a modifier of repeat expansion diseases, including Huntington’s disease (HD), fragile X syndrome, and autism. The age of HD onset correlates with ongoing ‘inchworm-like’ repeat expansions (1-3 CAG units/event) in HD brains, and is regulated by three modifiers: The first two, repeat tract length and purity exert their effects by enhancing and slowing CAG expansions, respectively, by affecting the formation of slipped-DNAs — mutagenic intermediates of instability; which are processed to expansions by the third modifiers, DNA repair proteins. FAN1 protects against hyper-expansions of repeats, by unknown mechanisms. We show FAN1, through iterative cycles bound, dimerized and cleaved slipped-DNAs, yielding striking patterns of distinct *exo*-nuclease *pauses* along slip-outs; 5′-C↓A↓GC↓A↓G-3′ and 5′-C↓T↓G↓C↓T↓G-3′. The transcriptionally-displaced CAG strand was excised slower than its complementary CTG strand, required A•A and T•T mismatches, as fully-paired hairpins *arrested* excision progression, while disease-delaying ***CAA*** interruptions further slowed FAN1 excision. In contrast, *endo*-nucleolytic cleavage was insensitive to slip-outs. Rare FAN1 variants were found in autism individuals with CGG/CCG repeat expansions. Excision of CGG/CCG slip-outs were similarly excised, with CGG being slower than CCG. The slip-out specific ligand, Naphthyridine-Azaquinolone, shown to induce contractions of expanded repeats in cells, required FAN1 for its effect, and protected slip-outs from FAN1’s *exo*- but not *endo*-nucleolytic digestion. FAN1’s ‘inchworm’ pausing of slip-out excision is suited to minimize incremental expansions and modulating disease onset.

## Introduction

Elucidating the mechanism of ongoing somatic tandem repeat expansions in Huntington’s patient tissues is critical to understanding what drives disease onset and may be insightful as to how onset can be modulated. Expansions of CAG/CTG repeats are responsible for >15 neurodegenerative neuromuscular diseases including Huntington disease (HD), Huntington disease like 2 (HDL2), spinocerebellar ataxias (SCAs = SCA1, SCA2, SCA3, SCA7, and SCA17), spinal and bulbar muscular atrophy (SBMA), and myotonic dystrophy (DM1/CDM1). While, at least 10 CGG/CCG repeat expansion loci have been associated with fragile sites and intellectual disability (FRAXA/*FMR1*, FRAXE, FRAXF, FRA2A, FRA7A, FRA10A, FRA11A, FRA11B, FRA12A, and FRA16A). Ongoing increases in repeat lengths in affected tissues, thought to be driven by DNA repair proteins, correlates with disease age-of-onset. Soon after the discovery of repeat expansion as a cause of disease in 1991, it was found that ongoing somatic repeat expansions were occurring in affected tissues of HD, DM1, and fragile X syndrome (FXS), including the brain, muscle, heart, and other tissues (Telenius et al., 1994). Somatic expansions were tissue-specific and development age-specific (Castel et al., 2011; Pinto et al., 2020; Wöhrle et al., 1992). The progression of expansion could be considerable and appeared to drive disease onset and severity (Kennedy et al., 2003; Swami et al., 2009). Thus, understanding the mechanism of somatic repeat expansions is critical.

Early studies to understand the mechanism of somatic repeat expansions revealed that expansions occurred in what was termed “synchronous” manner, where the majority—if not all—of the mutant alleles in a cell population expanded together by gaining approximately the same incremental number of repeats over a given time. For example, a DM1 cell line with (CTG)216 increased by ∼7–10 repeats/30 days (Peterlin et al., 1996; Yang et al., 2003). Similar incremental repeat expansions have been reported for CAG, CTG, and CGG repeat expansions in HD, DM1, and fragile X patient cells, respectively (Goold et al., 2019; Kim et al., 2020; Lokanga et al., 2013; Peterlin et al., 1996; Yang et al., 2003). Evidence from post-mortem patient brains also supports incremental changes of 1 to 3 CTG/CAG units per mutation event (Pinto et al., 2020; Telenius et al., 1994). In HD mice, expansion rates are ∼3.5 CAG units/month/brain cell (Lee et al., 2010, 2011b; Nakamori et al., 2020) . “Inchworming” forward, each expansion mutation event predominantly gains one repeat unit/event, but can involve changes of 5–15 repeat units/event (Higham et al., 2012; Veitch et al., 2007; Wheeler et al., 2007). In patient brains, the accumulation over years, of these ongoing expansions can be dramatic (up to an additional 1,000 repeats) and exacerbate disease (Kennedy et al., 2003) (Swami et al., 2009). Mathematical modeling of somatic repeat expansions suggest that it is the net cumulation of ultra-frequent (daily) repeat tract mutation events (Higham and Monckton, 2013; Higham et al., 2012), but molecular evidence explaining the mechanism by which the incremental inchworm-like expansions arise has been elusive.

The concept that ongoing somatic repeat expansions is driving disease age of onset and progression has recently been supported by powerful HD human genetic data revealing modifiers of disease onset to be DNA repair genes — whose protein products had previously been demonstrated to modify somatic repeat instability in CAG and CGG expanded mouse models (Loupe et al., 2020; Zhao and Usdin, 2018). These genes were also revealed as age of onset modifiers of numerous spinocerebellar ataxias (SCAs) (Bettencourt et al., 2016; Ciosi et al., 2019). Among them, *FAN1* is the strongest modifier of age-of-disease onset for seven neurodegenerative CAG trinucleotide repeat expansion diseases, including HD, SCA1-3, SCA6, SCA7 and SCA17. Different FAN1 variants can delay or hasten disease onset by up to six years (Deshmukh et al., 2021). Non-coding disease onset-delaying rs35811129 (Lee et al., 2019) and coding onset-hastening *FAN1* variants (p.R377W and p.R507H, SNP rs151322829 and rs150393409) have been identified (Lee et al., 2019), where the former may lead to increased *FAN1* expression (Goold et al., 2019) and the latter may have unknown effects upon FAN1 functions. Functionally, FAN1 nuclease is an enigma. A recent review covered the genomic region in which the FAN1 gene resides, its protein domains, its variants, the varied disease associations of FAN1, the breadth of functions considered for the FAN1 protein, and its biochemical functions (Deshmukh et al., 2021).

*FAN1* has been implicated in numerous clinical presentations. Bona fide recessive nuclease-ablating *FAN1* mutations lead to karyomegalic interstitial nephritis (Deshmukh et al., 2021). *FAN1* is one of six genes in region of the human genome associated with 15q13.3 microdeletion/microduplication syndrome, where copy number variations of zero, one, two, three, or four copies encompassing *FAN1*, can lead to autism, schizophrenia, and epilepsy (Deshmukh et al., 2021). Attempts to discern which of the 15q13.3 genes is responsible, revealed genetic variations in *FAN1* to be associated with both autism and schizophrenia (Ionita-Laza et al., 2014), but a mechanistic link of FAN1 to these disorders was not obvious. Interestingly, the same FAN1 coding variants, p.R377W and p.R507H, identified as modifiers of HD, were identified as modifiers of autism and schizophrenia (Ionita-Laza et al., 2014). Recently, ASD was shown to be significantly associated with the expansion of various repeat motifs at ∼2500 loci (Trost et al., 2020). In particular, ASD has long been associated with CGG expansions in fragile X syndrome and other CGG/CCG expanded fragile sites (Deshmukh et al., 2021). A connection, if any, by which *FAN1* may be linked to autism, is lacking but may be connected through repeat instability.

FAN1 can suppress hyper repeat expansions. Most of the HD age-of-onset modifying variants occur in DNA repair genes that had previously been demonstrated to drive somatic expansions of the mutant CAG repeats (Lee et al., 2015, 2019). This supports the hypothesis that HD-modifying variants in DNA repair genes may modulate their own expression levels or their activity to modulate somatic repeat instability, which will then affect disease onset. In contrast to most of the HD-modifier genes, FAN1 had never been considered as a regulator of repeat instability. Recent cellular data suggest possible paths by which FAN1 may act on repeat instability. *FAN1* dosage seems important, as several of the disease-delaying non-coding FAN1 variants show increased *FAN1* expression levels in human brains, supporting the concept that increased FAN1 levels may increase its ability to suppress disease onset (Ciosi et al., 2019; Goold et al., 2019; Kim et al., 2020; Lee et al., 2015, 2019). HD patient cells, including those differentiated to post-mitotic medium spiny neurons, the vulnerable cell type in HD, showed incremental broadening of the CAG expansion sizes, an effect that was enhanced when FAN1 was knocked-down (Goold et al., 2019; Kim et al., 2020). For example, cells starting with (CAG)121 showed ongoing gains of 1 CAG repeat every 17.7 ± 1.1 days, which, in the absence of FAN1 increased to 1 CAG repeat every 9.1 ± 0.6 days (Goold et al., 2019). Similarly, completely knocking-out *FAN1* in HD patient-derived iPSC cells with (CAG)72 by CRISPR/Cas9, lead to gradual increases in the modal repeat size by ∼1-5 units over six months of cell culture (Kim et al., 2020). HD mice showed age-related somatic expansions of a (CAG)48 incurred as gains of ∼4-5 CAG repeats/8 months in a subset of striatal cells, whereas *Fan1*-deficient mice of the same age, incurred greater gains of ∼9-10 repeats/8 months; nearly twice as many over the same time (Loupe et al., 2020). These data suggest that FAN1 restrains CAG repeat expansion. Similarly, knocking-out *Fan1* in fragile X mice led to enhanced rates of incremental CGG repeat expansions in the brain (Zhao and Usdin, 2018). Both the cellular and mouse data suggest that FAN1 suppresses but does not ablate the incremental inchworm-like CAG and CGG expansions. The mechanism by which FAN1 may regulate repeat instability is unknown, as there have been no studies that directly assess FAN1 processing of repeat DNAs. Together, these findings posit the testable concept that increases or decreases of at least one of FAN1’s functions may suppress or enhance somatic repeat expansions, that then lead to delayed- or hastened-onset of HD, respectively. Understanding the biochemical functions of FAN1, in particular it’s nuclease activity, relative to repeat instability diseases is crucial towards learning how this potent modifier might exert its effects upon diseases.

Repeat expansions arise by aberrant or escaped repair of slipped-DNA structures that form by out-of-register annealing of complementary repeat strands during DNA repair, replication or transcription (Nakamori et al., 2011; Pearson, 2002). Slipped-DNAs occur at the expanded CAG/CTG repeat disease locus in DM1 patient tissues, where their levels directly correlated with the levels of somatic repeat expansions, providing support for the involvement of slipped-DNAs in expansion mutations (Axford et al., 2013). Further support for the involvement of slipped-DNA in instability is that observation that interruptions of the CAG tract purity with *CAA* motifs are associated with genetically stable repeats in HD, SCA2, and SCA17 repeats (Findlay Black et al., 2020; Wright et al., 2019, 2020). The purity of the *HTT* CAG tract is one of the strongest modifiers of disease onset (Ciosi et al., 2019; Lee et al., 2019; Wright et al., 2020). For example, CAA interruptions of the CAG tract are associated with HD onsets delayed by ∼3-6 years (Wright et al., 2020), whereas the pure CAG tracts are associated with hastened disease onset by ∼13-29 years (Wright et al., 2020). Data suggest that CAA interruptions protect the repeat from length variations where the interruptions inhibit the formation of slipped-DNAs, but alternative processing of interrupted slip-outs, is also possible (Goldberg et al., 1995; Marquis Gacy et al., 1995; Pearson et al., 1998a). *In vitro* slipped-DNAs are excellent models of mutagenic intermediates of instability, have been biophysically characterized (Axford et al., 2013; Lin et al., 2010; Nakamori et al., 2011; Panigrahi et al., 2010; Pearson, 2002; Pearson et al., 2005; Reddy et al., 2014; Schmidt and Pearson, 2016; Tomé et al., 2013), and are similar to slipped-DNAs in patient tissues (Axford et al., 2013). Targeting slipped-DNA with slip-out specific ligands can induce repeat contractions of the mutant tract in brains of HD mice, supporting the involvement of slipped-DNAs in instability, as well as the utility of these as slip-out models (Nakamori et al., 2020). How slipped-DNAs are processed and understanding the effect of instability-modfying interrupting motifs and ligands, must involve a DNA nuclease.

A DNA nuclease must be involved in the process of disease-associated repeat expansions, contractions, and maintenance, presumably to process the slipped-DNAs that form during mutagenic length changes. However, it is unclear which is the key nuclease involved in repeat instability, as recently reviewed (Mitra et al., 2021; Trost et al., 2020; York and Lat, 2021). The emerging human genetic evidence implicating FAN1 as the strongest modifier of disease onset for seven CAG/polyQ diseases (Bettencourt et al., 2016; Ciosi et al., 2019; Lee et al., 2015; Mergener et al., 2020), highlights the need to test its nuclease activity on repeats. FAN1’s nuclease activity, characterized for its involvement in DNA interstrand crosslink repair, recovery of stalled replication forks, and homologous recombination, has not been assessed for its ability to act upon disease-associated repeats or the unusual structures they can form. Based upon the recent association of ASD and SCZ with repeat expansions (Trost et al., 2020), the FAN1- 15q13.3-ASD/SCZ association, and the association of FAN1 variants and age of onset for HD and SCAs (Mitra et al., 2021; Trost et al., 2020; York and Lat, 2021), a shared mechanistic connection of FAN1 to repeat expansions was recently hypothesized (Deshmukh et al., 2021). Here we tested the effects of *FAN1* variants (protein levels, WT, p.R507H, p.R377W, nuclease-dead mutants p.D960A and p.D981A-R982A) on slipped-CAG/CTG DNAs – mutagenic intermediates of instability, shown to form in patient tissues (Axford et al., 2013).

## Results

### FAN1 and its variants bind CAG and CTG slipped-DNAs

To assess the interaction of FAN1 with slipped-DNAs, we purified from baculovirus-infected insect cells recombinant forms of human FAN1 differing only by the presence of variants (WT = FAN1^p.WT^, p.R377W = FAN1^p.R377W^, p.R507H = p.FAN1^p.R507H^), or mutations demonstrated to perturb FAN1-self dimerization conformation leading to catalytically unproductive DNA binding (p.D981A-R982A = FAN1^p.D981A-R982A^) (MacKay et al., 2010; Zhao et al., 2014) or nuclease activity (FAN1^p.D960A^) (Zhao et al., 2014). The locations of the disease-linked variants are shown relative to characterized functional domains of FAN1 and protein preparations having a single electrophoretic species are shown in Figures 1A and 1B.

**Figure 1:**
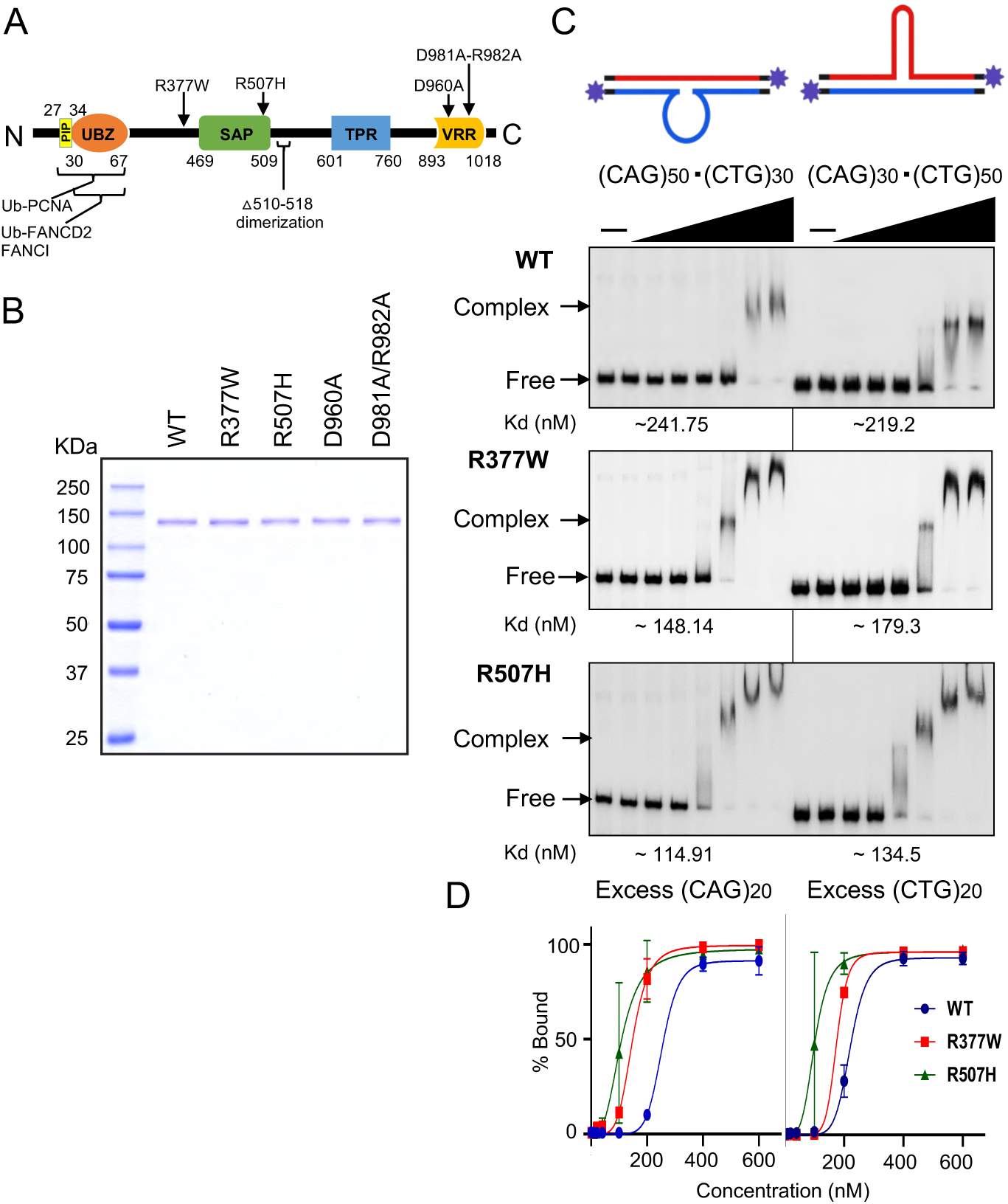
**FAN1 and its variants binds CAG and CTG slipped-DNAs**: (A) FAN1 protein domain map showing locations of variants (R377W & R507H), mutants (D960A & D981A-R982A), dimerization regions, SAP, TPR, and VRR_NUC (nuclease) domains; functions of various FAN1 domains are reviewed in (Deshmukh et al., 2021). (B) Representative SDS-PAGE analysis of various baculovirus expressed and purified full-length human FAN1 protein forms (C) Representative DNA-binding (mobility-shift) assay of FAN1^p.WT^, FAN1^p.R377W^, and FAN1^p.R507H^ on excess CAG/CTG slip-out DNA. Schematic of DNA substrates is shown. Concentration-dependent DNA-binding with 10-600 nM of FAN1 (0,10, 20, 40, 100, 200, 400 and 600 nM) on 50 nM slipped-DNAs (3′-end filled with ^32^P-*α*dNTPs, purple star). Samples were resolved on 4% PAGE for 1 hr 30 min at room temperature, gels dried and exposed to PhosphorImager and observed under Typhoon-FLA8000. (D) Data were analyzed by ImageQuant and plotted on GraphPad Prism 8.2 (representative gels of n=3 independent replicates). See also **Fig. S1**.

We examined FAN1 binding to various structurally characterized forms of disease-relevant lengths of CAG/CTG containing DNAs (Axford et al., 2013; Lin et al., 2010; Nakamori et al., 2011; Panigrahi et al., 2010; Pearson, 2002; Pearson et al., 2005; Reddy et al., 2014; Schmidt and Pearson, 2016; Tomé et al., 2013). These include fully-paired duplex expanded repeat strands, (CAG)50•(CTG)50; control repeat-free duplex DNA (Figure S1A top); and slipped-DNAs having an excess of Δ(CAG)20 = [(CAG)50•(CTG)30]; or an excess of Δ(CTG)20 = [(CAG)30•(CTG)50] (Figure 1C top) (Pearson and Sinden, 1996; Pearson et al., 1997, 1998c, 1998b).

FAN1 DNA binding efficiency was determined directly in a concentration-dependent manner by electrophoretic mobility shift assay. FAN1^p.WT^ did not bind to fully-duplexed DNAs with or without CAG/CTG repeats (Figure S1A). FAN1^p.WT^ bound to slipped-DNAs with an excess of Δ(CAG)20 or Δ(CTG)20, with dissociation constants (Kd) of 241.75 and 219.2 nM, respectively, indicating structure-specific recognition of slipped-DNAs (Figure 1C, quantifications are in Figure 1D, and summarized in Table1). The FAN1^p.R377W^ and FAN1^p.R507H^ variants being proximal to the conserved DNA binding residue Y374 (Wang et al., 2014) and within the SAP DNA-binding domain, respectively, warranted testing of their effect on DNA binding. Both FAN1^p.R507H^ and FAN1^p.R377W^ showed mildly increased binding affinities, up to two-fold, to Δ(CAG)20 and Δ(CTG)20 slip-outs compared to FAN1^p.WT^, but the differences were not statistically significant (Figure 1C and D). Thus, FAN1 specifically recognizes CAG/CTG slip-outs, and both HD disease-hastening variants interacted with slipped-DNAs greater than FAN1^p.WT^ (Figure 1C and 1D). Each FAN1 form yields multiple shifted complexes with increasing protein concentration. The nuclease-dead FAN1^p.D960A^ showed DNA binding affinity comparable to FAN1^p.WT^ (Figure S1, Kd = 210 nM). However, FAN1^p.D981A-R982A^, where D981 is involved in DNA binding and dimerization of FAN1 protein (Yan et al., 2015; Zhao et al., 2014), shows minimal binding under matched conditions (Figure S1B bottom gel). It is important to note that the FAN1^p.D960A^, but not the FAN1^p.D981A-R982A^, retains slipped-DNA binding capacity, thereby permitting the former to be used as a loss of nuclease, but not DNA binding. Clearly, DNA binding capacity of FAN1 mutants does not always reflect its nuclease activity. We conclude that FAN1 specifically recognizes slipped-CAG/CTG DNAs, with binding affinities of FAN1^p.R507H^ > FAN1^p.R377W^ > FAN1^p.WT^ = FAN1^p.D960A^ >>> FAN1^p.D981A-R982A^. The Kd ranges for WT FAN1 we observe are similar to previous reports for repeat-free DNA substrates (Wang et al., 2014).

### FAN1 binds flap-bourne and duplex-bourne slip-out DNAs

FAN1 has been characterized to bind to flap-DNAs, that model replication forks, open transcription bubbles, or active repair (Kratz et al., 2010; MacKay et al., 2010; Wang et al., 2014; Zhao et al., 2014). Repeats can form slipped-DNAs during replication, transcription, and repair (Axford et al., 2013; Lin et al., 2010; Nakamori et al., 2011; Panigrahi et al., 2010; Pearson, 2002; Pearson et al., 2005; Reddy et al., 2014; Schmidt and Pearson, 2016; Slean et al., 2013; Tomé et al., 2013). Slip-outs can arise in the flap or duplex region, and can be composed of CAG or CTG repeats, where the former has more random-coil character, but both can assume intrastrand hairpin-like conformations (Axford et al., 2013; Lin et al., 2010; Nakamori et al., 2011; Panigrahi et al., 2010; Pearson, 2002; Pearson et al., 2005; Reddy et al., 2014; Schmidt and Pearson, 2016; Slean et al., 2013; Tomé et al., 2013). These Flap-bourne and Duplex-bourne slip-outs are referred to as Flap-(CAG)20, Flap-(CTG)20, Duplex-(CAG)20, and Duplex-(CTG)20, respectively (Figure S2 top).

As controls, we assessed binding of FAN1 to unstructured long and short flap-DNAs, USL and USD, having similar lengths of single-stranded flaps as the Flap-bourne and Duplex-bourne slip-outs, respectively (Figures S2). The HD-linked FAN1 variants have only mild differences in binding affinity for the Flap-bourne and Duplex-bourne slip-outs relative to wildtype FAN1, but the differences were not statistically significant (Figure S2, see Table1). We conclude that FAN1 and its variants can recognize repeat slip-out DNAs in various contexts.

### FAN1 and its variants can dimerize on slipped-DNAs

Two FAN1 molecules act as a dimeric complex to process DNA structures. FAN1 alone is monomeric (Pennell et al., 2014), but DNA binding promotes “head to tail” FAN1 dimerization (Jin and Cho, 2017; Zhao et al., 2014). Three distinct FAN1 dimer–DNA conformations can arise, each involving different DNA-contact interfaces (Jin and Cho, 2017; Rao et al., 2018; Zhao et al., 2014). DNA binding and *endo*-nucleolytic cleavage preference can differ markedly between monomeric and dimeric forms of FAN1 (Jin and Cho, 2017). Dimerization is not required for FAN1’s *exo*-nucleolytic function (Zhao et al., 2014). FAN1 mutants defective in DNA binding can affect dimerization, and mutants that affect dimerization can affect DNA binding or cleavage modes (Jin and Cho, 2017). For example, deletion of residues 510-518 abolished FAN1 dimerization (Rao et al., 2018; Wang et al., 2014), modified the DNA binding mode and diminished the ability to cleave long (15-40 nucleotides), but not short (1-5 nucleotide) DNA flaps (Jin and Cho, 2017; Rao et al., 2018). Similarly, the triple mutant K525E/R526E/K528E showed greatly diminished in DNA-induced dimerization, undetectable DNA binding by band-shift, undetectable *endo*-nuclease activity, but *exo*-nuclease activity similar to wildtype FAN1 (Zhao et al., 2014).

Since FAN1 dimers differentially process long versus short flaps (Rao et al., 2018), we tested whether FAN1 can dimerize on DNAs with long slipped-outs of 20 repeats (60 nucleotides). Moreover, since the HD variant FAN1^p.R507H^ is proximal to the residues required for dimerization (residues p.510-518) (Figure 1A), we monitored its ability to dimerize on slip-outs of 20 excess repeats. All FAN1 forms showed similar abilities to dimerize on slipped-DNA (Figure S3). That the FAN1^p.R507H^ and FAN1^p.R377W^ variants were able to dimerize upon DNA binding with comparable abilities to FAN1^p.WT^ is consistent with their having similar DNA binding affinities. Their ability to dimerize is also consistent with their ability to form multiple electrophoretically shifted FAN1-DNA complexes (Figures 1, S1, & S2). That none of the variants affected DNA binding is consistent with their being distal to any of the DNA binding interfaces of any of the bound forms of FAN1 (Zhao et al., 2014).

### FAN1 *endo*-nucleolytic cleaves slipped-DNAs similar to repeat-free DNA flaps

FAN1 has *endo*- and *exo*-nuclease activities, incising internal sites of DNA and hydrolyzing nucleotides from the 5′→3′ end of a DNA flap, respectively. *Endo*- and *exo-* nucleolytic activities can be distinguished by labeling the 5′ or 3′ ends of the DNA strand, respectively. We examined the *endo*-nuclease activity of FAN1 and its variants on Flap- (CAG)20, and Flap-(CTG)20 DNAs, and repeat-free unstructured long flap (USL) (Figure S4). All flap DNAs showed exactly the same *endo*-nucleolytic cleavage patterns, by all FAN1 forms. This pattern reflected the *endo*-nucleolytic cleavage patterns of all previous FAN1 studies, each using various repeat-free flap-DNAs, having different sequences (Kratz et al., 2010; MacKay et al., 2010; Pennell et al., 2014; Smogorzewska et al., 2010; Wang et al., 2014; Yoshikiyo et al., 2010; Zhao et al., 2014). Our FAN1 preparations were free of contaminating nucleases, as multiple preparations of the nuclease-dead FAN1^p.D960A^, deficient in *endo*- and *exo-*nucleolytic activities (Smogorzewska et al., 2010; Zhao et al., 2014), were, as expected nuclease-free (Figure S4, last lane of most gels, see also Methods). The rate of *endo*-nucleolytic cleavage, calculated as previously described (MacKay et al., 2010), did not vary significantly between FAN1 forms (Table S2). As with previous studies, only the flap DNA strand showed predominant cleavage by FAN1 (not shown). Thus, neither the presence nor absence of a slip-out, or of the HD variants, altered FAN1’s *endo*-nucleolytic activities.

### FAN1 *exo*-nucleolytic cleavage sites depend on slip-out sequence and location: *Exo*-nucleolytic pausing in Flap-bourne slip-outs

Unusual DNA conformations, like mismatches or unwound bubbles, and specific sequences, can be hurdles to the progression of lambda *exo*-nuclease (Lee et al., 2011a; Perkins et al., 2003). To assess whether slipped-DNA structures may impede FAN1’s *exo*-nuclease, we used a high-resolution electrophoretic gel-method, developed by Perkins *et al*., to map at nucleotide levels the locations of *exo*-nuclease pauses (Perkins et al., 2003). Electrophoretic bands are DNA cleavage products, and *exo*-nucleolytic pauses are characterized by darker bands, where band intensity is related to the pause strength, which is the product of the duration and probability of pausing. Intense bands are stronger sustained *exo*-nucleolytic pause sites.

Striking cleavage site preferences and digestion rate differences were evident for Flap- (CAG)20 versus Flap-(CTG)20, versus repeat-free flaps, in time-dependent FAN1 *exo*- nucleolytic digestions (Figure 2A). Digestion of CAG slip-outs was slower than CTG slip-outs. Cleavage occurred throughout the repeat tracts, while the repeat-free flap was devoid of cleavage. Flap-(CAG)20 was cleaved preferentially at every 1^st^ and 2^nd^ nucleotide throughout the length of the CAG slip-out, cleaving between the C-A and between the A-G, 5′-C↓A↓GC↓A↓G-3′ (Figure 2A, middle panel, strand regions 2-3, schematically indicated). The darker bands at every cleavage site throughout the repeat are the *exo*-nucleolytic pauses. Cleavage progresses over time from the 5′ end of the CAG slip-out. A striking set of accumulated digestion products were evident at the 3′ end of the CAG tract (Figure 2A, middle panel region 3, arrow heads). These intense accumulated *exo*-nucleolytic pauses are referred hereafter as “hotspots”. As per Perkins *et al*., *exo*-nucleolytic pauses were mapped using chemical sequencing size markers (Perkins et al., 2003) and found to localize within the last three CAG units of the CAG tract. Single-molecule data cannot reveal the location of nuclease pause sites (Perkins et al., 2003). Digestion did eventually proceed beyond the localized pauses (Figure 2A, lower portion of gels). The cleavage rate of each region (1-5) of each substrate is quantified in Figure S5. The Flap-(CTG)20 digestion pattern is strikingly distinct from the Flap-(CAG)20 cleavage. Under these matched conditions, scission is prominent at multiple sites along only the 3′ strand of the slip-out, revealing a set of nuclease pauses (Figure 2A, top of rightmost panel, region 3). Increasing concentrations of Flap-(CTG)20 substrate revealed scissile sites throughout the repeat tract, occurring between each nucleotide of the repeat, 5′-C↓T↓G↓C↓T↓G-3′ (Figure 2A, right most panel, rightmost lane). The cleavage rate of each region (1-5) of each substrate in an extended time-course is quantified in Figure S5. Rate of CAG slip-out digestion was significantly slower than CTG, which was comparable to repeat-free flaps (162.32 ± 34.94, 75.95 ± 43.01 and 142.37 ± 47.62 nucleotides/min, respectively, see extended time-course Figure S6, summarized in Table S2). Flap-(CAG)20 is digested ∼2 fold slower than Flap-(CTG)20. The rates of cleavage of the repeat-free and slipped-CTG were comparable to previously reported FAN1 digestion rates of other repeat-free DNAs (MacKay et al., 2010; Wang et al., 2014), while the slipped-CAG was considerably slower (p<0.0001, Welch’s *t* test). In contrast, the repeat-free flap, incurred a single predominant cleavage in the flap, (at ∼25 nucleotides into the 74 nt flap) and a series of cuts in the duplex region ∼4-5 nucleotides beyond the ssDNA-dsDNA junction (Figure 2A, regions 2 and 5 of leftmost panel). All flap-DNAs, with or without slip-outs, showed a similar cleavage pattern in the duplex region beyond the flaps/slip-outs (Figure 2A, region 5). This pattern reflected the *exo*- nucleolytic cleavage patterns of all previous FAN1 studies, each using various repeat-free flap-DNAs of varying sequences (Kratz et al., 2010; MacKay et al., 2010; Pennell et al., 2014; Smogorzewska et al., 2010; Wang et al., 2014; Yoshikiyo et al., 2010; Zhao et al., 2014). We conclude that FAN1 cleavage patterns of flap-bourne slip-outs are determined by slip-out sequence, CAG versus CTG. Cleavage occurs at nearly every nucleotide of the repeat tract only, distinct from the previously reported successive cleavage at every third nucleotide on non-CAG/CTG flaps (Wang et al., 2014). Nearly every cleavage event in the repeat tracts is evident as an *exo*-nucleolytic pause, with the most intense pausing occurring at the 3′ ends of the CAG and CTG slip-outs. Pause site hotspots are summarized in Figure S8. Under similar conditions the unstructured repeat-free flaps do not show cleavages. throughout the flap. Moreover, the rate of FAN1 digestion is significantly slower for CAG than CTG slip-outs. Presumably, the structural differences between CAG and CTG slip-outs account for their differential cleavage rates and patterns (Axford et al., 2013; Cleary et al., 2010; Lin et al., 2010; Nakamori et al., 2020; Pearson et al., 2005).

**Figure 2:**
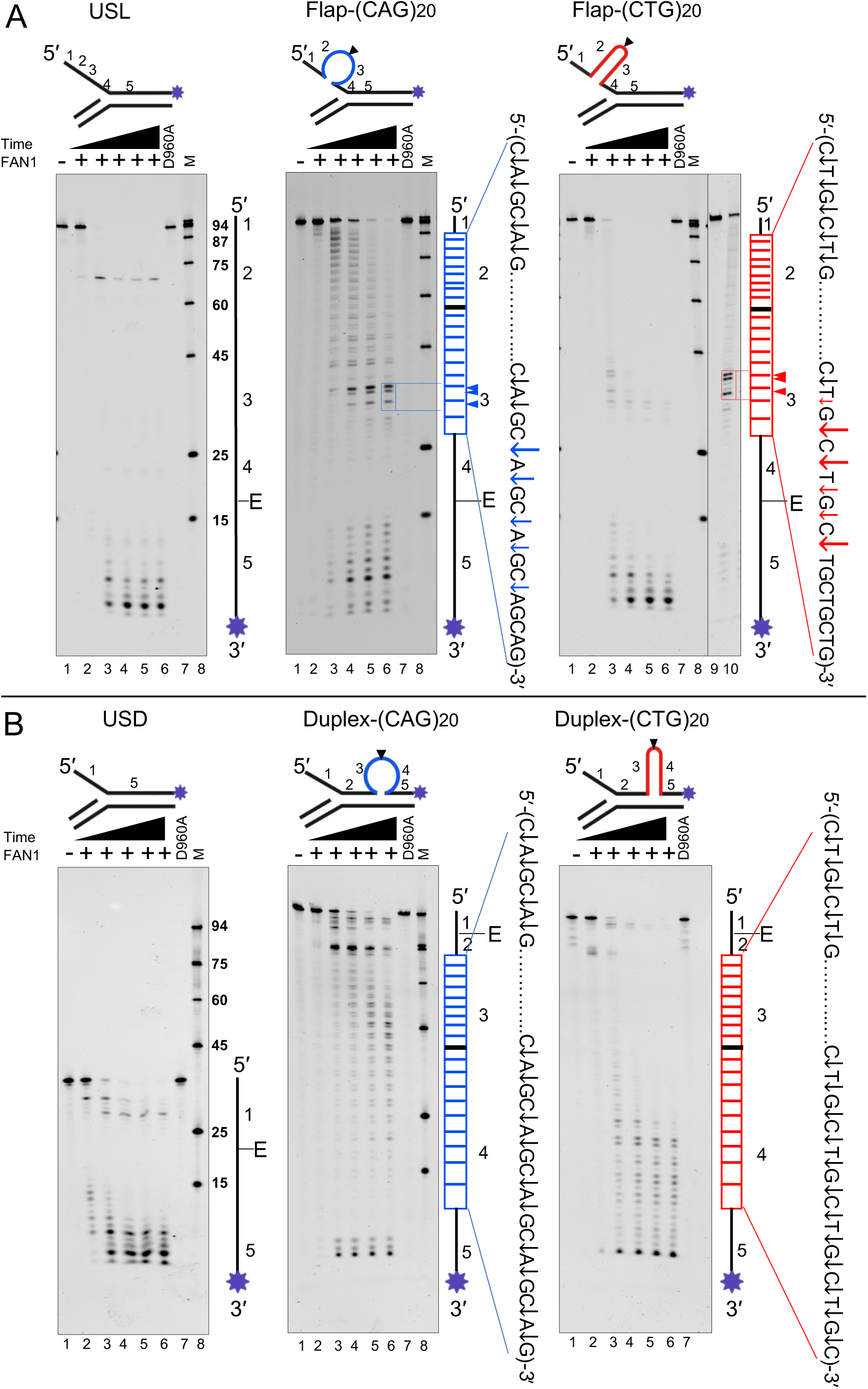
**FAN1 *exo*-nucleolytic cleavage / pause sites depend on slip-out sequence and location:** (A) USL (unstructured long flap), Flap-(CAG)20, and Flap– (CTG)20 DNA substrates (B) USD, Duplex-(CAG)20 and Duplex-(CTG)20 substrates. All DNAs have 3′-FAM labelled on flap strand (purple star), schematics are shown above each denaturing gel, and labeled strand schematic indicated along the length of the gel, with DNA regions numbered, and repeat tract and its center (black line) indicated, “E” denotes elbow region between ds DNA to ssDNA junction. Following digestions products were denatured and resolved on denaturing sequencing gels. A. Lane 1 indicates, substrate alone and lanes 2-6 show time-dependent digests (0, 5, 10, 15 and 20 min) at 200 nM of human FAN1 and 100 nM DNA substrate (protein: DNA (2:1)). Lane 9 and 10 in rightmost panel of panel A shows substrate alone and nuclease activity of FAN1 at reduced protein concentration (protein: DNA 1:2) for 10 minutes respectively, permitting identification in lane 10 of individual FAN1 cleavage/pause sites on Flap-(CTG)20 DNA substrate. Lane 7 shows absence of nuclease activity with 200 nM FAN1^p.D960A^ for 20 min., indicating that our FAN1 preparations were free of contaminating nucleases, as multiple preparations of the nuclease-dead FAN1^p.D960A^, known to be deficient in *endo*- and *exo-*nucleolytic activities (Smogorzewska et al., 2010; Zhao et al., 2014), were, as expected, consistently nuclease-free. Lane 8 shows size marker. *Exo*-nuclease scissile sites/pause sites were mapped using Maxam-Gilbert chemical DNA sequencing. For quantification of digestions in each DNA region see Figure S5. Representative gels for n=3 independent replicates.

As noted above rates of digestion varied by location within the repeat tract, beginning to slow at the beginning of the repeat, middle of the repeat, and end of the repeat, with excision rates that are 1-, 9-, and 28-fold slower, respectively (Figure S6 & Table S2). Nuclease digestion rates were determined, where reactions were performed with identical protein : DNA ratios, over time (Iyer et al., 2006; MacKay et al., 2010; Subramanian et al., 2003; Wang et al., 2014). Michaelis-Menten type kinetics, where the enzyme is fully saturated by DNA substrate, and rates calculated on end point digestion, have been reported for rad27/FEN1 on slipped-CAG DNAs (Liu and Bambara, 2003). However, unlike rad27/FEN1 which does not cleave within the repeat tract (Liu and Bambara, 2003), FAN1 yields numerous cleavages throughout the repeat tract as well as accumulated intense paused cleavage products, supporting differential digestion rates along the DNA, making such Michaelis-Menten calculations difficult, if not impossible to interpret for the FAN1-repeat digestions (not shown).

### FAN1 *exo*-nucleolytic cleavage throughout Duplex-bourne slip-outs

Repeat slip-outs extruding from the duplex region adjacent to a 5′-single-stranded flap, as arise upstream of replication forks, transcription bubbles, or sites of damage repair, were digested by FAN1. Cleavage patterns were assessed for duplex-bourne slip-outs, Duplex-(CAG)20 and Duplex-(CTG)20 (Figure 2B). Again, CAG slip-outs were digested significantly slower than CTG slip-outs (Figure S6 & Table S2). Slip-out regions showed distinct cleavage patterns for CAG versus CTG slip-outs (regions 3 and 4, Figure 2B, middle and rightmost). Cleavage patterns of the Duplex-bourne slip-outs paralleled that of their Flap-bourne counterparts. FAN1 cleaved preferentially along the CAG slip-out; 5′-C↓A↓GC↓A↓G-3′ and the CTG slip-out; 5′-C↓T↓G↓C↓T↓G-3′ (Figure 2B). However, nuclease pauses were not evident in the Duplex-bourne slip-outs. The cleavage pattern in the single-stranded flap and the duplex region beyond the slip-outs was similar for all substrates (cutting predominantly regions 1 & 5) (Figure 2B, for regional cleavage rate for each substrate see Figure S5). We conclude that FAN1 cleavage rates and patterns are determined by slip-out sequence, CAG versus CTG, and location of the slip-out, where Flap-bourne but not Duplex-bourne slip-outs accumulate paused cleavage products.

### FAN1 processes slip-outs in a dose-dependent manner

Considering disease-delaying non-coding *FAN1* variants are expected to give rise to the over-expression of FAN1 protein (Goold et al., 2019; Kim et al., 2020), we performed controlled concentration-dependent nuclease assays (Figure S7B). Cleavage activity is directly proportional to FAN1 concentration. Qualitatively and quantitatively, FAN1^p.R377W^, FAN1^p.R507H^ and FAN1^p.WT^ show similar cleavage patterns on the slipped-DNAs (Figure S7A and S7B). Both Flap-(CAG)20 and Flap-(CTG)20 showed similar cleavage pauses for all FAN1 forms. However, FAN1^p.R377W^ and FAN1^p.R507H^, showed specific *exo*- nuclease pauses on Duplex-(CTG)20, that were not produced by FAN1^p.WT^ (Figure S7B rightmost panel). The nuclease activity of all FAN1 forms, increased in a dosage-dependent manner, consistent with the report that *FAN1* over-expression of the onset-delaying variants could clinically override the onset-hastening FAN1 coding variants, when both are present in the same HD individual (Kim et al., 2020). We further analyzed rate of digestion for FAN1 variants on Flap-bourne and Duplex bourne slip-out DNA, Variants showed mild, non-significant differences in digestion rates (Figure S7C summarized Table S2).

### FAN1 excises distributively and pauses in the repeat

The in-homogenous cleavage patterns throughout CAG and CTG repeat tracks of the slipped-DNAs (Figure 2) warranted further investigation. The production of ladder band digestion products along the repeat tract by FAN1 suggested a distributive cleavage process, where iterative cycles of enzyme binding, scission, and dissociation lead to DNA degradation. This contrasts to processive *exo*-nucleases where the enzyme remains associated with the DNA during sequential removal of nucleotides. FAN1 acts distributively on slipped-DNAs, as flooding reactions with unlabeled competitor DNA following the initiation of digestion, slowed nuclease progression (Figure 3). Were digestion processive, involving a single binding event through to complete digestion, such flooding with competitor DNA would have no effect on the digestion rates.

**Figure 3:**
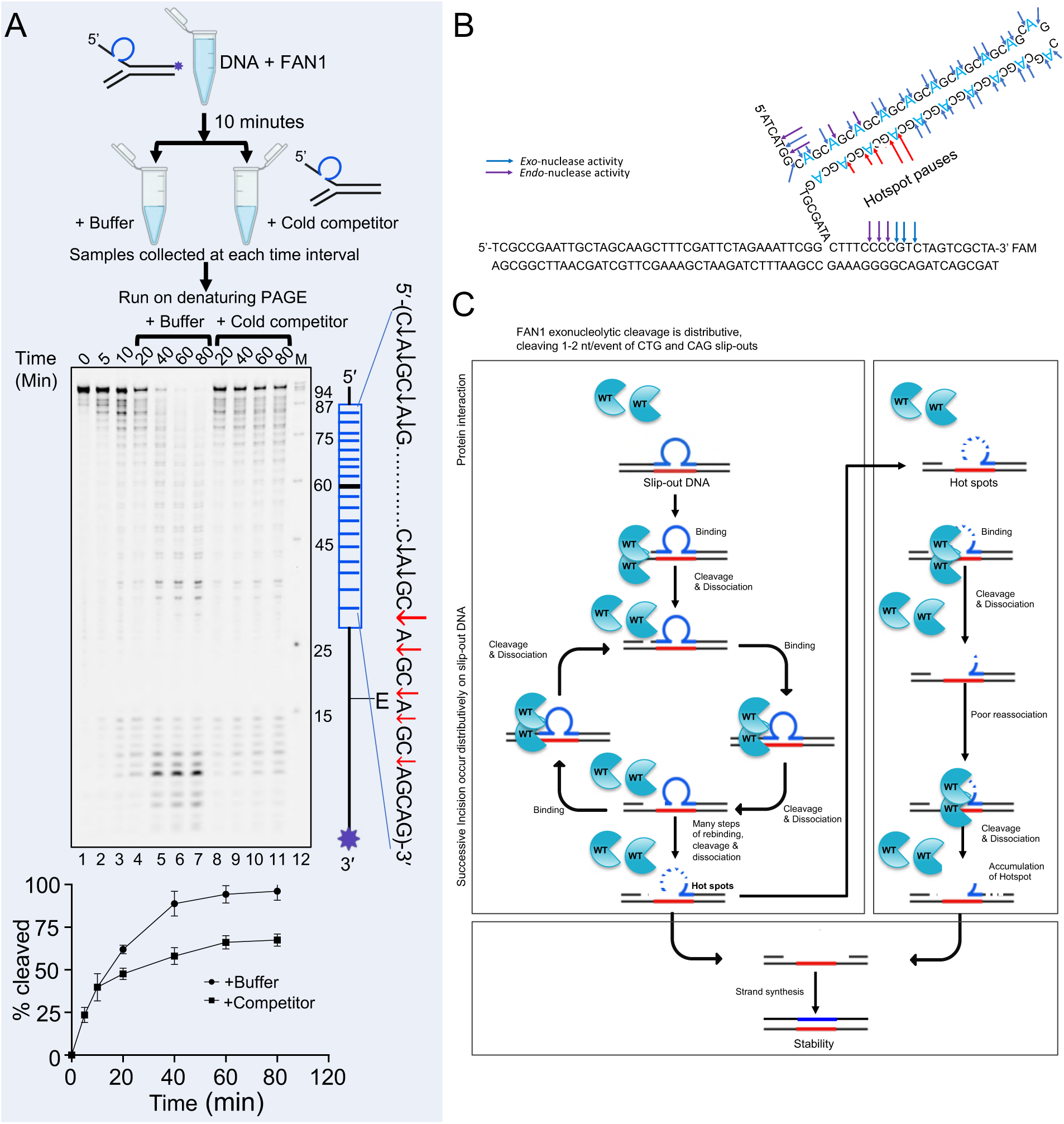
**FAN1 *exo*-nucleolytically digests distributively and pauses in the repeat: A)** Processivity versus distributivity test. 3′-FAM labelled slipped-DNA (purple star), schematics and labeled strand schematic indicated along the length of the gel, and repeat tract and its center (black line) indicated, “E” denotes elbow region between ds DNA to ssDNA junction. Time course of *exo*-nuclease reaction (50 nM FAN1 and 500 nM 3′-FAM labelled Flap-(CAG)20), which was split at the 10 minute timepoint and challenged with either buffer (lanes 4-7) or an excess (2000 nM) of cold (unlabeled) competitor substrate (lanes 8-11). Representative gel of n=3 replicates. Percentage of cleavage products for each split reaction were graphed, as shown below the gel. At 20 minutes, in the presence of competitor all product levels were essentially unchanged, whereas in the mock-buffer treated reaction digestions continued, indicating that successive cleavage events do not occur processively, but are highly distributive, with cycles of FAN1 binding, cleaving, pausing, and dissociating. **B)** Cleavage / pause sites were shown in schematic. **C)** Proposed model of FAN1 processing slip-out DNA, where each successive binding, cleavage, pause, and dissociation steps lead to accumulated DNA digestion products, to particularly enriched toward the 3’ end at hotspot pause sites, to which FAN1 has poor reassociation and cleaves in duplex DNA and removes slip-out. This gapped DNA may potentially act as substrate for DNA polymerases or ligases to expand or fix the repeat length. See also Fig. S9.

The accumulation of FAN1 paused cleavage products suggests that either the flap length and/or alterations of the physical form of DNA, as it is digested, may affect progression of FAN1 digestion. Paused digestion may occur due to alteration of at least one step of the digestion process. Considering FAN1 acts distributively, accumulated paused products could be poor binding to the hotspot DNA (reassociation), poor cleavage of the hotspot, or poor dissociation from the cleavage product. Experiments support the reduced binding of FAN1 to the accumulated nuclease pause products (Figure S8A), likely due to altered DNA conformation, going from intra-strand slip-out to single-stranded tracts of only <3 repeat units, too short for intra-strand base-pairing. We further examined *exo*- nucleolytic pausing hotspot observed in Duplex-(CTG)20 (Figure S7B rightmost panel) for DNA binding and nuclease activity. FAN1 and its forms show significantly reduced DNA binding to hotspot DNA substrate (p<0.0001, Welch’s *t* test) (Figure S9A). Suggesting hotspot DNA is poor substrate for FAN1 and its forms. To test the length dependency, we analyzed hotspot + 1nt, hotspot and hotspot - 1nt DNAs. Interestingly, clear difference of cleavage pauses was observed with hotspot + 1nt and hotspot - 1nt nucleotide (Figure S9B). It is notable that all FAN1 *exo*-nucleolytic pauses arose in similar locations, near the terminal 3-4 repeat units of the repeat tract, supporting a length-structure dependency.

### Fully-paired hairpins arrest FAN1, while A•A and T•T mismatches permit slip-out excision

The presence of a slip-out clearly affects FAN1 cleavage pattern and rate, which we have demonstrated can vary between CAG and CTG slip-outs. CAG and CTG slip-outs assume intra-strand hairpins with A•A and T•T mismatches, respectively (Marquis Gacy et al., 1995). It is unknown whether the hairpin-like structure of CAG and CTG slip-outs could be a determining factor in FAN1 cleavage patterns or rates. Nor is it known if the A•A and T•T mismatches contribute to the cleavage patterns of the (CAG)20 and (CTG)20 slip-outs. Perfect repeat-free stem-loops may be processed differently by FAN1 compared to CAG or CTG slip-out hairpins. The diversity of DNA conformation is limited for a hairpin tip that is locked at a fixed point, as would occur at perfect palindromic inverted repeats, where the hairpin tip is anchored by the center of inversion. In contrast, the hairpin tip of a slipped repeat tract that can constantly shift the location of the hairpin, a dynamic that can change as FAN1 continually degrades the repeat. CAG hairpins with recessed 5′ ends, similar to the FAN1 substrates used herein, can rapidly and spontaneously, undergo slippage thereby shifting the location of the hairpin tip (Xu et al., 2020).

We assessed FAN1 cleavage of a fully-paired stem-loop, devoid of repeats, with similar number of nucleotides as the CAG/CTG slip-outs (Flap-IS). This repeat-free stem-loop was also devoid of A•A or T•T mismatches. The cleavage pattern of the repeat-free hairpin, under identical conditions as the slipped-DNAs, showed incomplete digestion, with scission occurring predominantly at the 5′ base of the “anchored” hairpin tip (Figure 4A).

**Figure 4:**
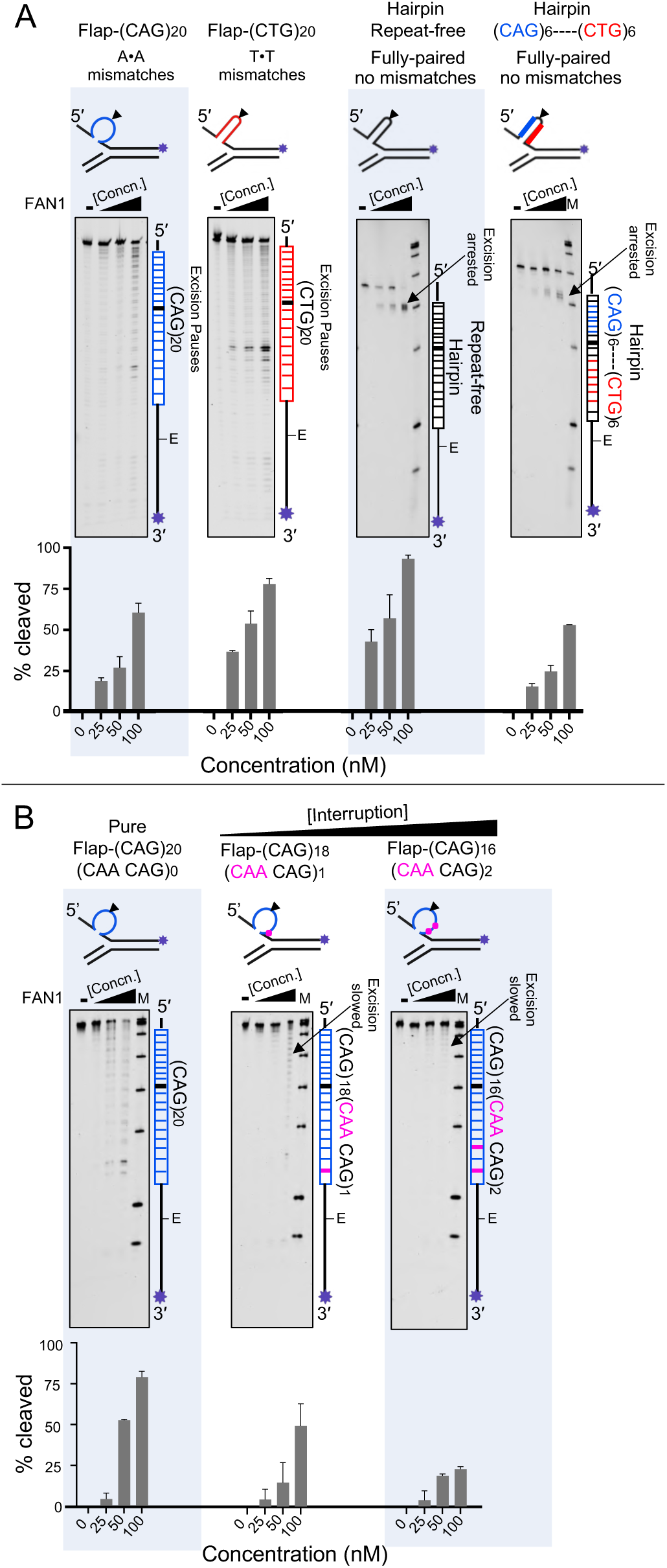
**FAN1, *exo*-nucleolytic pauses, arrests, A•A/T•T mismatches, & CAA interruptions: A)** Flap-(CAG)20, Flap- (CTG)20, repeat-free fully-paired hairpin and fully-paired (CAG)6---(CTG)6 hairpin DNA 3′ FAM labeled on flap strand (purple star in schematic above and alongside each gel). 100 nM of each DNAs was incubated with 25, 50 and 100 nM of FAN1^p.wt^ for 20 minutes reactions stopped by adding 95% formamide, and products resolved on 6% denaturing PAGE. Cleavage was quantified by ImageQuant and plotted by GraphPad Prism 8.2 Representative gels of n=3 replicates. Excision pauses are evident throughout the repeat, while excision arrests are prior to repeat, as indicated with arrows. **B)** Pure CAG tracts and interrupted tracts (CAA interruptions = violet dots/bars in schematic above and alongside each gel, with 3′ FAM label = purple star) were incubated with 100, 200, and 300 nM of FAN1^p.WT^ for 20 minutes, reactions were stopped with 95% formamide and samples were resolved on 6% denaturing PAGE and analyzed by ImageQuant and plotted by GraphPad Prism 8.2. Experiments were done in triplicate, shown are representative gels. Error bars represent ±SD. Excision pauses are evident throughout the Pure repeat, while pauses are enriched at the 5’ end of CAA-interrupted repeat tracts due to slowed excision, as indicated with arrows.

The intensity of cleaved products persisted at the hairpin base, with little evidence of further excision into or past the repeat-free hairpin (Figure 4A). Thus, the stem and tip of the fully-paired repeat-free hairpin was protected from any scission, a pattern distinct from either CAG or CTG slip-outs, essentially *arresting* FAN1 excision. Unlike the CAG and CTG slip-outs, the repeat-free hairpin was not completely digested, and did not incur successive, homogenous cleavages or *pauses* throughout, where the hairpin tip was recalcitrant to cleavage by FAN1. Moreover, subsequent excision in regions past the hairpin were also protected, suggesting that the fully-paired hairpin *arrested* excision by FAN1. The poor ability of FAN1 to cleave the anchored hairpin tip is consistent with ssDNA being one of FAN1’s worst substrates (Kratz et al., 2010; Smogorzewska et al., 2010). In contrast, distinct scissions at the hairpin tips by the single-strand specific mung bean *endo*-nuclease occur on numerous repeat-free hairpins of differing sequences (Kabotyanski et al., 1995). Similarly, the CAG and CTG slip-outs, when digested with mung bean *endo*-nuclease digestion show a single slip-out with distinct hairpin tips, that are sensitive to mung bean nuclease (Nakamori et al., 2020; Pearson, 2002). However, the location of these tips may alter with the successive shortenings of the repeat by FAN1 digestions.

Repeat-free stem loops have anchored hairpin tips, with sensitivity at the hairpin tips for numerous nucleases, regardless of stem-loop sequence (Kabotyanski et al., 1995). The base of a repeat-free stem loop, the three-way junction from which the hairpin extrudes can also serve as a substrate for numerous nucleases. Similarly, CAG or CTG repeat hairpins will extrude as a single hairpin, where the location of the hairpin tip is defined by the length of the excess repeats, and these hairpin tips can also be cleaved by single strand nucleases at their hairpin tips and by several DNA junction-specific nucleases (Pearson, 2002).

The inability of the repeat-free hairpin to be digested may be due to the presence of the fully-paired DNA that it presents to FAN1, much like the fully-paired duplex to which the flaps are joined, which are also recalcitrant to excision. To this degree the presence of the A•A or T•T mismatches in the (CAG)20 and (CTG)20 slip-outs may facilitate the progressive excision by FAN1. Alternately, FAN1 may have a sequence preference for CAG and CTG repeats.

We tested the role of repeat sequence versus A•A or T•T mismatches using a fully base-paired 5′-(CAG)6→←(CTG)6-3′ hairpin (center inversion point indicated by →←), which would form a fully-paired CAG/CTG intrastrand stem-loop having the same number of nucleotides as the repeat-free hairpin, (CAG)20 and (CTG)20 slip-outs (Figure 4A rightmost panel). This fully-paired 5′-(CAG)6→←(CTG)6-3′ hairpin, like the repeat-free hairpin, would be devoid of A•A or T•T mismatches, and would not be able to re-align between successive cleavages. The 5′-(CAG)6→←(CTG)6-3′ hairpin was recalcitrant to excision, yielding an accumulation of cleaved products identical to the fully-paired repeat-free hairpin (Figure 4A rightmost panel). Thus, A•A and T•T mismatches in the (CAG)20 and (CTG)20 slip-outs are critical for the cleavage patterns of FAN1.

It appears that the perfect repeat-free hairpin is recalcitrant to digestion by FAN1, possibly due to the presence of the fully-paired DNA that it presents to FAN1, much like the fully-paired duplex to which the flaps are joined. To this degree the presence of the A•A or T•T mismatches may facilitate the progressive digestion of FAN1.

### FAN1 processes pre-formed CAA-interrupted CAG slip-outs

The purity of the *HTT* CAG tract is a strong modifier of disease onset (Ciosi et al., 2019; Lee et al., 2019; Wright et al., 2020). In most HD and most non-affected individuals (>95%), the *HTT* repeat CAG tract is interrupted with a single *CAA* motif at the penultimate 3′ end unit of the tract; 5′-(CAG)N-*CAA*-CAG-3′. HD individuals with pure uninterrupted repeat tracts; 5′-(CAG)N-3′, showed hastened disease onset by ∼13-29 years, relative to those with a single interruption. In contrast, individuals with multiply-interrupted repeats; 5′-(CAG)N-***CAA***-CAG-***CAA***-CAG-3′ showed delayed disease onset by ∼3-6 years (Wright et al., 2020). It is thought that the CAA interruptions protect the repeat from length variations. It has long been known that CAA interruptions disrupting the purity of CAG repeat tracts are associated with genetically stable repeats in HD, SCA2, and SCA17 repeats (Findlay Black et al., 2020; Wright et al., 2019, 2020).

Interruptions may stabilize the repeats by either of two routes: Experimental data supports the ability of the CAA interruptions to greatly inhibit the formation of mutagenic slipped-DNA structures, and this is thought to be the source of the reduced instability of the interruptions (Goldberg et al., 1995; Marquis Gacy et al., 1995; Pearson et al., 1998a). Alternately, the presence of CAA interruptions in preformed slip-outs may alter the ability of the slip-outs to be processed by DNA repair proteins. It is notable that the FAN1 *exo*- nucleolytic pause sites on CAG slip-outs localized to the 3′ region that might be affected by the CAA interruptions (Figure S8).

We tested the possibility that FAN1 and its disease-hastening variants may differentially cleave the singly- or doubly-CAA-interrupted slip-outs, versus, the pure CAG slip-outs, where all substrates had a total of twenty repeat units (Figure 4B). Interruption patterns in our substrates matched those present in the *HTT* CAG tract of the genetically stable and delayed-onset HD population (doubly-interrupted), typical age-of-onset (singly-interrupted), and genetically unstable and hastened-onset (pure) HD population (Wright et al., 2020). FAN1 was able to process the CAA-interrupted slip-outs with reduced efficiency compared to pure CAG slip-outs, and two interruptions were more inhibitory than one: (pure (CAG)20(***CAA***CAG)0 > single-interruption (CAG)18(***CAA***CAG)1 > double-interruption (CAG)16(***CAA***CAG)2. It is notable that excision pauses are evident throughout the pure repeat, while pauses are enriched at the 5’ end of CAA-interrupted repeat tracts due to slowed excision, as indicated with arrows. Thus, unlike the arrested excisions upstream of the fully-paired hairpins, the CAA-interrupted slip-outs are processed, yet slower with increasing numbers of interrupts. This suggests that the effect of CAA interruptions may not be limited to inhibiting slip-DNA formation, but may also include differential processing by repair proteins. The FAN1 variants, FAN1^p.R377W^, FAN1^p.R507H^ processed the interrupted slip-outs with comparable efficiencies to FAN1^p.WT^ (Figure S10). In some reactions of the FAN1^p.R507H^, the relative intensity of the strong pause scissile sites was inversed, suggesting that pausing location may be shifted between the pure and interrupted slip-outs from 5′-C↓A↓GC↓A↓GC↓AGCAGCAG-3′ to 5′-C↓A↓G***C***↓***AA***CAG***CAA***CAG-3′ (Figure S10, compare lane3 of leftmost gel to lane 2 of middle to lane 2 of rightmost gel). In these instances, the terminal nucleotide at the strong pause site products would vary.

The above results indicate the manner by which FAN1 may process CAG/CTG slip-outs, begging the question as to how FAN1 may be involved in CGG/CCG expansion diseases.

### Rare missense FAN1 variants in autism individuals with expanded CGG repeats

A shared mechanistic connection of FAN1 to epilepsy, autism spectrum disorder (ASD), and schizophrenia (SCZ) through CGG repeat expansions was recently hypothesized (Deshmukh et al., 2021). This hypothesis was based upon numerous observations. Copy number variants in the 15q13.3 genomic region, containing *FAN1*, have been linked to epilepsy, ASD, and SCZ (Deshmukh et al., 2021). Expansions of CGG at numerous genes (Table S3), have been shown to be involved in epilepsy, ASD, or SCZ. Genetic association studies on SCZ and ASD revealed that, among the six genes present in the 15q13.3 region, *FAN1* was most likely responsible (Ionita-Laza et al., 2014). We recently demonstrated association of tandem repeat expansions to ASD, including CGG (but limited CAG), revealing that expansions are prominent in individuals with ASD relative to their siblings without ASD, and may account for 2.6% of the ASD risk (Trost et al., 2020). We recently made a similar association of SCZ to expanded repeats (Mojarad et al., 2021). Though the mechanism of the association is uncertain, that *Fan1*-knockouts lead to enhanced somatic CGG expansions in FXS mice (Zhao and Usdin, 2018) supports the possibility that genetic variants in *FAN1* may be involved in the enrichment of repeat expansions in ASD (Deshmukh et al., 2021).

Towards delineating a connection of *FAN1* with ASD, we analyzed a cohort of 4,969 individuals with ASD, 1,913 siblings without ASD and their 7,945 parents to determine whether genetic variants in *FAN1* are associated with CGG repeat expansions in individuals with ASD. Size of CGG repeats in the genome (herein as “genome-wide CGG repeats”) was estimated by the sum of genome-wide paired in-repeat read counts for CGG/CCG motifs per sample using ExpansionHunter Denovo, as previously reported (Dolzhenko et al., 2020; Trost et al., 2020). “Expanded CGG repeats” is defined as the presence of a positive paired in-repeat read count (i.e., contains sequence reads that are completely filled with CGG repeats). We determined the correlation between rare (minor allele frequency; MAF <1%) variants (including copy number deletion, copy number duplication, loss of function variants, missense variants and synonymous variants) in *FAN1* and genome-wide CGG repeats (Figure S11). We found that rare missense variants (MAF <0.5%) in *FAN1* were significantly correlated with genome-wide CGG repeats in individuals with ASD (p=0.013; Wilcoxon rank sum test) (Figure 5A).

**Figure 5.**
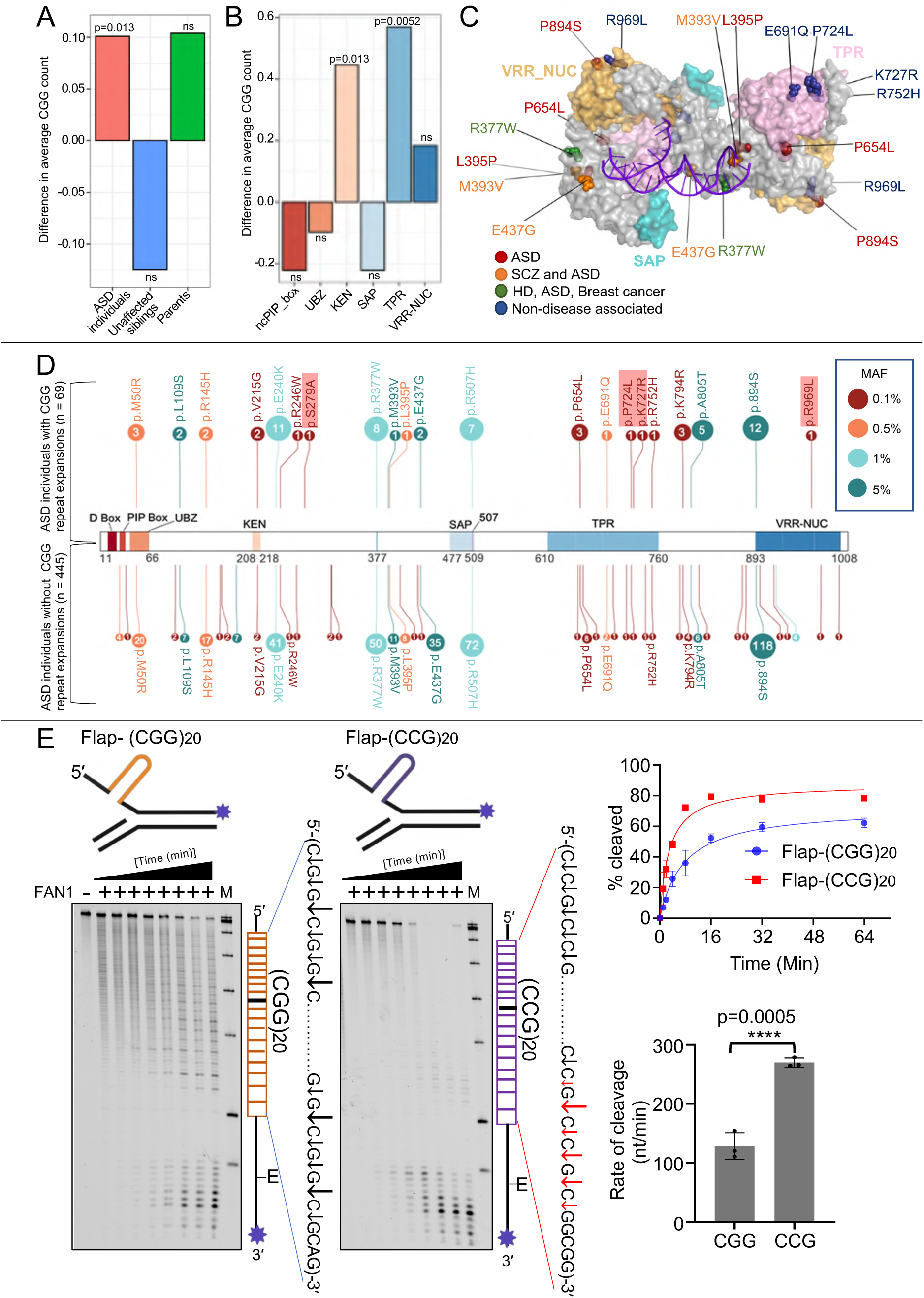
**A) CGG repeat count of ASD individuals with rare missense variants (MAF <0.5%) in *FAN1*.** Mean lengths of CGG repeat count in each population (children with ASD, n=4,969; children without ASD, n=1,913; and parents, n=7,945) are compared between individuals with and without rare missense variants in *FAN1* using Wilcoxon’s test. Tandem repeat count was based on the sum of in-repeat read counts for CGG (/CCG), as calculated by ExpansionHunter Denovo. Positive value of ‘Difference in average CGG count’ represents longer mean lengths in genetic variant carriers. Significant p-values are reported, with non-significant results denoted as ‘ns’. **B) CGG repeat count of ASD individuals with rare missense variants (MAF <0.5%) in annotated protein domains of FAN1 (NM_014967).** Mean lengths of CGG repeat counts in ASD individuals are reported for individuals who have rare missense (MAF <0.5%) variants in functionally annotated FAN1 protein domains using Wilcoxon’s test. Positive value of ‘Difference in average CGG count’ represents longer mean lengths in genetic variant carriers. Significant p-values are reported, with non-significant results denoted as ‘ns’. **C) FAN1 variants identified in ASD individuals with CGG repeat expansions.** Missense variants identified in ASD individuals with CGG repeat expansions are mapped to functionally annotated FAN1 protein (NM_014967) domains and displayed on the dimeric ‘unwinding’ FAN1 protein structure (PDB accession: 4REA). Variants are colored according to previously published disease associations: ASD – red; Schizophrenia (SCZ) and ASD – orange; Huntington’s Disease (HD), ASD and breast cancer – green; no previously reported disease association – blue. **D) FAN1 protein-coding missense variant distribution (NM_014967):** Graph depicts a lollipop plot displaying FAN1 protein-coding missense variants identified in ASD individuals with CGG repeats (top; n = 69), and ASD individuals without genome-wide CGG expansions (bottom; n = 445), relative to a linear schematic representation of the FAN1 protein (NM_014967 isoform). Each variant is depicted as a circle, with the number inside the circle representing the number of times the variant was identified in the cohort and the size of the circle reflecting the variant prevalence in the selected cohort. Variants are coloured according to variant allele frequency; dark red: ≤0.1%, orange: ≤0.5%, light green: ≤1%, dark green: ≤5%. Coloured boxes represent functionally annotated FAN1 domains with numbers representing the domain range on the FAN1 protein. Variants identified only in ASD individuals with CGG repeats but not in those without CGG repeats are highlighted in red. **E) FAN1 *exo*-nucleolytically pauses on CGG/CCG slip-outs:** Time dependant FAN1 *exo*-nuclease activity was performed on 100 nM of CGG/CCG slip-out DNA in presence of 50 nM FAN1. DNAs have 3′-FAM labelled on flap strand (purple star), schematics are shown above each denaturing gel, and labeled strand schematic indicated along the length of the gel, and repeat tract and its center (black line) indicated, “E” denotes elbow region between ds DNA to ssDNA junction. Fractions were collected at each time interval from (0, 2, 4, 8, 16, 32, 64, and 128 minutes). Reactions were stopped by adding 95% formamide, heat denatured and resolved on 6% denaturing PAGE at 2000 V for 1 hr. Image was taken on Typhoon fluorescence channels and analyzed by ImageQuant. Rate of cleavage (nucleotide/minutes) is calculated as described in (Subramanian et al., 2003) and statistically assessed (Welch’s *t* test) per (Kurita et al., 2009), and plotted on GraphPad Prism (Representative gels of n=3 replicates). Error bars represent ±SD. Note: CGG/CCG repeats can form intra-strand hairpins, slipped-DNAs, G4 quadruplexes, Z-DNA, and *i*-motif structures, for ease we present hairpins. gels at 2000 V for 1hr. Representative gel of n=3 replicates, for quantifications see Figure S12. Slip-DNA schematic indicating FAM label (purple star) is shown above the denaturing gel, and labeled strand schematic indicated along the length of the gel, with repeat tract and its center (black line) indicated, “E” denotes elbow region between ds DNA to ssDNA junction. NA does not affect DNA-binding, *endo*- or *exo*-nucleolytic activity per se, as the digestion of the CTG slip-out, which cannot be bound by NA, was unaffected by NA (Figure S12).

Individuals with ASD who carry rare missense variants in specific FAN1 protein domains have significantly more genome-wide CGG repeats (Figure 5B). Our analysis revealed that rare missense variants (MAF < 0.5%) in the KEN-box and TPR domain of FAN1 are significantly correlated with genome-wide CGG repeats in ASD individuals (p=0.013 for KEN domain; p=0.0052 for TPR domain; Wilcoxon rank sum test) (Figure 5B). The missense variants in *FAN1* identified in ASD individuals with expanded CGG repeats are reported in Table S4. Variants that have been previously reported to be associated with ASD, SCZ, HD, KIN and breast cancer were also identified in this cohort (Figure 5C).

We next assessed the association of each missense variant of *FAN1* on genome-wide CGG repeats by comparing their prevalence in ASD individuals with expanded CGG repeats to those without expanded CGG repeats (Figure 5D and Table S4). We identified numerous missense variants, including p.R377W and p.R507H, which had previously been reported as ASD-relevant missense variants (Ionita-Laza et al., 2014), as HD-relevant (Lee et al., 2019), and other disorders (Deshmukh et al., 2021). In our study, these FAN1 variants were present at similar rates between individuals with and without expanded CGG repeats. We identified a novel p.S279A variant that was identified exclusively in one ASD individual with expanded CGG repeats (Figure 5D). The p.P724L (rs761141548) variant in the TPR domain, is among four missense variants that were identified exclusively in ASD individuals who have expanded CGG repeats when compared to those without expanded CGG repeats, and the individual with this variant also harbors a CGG expansion in *FMR1*. We observe that rare (MAF <0.1%) missense variants cluster to the TPR domain, and two of four missense variants identified exclusively in ASD individuals with expanded CGG repeats are located in this domain.

### FAN1 pauses on CGG/CCG slip-outs

From our data above, it appears that FAN1 can process CAG/CTG repeat slip-outs, where the transcriptionally-displaced DNA coding strand, (CAG)N is processed much slower than CTG. This situation might apply to each of the CAG/polyQ diseases, including HD. Were this to extend to the numerous fragile X like repeat expansion loci (Table S3), where the transcriptionally-displaced DNA would be the (CGG)N strand, we might expect that this strand would be processed slower than the (CCG)N strand, by FAN1. We located individual gene-linked repeat loci, using anchored in-repeat reads and determined the strand-specific repeat motif ((CGG)N or (CCG)N) by assessing the transcriptional directions of the loci relative to the gene. We found a trend of more (CGG)N repeat expansions in ASD individuals with rare missense mutations (MAF <0.5%) in FAN1 (Figure S11). However, this trend may be due to more prevalence of expanded (CGG)N repeats than (CCG)N repeats in the genome.

That FAN1 regulates somatic instability of CGG repeats in FXS mice and the association of CGG repeats with intellectual disorders and FAN1 variants in these disorders, prompted us to assess FAN1 cleavage of (CGG)20 and (CCG)20 repeats. Cleavage was unique, cleaving at each nucleotide, with varying preferences. Specifically, FAN1 cleaved 5′-C↓G↓G↓C↓G↓G↓C↓G↓G-3′ and 5′-C↓C↓G↓C↓C↓G↓C↓C↓G-3′, with *exo*-nucleolytic pausing throughout, particularly enriched at the G-C phosphate for each strand (Figure 5E). Digestion rate of CGG was significantly slower than that of CCG (p=0.0005, Welch’s *t* test) (132.273 ± 30.81 and 265.567 ± 0.884 respectively), much like CAG was slower than CTG digestion. Rates of digestion varied by location within the repeat tract, being considerably slower at the beginning, midway and endpoints, consistent with the in-homogeneous cleavage patterns (Figure 5E Table 2). Also, scission hotspots arose in both CGG and CCG slip-outs at the 3′ ends, similar to the intense pauses in the CAG/CTG slip-outs (Figure 2). Thus, FAN1 can also act on CGG/CCG repeat slip-outs, which may explain in part Fan1’s ability to regulate somatic CGG instability in FXS mice (Zhao and Usdin, 2018).

### Naphthyridine-Azaquinolone (NA) alters FAN1 cleavage on (CAG)20 slip-outs

FAN1 appears to protect against hyper-expansions of CAG repeats in HD patient cells and in tissues of HD transgenic mice, as ablation of *FAN1* causes increased CAG expansions (Goold et al., 2019; JM, 2020; Kim et al., 2020). This protective role may be through FAN1’s nuclease activity; acting to remove excess repeats, on slipped-DNA mutagenic intermediates, for example. Genetic ablation of *FAN1* in HD patient cells or HD mice eliminates all of FAN1’s activities, including DNA-binding, dimerization, *endo*- nucleolytic and *exo*-nucleolytic activities.

It has been demonstrated that the expansion bias is the net accumulation of ultra-frequent expansions and contractions (Higham et al., 2012). We tested the hypothesis that FAN1 might protect against hyper-expansions by maintaining an equilibrium of repeat contractions. The CAG slip-out specific small molecule, *N*aphthyridine-*A*zaquinolone (NA), was recently shown to both diminish CAG expansion levels and induce contractions of expanded CAG repeats in HD patient cells and in the striatum of HD mice (Nakamori et al., 2020). NA may exert these effects by modifying the processing of slipped-DNA mutagenic structures by key enzymes, as NA binds specifically throughout the CAG slip-out (Nakamori et al., 2020).

We tested the requirement of FAN1 for the ability of NA to induce contractions of an expanded (CAG)850 tract in an established human cell model of CAG instability (Nakamori et al., 2020). This model shows CAG tract length instability, as detected by small-pool PCR across the repeat, where instability depends upon active transcription across the CTG DNA strand, as occurs at the *HTT* gene (Nakamori et al., 2011). NA was able to significantly induce CAG contractions in these FAN1-proficent cells, as previously demonstrated (Figure 6A, p=0.000059, Pearson’s chi-square test) (Nakamori et al., 2020). siRNA knockdown of *FAN1* exacerbated repeat instability of the expanded tracts (Figure 6A, p=0.00523, Pearson’s chi-square test) in the absence of NA, a finding that is consistent with the protective role of FAN1 against CAG instability (Goold et al., 2019; Kim et al., 2020; Loupe et al., 2020). NA was unable to induce CAG contractions in cells knocked-down for *FAN1* compared to control cells expressing FAN1 (p=0.742, Pearson’s chi-square test) (Figure 6A). Thus, a fully functional FAN1 was required for NA’s induction of CAG contractions in cells.

**Figure 6:**
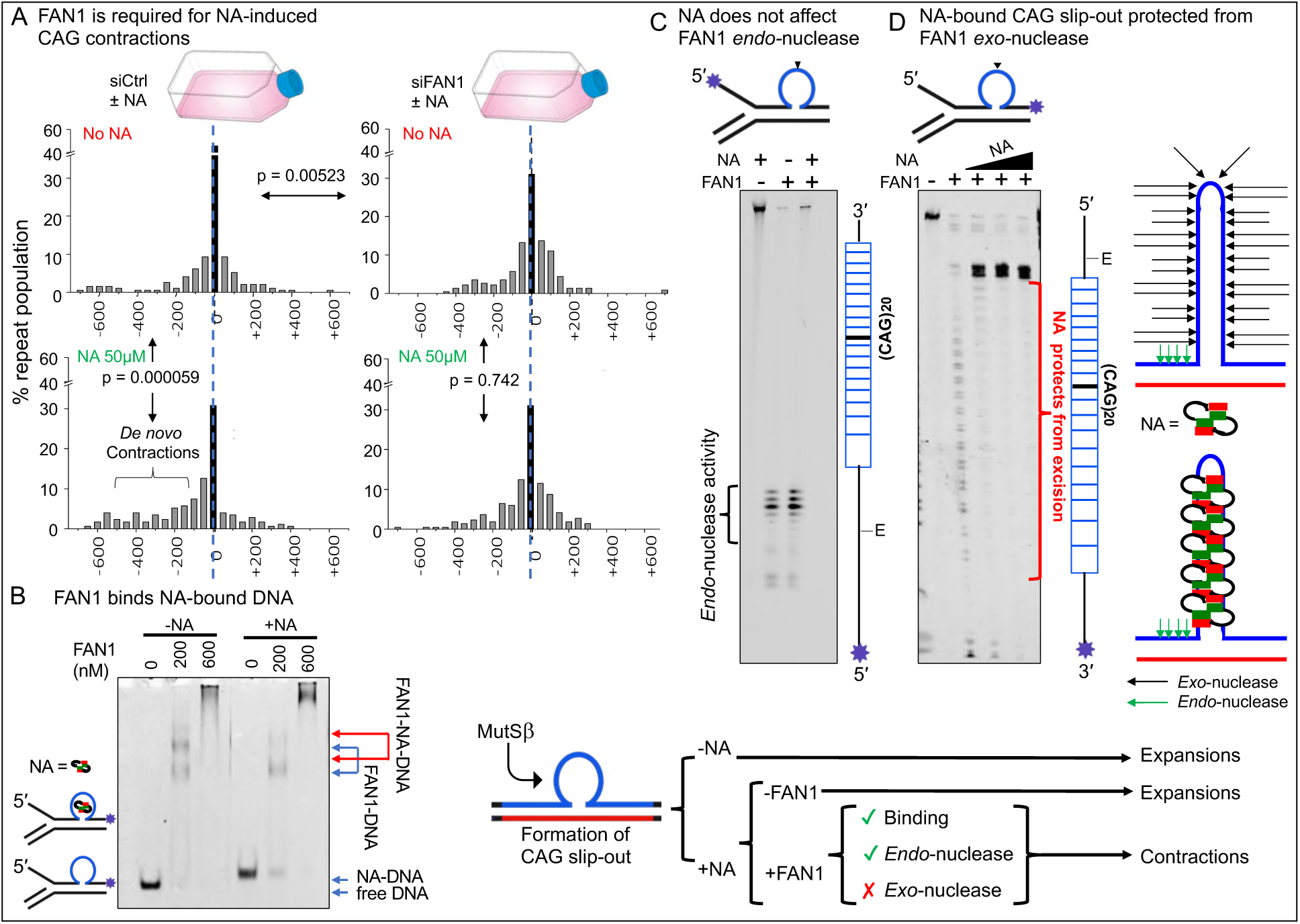
**Naphthyridine-Azaquinolone (NA) protects (CAG)20 slip-outs from FAN1 *exo*-nucleolytic cleavage: A)** si-control and siFAN1-RNA mediated knock-down FAN1 in HT1080(CAG)850 cells treated with NA spanning 40 days. The si-control cells show significant NA-induced CAG contractions (p=0.009) siFAN1-RNA knock-down FAN1 cells do not show NA mediated contraction; **B)** FAN1 binds to NA-bound Duplex-(CAG)20; **C)** FAN1 *endo*-nuclease activity is unaffected on NA-bound Duplex-(CAG)20, where FAM label is at 5′-end of slip-out strand (purple star) +/-10 μM NA. Slip-DNA schematic indicating FAM label (purple star) is shown above the denaturing gel, and labeled strand schematic indicated along the length of the gel, with repeat tract and its center (black line) indicated, “E” denotes elbow region between ds DNA to ssDNA junction. **D)** *Exo*-nuclease activity is blocked on NA-bound Duplex-(CAG)20, where FAM label 3′-end of slip-out strand (purple star). 100 nM of DNA was incubated with increasing concentrations of NA (3.125 μM, 6.25 μM and 12.5 μM) and treated with 100 nM of FAN1, reactions were stopped by adding 95% formamide EDTA, and resolved on 6% denaturing sequencing gels at 2000 V for 1hr. Representative gel of n=3 replicates, for quantifications see Figure S12. Slip-DNA schematic indicating FAM label (purple star) is shown above the denaturing gel, and labeled strand schematic indicated along the length of the gel, with repeat tract and its center (black line) indicated, “E” denotes elbow region between ds DNA to ssDNA junction. NA does not affect DNA-binding, *endo*- or *exo*-nucleolytic activity per se, as the digestion of the CTG slip-out, which cannot be bound by NA, was unaffected by NA (Figure S12).

We next asked if NA might affect one or another of FAN1’s actions on CAG slip-outs? NA-bound CAG slip-outs could still be recognized by FAN1, producing a FAN1-DNA-NA trimolecular complex (Figure 6B). NA-bound CAG slip-outs could still be *endo*- nucleolytically cleaved by FAN1 (Figure 6C). In contrast, NA blocked the *exo*-nucleolytic activity of FAN1 on CAG slip-outs. These results imply that FAN1 is involved in mediating NA-induced CAG contractions, and selectively blocking FAN1’s *exo*-nucleolytic activity on CAG slip-out DNA, while retaining its other activities, can lead to CAG contractions. The protection of the NA-bound CAG slip-out from excision was not the results of enzyme inhibition, as the digestion of the CTG slip-out, which cannot be bound by NA, was unaffected by NA (Figure S12). These results imply that FAN1 is involved in mediating NA-induced CAG contractions: NA-bound CAG slip-outs selectively escaped *exo*- nucleolytic cleavage by FAN1, but could still be bound by FAN1, effectively digested *endo*-nucleolytically, to lead to CAG contractions. It is possible that it is the retention of FAN1 binding, dimerization and *endo*-nuclease activities, while blocking *exo*-nucleolytic activities is what leads to contractions of NA-bound CAG tracts. To this degree, NA is a functional modulator of FAN1’s *exo*-nucleolytic activity on CAG slip-outs, acting specifically at the mutant *HTT* allele. Together, these results support the suggestion that in HD patient brains FAN1 normally acts by preventing hyper CAG expansion mutations through its nuclease activity.

## Discussion

Here, we have advanced the breadth of unusual DNA structures recognized and acted upon by FAN1, and hence broadened the spectrum of processes that may be mediated by FAN1. Notably, we provided persuasive evidence for the binding specificity and striking strand- and structure-specific cleavage patterns of FAN1 to both CAG and CTG slip-outs DNAs.

### Incremental and successive cleavage by FAN1 on slipped-DNAs reveals a manner by which repeat instability might occur

Understanding the action of FAN1 on repeat DNAs is crucial towards learning if and how this potent genetic modifier can alter disease onset by modulating somatic repeat instability. In the case of an inherited expanded repeat, FAN1 appears to be “fighting a losing battle”; FAN1 suppresses against hyper-expansions of CAG and CGG repeats in somatic tissues (Goold et al., 2019; Zhao and Usdin, 2018). The spontaneous repeat expansions that arise in the presence of FAN1, show further enhanced rates and/or greater magnitudes of change in the absence of FAN1. In this manner FAN1 may suppress expansion rates by varying the degree of excision of slip-outs formed during each expansion event. Modulating FAN1 activity by FAN1 levels or functions may alter its ability to modulate somatic instability (Goold et al., 2019).

The “inchworm” repeat digestions by FAN1 suggests a manner by which incremental or “inchworm” growth rates of somatic repeat expansions can occur in patients. Expansions are the net gains of a few repeat units, which are the result of a great many successive mutation events (Higham and Monckton, 2013; Higham et al., 2012; Kaplan et al., 2007). For example, cultured mitotic and post-mitotic HD patient-derived induced pluripotent stem cells (iPSCs) differentiated to medium spiny neurons (MSNs), the vulnerable cell type in HD patients, showed incremental broadening of the expansion sizes over 6-10 months (Goold et al., 2019; Kim et al., 2020). Striatally differentiated cells starting with (CAG)121 showed gains of 1 CAG repeat every 17.7 ± 1.1 days. However, when *FAN1* was knocked-down in those cells, the CAG expansion rate increased to 1 CAG repeat every 9.1 ± 0.6 days (Goold et al., 2019; Kim et al., 2020). These data suggest that FAN1 restrains CAG repeat expansion in cultures that contain a high proportion of differentiated post-mitotic striatal neurons. HD mouse models with (CAG)∼100-180 show a similar pattern of ongoing spontaneous CAG expansions in striatum, showing broadly distributed sizes of additional repeats gained at a rate of ∼3.5 CAG units/month/cell. Recently, in the striatum of HD mice with repeat lengths typical of most HD patients, the age-related somatic expansions of a (CAG)48 over 8 months incurred gains of ∼4-5 CAG repeats, whereas *Fan1*-deficient mice of the same age, that inherited (CAG)48 incurred gains of ∼9-10 repeats; nearly twice as many over the same time (Loupe et al., 2020). Importantly, an intermediate rate of expansions was observed in mice that were heterozygous for functional Fan1. Evidence from post-mortem brains and tissues from HD, SCA1, SCA3, SCA7, SBMA, and FXS patients also supports incremental changes of one repeat unit per mutation event (Hashida et al., 2001; Lokanga et al., 2013; Maia et al., 2017; Martins et al., 2006; Pinto et al., 2020; Santos et al., 2014; Shelbourne et al., 2007; Trang et al., 2015; Watanabe et al., 2000; Yang et al., 2003). Together these findings support a vast number of incremental expansions accumulated over time, and FAN1 levels suppress expansion rates. To date, no enzymatic data can explain this curious incremental mutation process. The successive incremental *exo*-nucleolytic cleavages along the repeat by FAN1 would provide multiple opportunities of step-by-step single repeat expansion events, essentially “ratcheting-down or -up” the tract length.

The highly distributive nature of FAN1 excision, with a lag phase between each event, permits for multiple inter-cleavage opportunities, including structural alteration of the slipped-DNA as well as action by other enzymes (Figure 7A, Figure S13). In the former, the structure of the cleaved DNA product from which FAN1 dissociates, can “breathe” such that the DNA conformation to which FAN1 re-associates may be considerably different from which it just dissociated (Figure 7A, Figure S13). “Inchworm creeps” of a soliton bubble, causing a net translational slippage of the register pairing of the two repeats strands and change the structure of the slip-out between each FAN1 cleavage vent (Figure S13). Such structural dynamics are very rapid, shifting the location of the free-end and the repeat hairpin tip (Pearson, 2002; Slean et al., 2013; Völker et al., 2014; Xu et al., 2020). Between each iterative cycle of FAN1 binding-cleavage-dissociation, there is an opportunity for the fixation of the expansion slip-out by other enzymes such as a polymerase or ligase. That FAN1 excision is particularly slow on CAG slip-outs, might favor such opportunities, over FAN1’s rapid excision of CTG slip-outs (Figure 7B).

**Figure 7:**
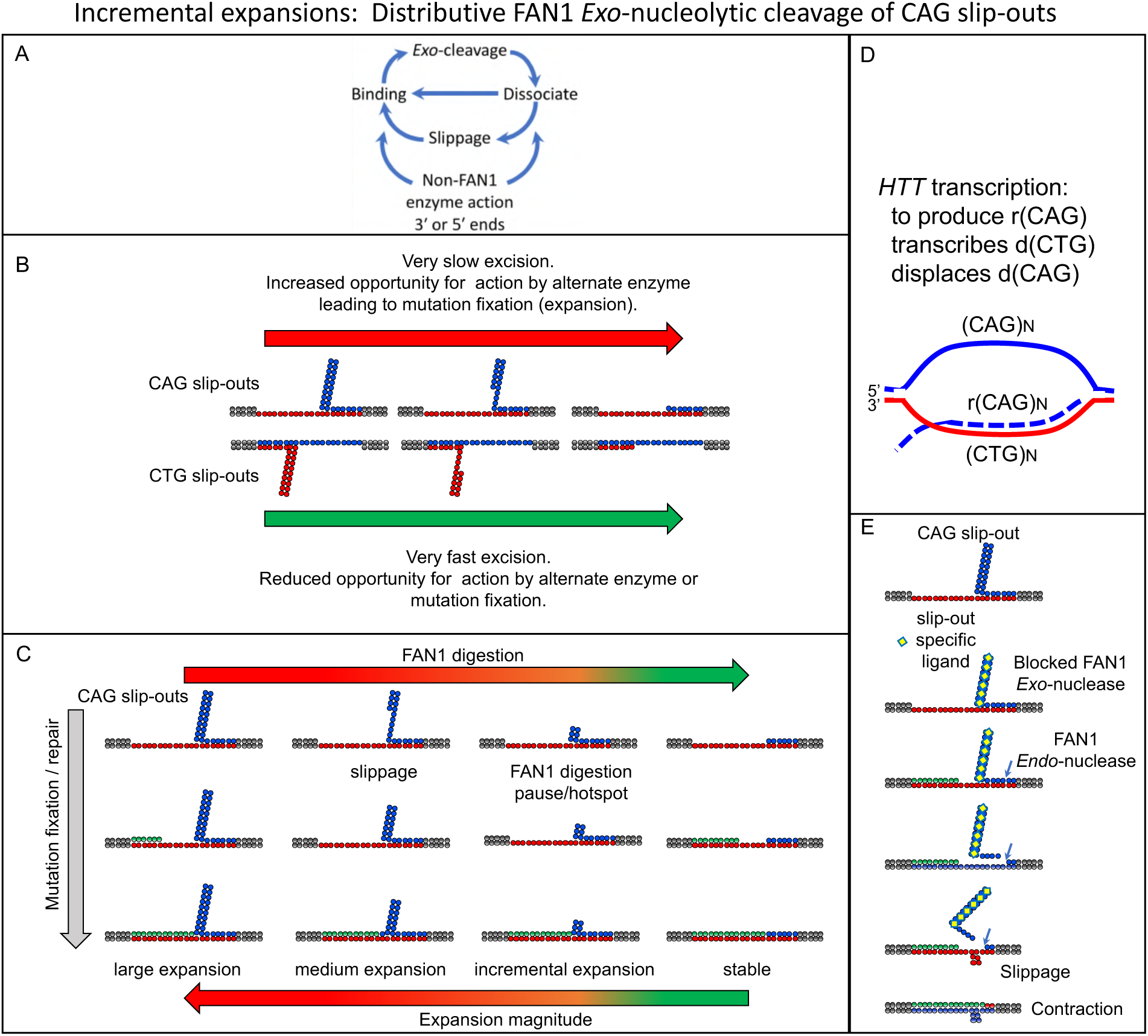
Model: FAN1’s ‘inchworm’ distributive mechanism of successive pausing of repeat excision, slower excision on transcriptionally-displaced CAG strand, than transcribed CTG strand, is well-suited for removing excess repeat slip-outs, to minimize incremental expansions, and allow for NA-induced contractions. See Discussion for details.

### FAN1 on transcriptionally-displaced (coding) strand

FAN1 regulates CAG and CGG repeats in a similar manner. Together, CAG- and CGG-containing genes constitute the two largest classes of repeat expansion diseases. The consistency of FAN1 processing between CAG and CGG repeats is intriguing. Both CAG and CGG repeats showed 1) *in vivo* suppression of somatic repeat instability by FAN1; 2) slower processing of the transcriptionally displaced DNA strand; 3) iterative cycles of incremental excisions throughout the repeat tract; and 4) *exo*-nucleolytic pauses throughout the repeat tract, particularly intense at the 3′ end of the tract.

The differential rates by which CAG and CTG slip-outs are digested by FAN1 may have biological impact. Similarly, CGG, like CAG is digested significantly slower than CCG and CTG. Fifteen of the CAG/CTG repeat diseases have the CAG in the coding strand of the disease gene, where it is the CTG strand that is transcribed to produce an rCAG RNA transcript, thereby displacing the CAG DNA strand, allowing it to form unusual DNA structures. Similarly, 10 of the CGG/CCG repeat genes have the CGG strand that is transcriptionally-displaced. CGG/CCG repeats can form intra-strand hairpins, slipped-DNAs, G4 quadruplexes, Z-DNA, and *i*-motif structures (Chen et al., 1995; Pearson and Sinden, 1996; Pearson et al., 1998a; Usdin and Woodford, 1995; Weisman-Shomer et al., 2000). Transcription across the expanded repeat enhances somatic repeat instability (Lin et al., 2006, 2009; Nakamori et al., 2011, 2020). Interesting parallels exist between somatic hypermutation of immunoglobulin genes and somatic repeat instability in neurodegenerative diseases (Peña-Diaz and Jiricny, 2012; Slean et al., 2008), as both co-opt DNA repair proteins to drive mutations and both are activated by transcription across the region targeted for mutation, a process in both cases, that induces mutagenic DNA structures. Transcription can facilitate formation of slipped-DNAs in the displaced repeat DNA strand (Freudenreich, 2018; Lin et al., 2010; Reddy et al., 2011, 2014). For HD, the *HTT* gene expression is higher in the cortex, caudate and putamen, where somatic CAG expansions are high, than in the cerebellum, where instability is the lowest (Aronin et al., 1995; Stine et al., 1995). In contrast, the antisense *HTTAS* expression, transcribing across the CAG DNA strand of the *HTT* gene, is considerably lower in controls and further reduced in brains of HD patients (Chung et al., 2011). Our finding that FAN1 excision is slower on the transcriptionally-displaced coding strands CAG and CGG, is consistent with a strand bias in repeat instability, and may account for the enrichment of disease loci with CAG and CGG strands in the affected genes. The faster cleavage of CTG and CCG over CAG and CGG respectively, may permit retention of excess CAG and CGG slip-outs (but not CTG or CCG slip-outs), that could lead to expansions.

### FAN1, *exo*-nucleolytic pauses, arrests, A•A/T•T mismatches, & interruptions

We show both pausing and arrest of FAN1 *exo*-nucleolytic activity, which may impact upon repeat instability. Pausing of *exo*-nucleases digestion has only recently been recognized, with little support for its having a biological contribution (Kurita et al., 2009; Lee et al., 2011a; Oijen et al., 2003; Perkins et al., 2003). While the *exo*-nucleolytic digestion of DNA has long been thought to proceed at a homogenous rate, pausing can occur. Unusual DNA conformations, like mismatches or unwound bubbles act as hurdles to nuclease progression (Kurita et al., 2009; Lee et al., 2011a). In one instance, the location of the pause was molecularly mapped to a specific sequence, also thought to slow progression by DNA conformation (Perkins et al., 2003). Our finding that FAN1 digestion paused throughout the repeat tract on slipped-DNAs, at nearly every nucleotide, with preferential pause hotspots at the 3′ end of the tracts, extends the pioneering observations of Perkins *et al*., and van Oijen *et al*., (Oijen et al., 2003; Perkins et al., 2003). Specifically, we observed *exo*-nucleolytic pausing on a biologically-, genetically-, and disease-relevant DNA substrates, by a nuclease that is genetically implicated in those very diseases (HD, numerous SCA’s, ASD), and FAN1 pausing illuminates the process of somatic mutations in those diseases. The cleavage pattern of FAN1 on slipped-DNAs is very different from the limited successive cleavage at every third nucleotide on a non-repeat DNA, that suggested a role of FAN1 in unhooking interstrand crosslinked DNA (Wang et al., 2014). The *exo*-nucleolytic pausing that we observe on slipped-DNAs, may reflect upon FAN1’s role in somatic repeat instability. The size of the incremental expansions, ∼1-5 repeat units/event in cells and tissues of patients and mice, correlates well with the locations of the *exo*-nucleolytic pauses we observe, occurring throughout the CAG, CTG, CGG, or CCG repeat tracts. While excision was faster for CTG than CAG slip-outs, CTG incurred more incision events, cleaving at every nucleotide, while CAG, incurred incisions at only two of every three nucleotides (Figure 7B). Moreover, strong pause hotspots, in each case landing at ∼3-5 repeats from the 3′ end of the repeat tract, indicates that FAN1 does not completely remove all of the slip-out by excision, but leaves ∼3-5 repeats. The extended lifetime or accumulation of these excess repeats may allow for their incorporation, thereby allowing for incremental expansions of ∼1-6 repeat units/event (Figure 7C).

Striking DNA structure-specific variations of FAN1 cleavage were observed. Fully-paired intra-strand hairpins arrested FAN1 excision, leaving greatly reduced cleavage beyond the 5′ base of the hairpin. In contrast, the presence of a CAG or CTG slip-outs, with embedded intrastrand mismatches, clearly affects FAN1 cleavage pattern and rate; slip-outs of CAG are more slowly digested than CTG slip-outs. The A•A and T•T mismatches are critical for the cleavage appear to be required for scission by FAN1. CTG slip-outs assume a more distinct intra-strand hairpin-like structures compared to CAG slip-outs, which interconverts between random-coil and intra-strand hairpin (Hartenstine et al., 2000; Marquis Gacy et al., 1995; Mitas et al., 1995; Pearson, 2002; Teng et al., 2018; Yu et al., 1995). Hairpin formation and stability is sensitive to repeat tract length. Hairpins readily form for tracts >8 repeat units but are diminished as the tract shortens to less than 6 repeats, and can be detected with as few as 3 repeat units. The different stacking interactions of the A•A and T•T mismatches in CAG and CTG hairpins are likely to contribute to hairpin formation and FAN1 action (Tm = 38°C versus 47°C) (Panigrahi et al., 2005; Pearson, 2002; Yu et al., 1995). When CAG and CTG slip-outs are cleaved at C-A and C-T by FAN1, the A•A and T•T mismatches would be the 5′ ends of many of the FAN1 cleaved products, to then serve as substrates for subsequent FAN1 cleavage. These unpaired bases, likely facilitate FAN1’s excision of the slip-outs, consistent with the suggestion of van Oijen et al (Oijen et al., 2003) that base-pair melting is the rate-limiting step in the action of lambda *exo*-nuclease. The inability of FAN1 to excise the fully-paired hairpins, supports the involvement of the intra-strand mismatches. Slipped-out repeats can undergo rapid structural fluctuations (hairpin realignments, hairpin tip location, hairpin tip size) (Pearson, 2002; Slean et al., 2013; Völker et al., 2014). The conformations assumed would be sensitive to repeat tract length the nucleotide at the 5′ end—both of which would change with successive FAN1 cleavage events (Figure 7A, Figure S13).

Both interruptions of the CAG tract purity and DNA repair gene variants are major modifiers of disease onset for HD, SCA1, SCA2, SCA3, and SCA17 (Ciosi et al., 2019; Findlay Black et al., 2020; Gao et al., 2008; Lee et al., 2019; Wright et al., 2019). This is of particular interest, since FAN1, one of the strongest HD modifiers, may processes the pure and CAA interrupted CAG tract differently. Interruptions of the CAG tract by CAA units in HD, SCA1, SCA2, SCA3 can delay disease onset by ∼13-29 years disease. Since CAA-interrupted repeat lengths are genetically and somatically more stable, it is thought that the interruptions protect the repeat from instability by two non-exclusive paths: first by blocking the formation of slipped-DNAs, or second, by altered enzymatic processing of the interrupted repeats. The former concept is supported by biophysical data from numerous groups (Goldberg et al., 1995; Marquis Gacy et al., 1995; Pearson et al., 1998a). The latter is supported by the partial loss of stabilization of interrupted repeat in DNA repair defective yeasts, via coexcision of slip-outs containing interruptions (Rolfsmeier et al., 2000). Our finding that FAN1 poorly excises the singly- and even worse for doubly-interrupted slip-outs, supports involvement of both DNA structural and repair protein paths by which interruptions can offer protection.

FAN1, one of the strongest in-*trans* HD modifiers, can, at least *in vitro*, differentially process one of the strongest in-*cis* HD modifiers, CAG tract purity. The presence of a CAA motif in an intrastrand CAG hairpin would produce two consecutive mismatches; following the C-G pair, would be A•A and A•G mismatches, producing a “4-base bubble”. Such tandem mismatches may present differently to FAN1 compared to the embedded A•A and T•T mismatches in pure CAG and CTG slip-outs. The ability to form intrastrand hairpins is impaired by CAA-interrupted CAG tracts (Goldberg et al., 1995; Marquis Gacy et al., 1995; Pearson et al., 1998a), essentially by “telestability” (AKA DNA allostery), where the sequence of one region of a duplex DNA affected the physical properties of a contiguous but remote region (Burd et al., 1975; Chaires, 2008; Goldberg et al., 1995). This is consistent with our observed overall poor FAN1 digestion of the interrupted tracts relative to the pure tracts. However, once formed, the interrupted slip-outs could be processed by FAN1. When interrupted CAG slip-outs are cleaved at C-A of CAA, this would create tandem A•A and A•C mismatches at the 5′ ends of the FAN1 cleaved products. These dangling unpaired bases may affect the exo-nucleolytic pausing pattern of FAN1, its variants, and of other enzymes. Terminal sequence effects upon slippage have been observed. Changing the terminal nucleotide of a repeat tract duplex to a non-repeat nucleotide efficiently inhibited slippage during polymerase-mediated reiterative synthesis (Chamberlin and Berg, 1964; Olivera and Lehman, 1968). This end effect on slippage is thought to be due to the altered structures of the dangling ends (Banavali, 2013; Bommarito et al., 2000; Rokita and Romero-Fredes, 1989).

CAG hairpins with recessed 5′ ends, similar to the FAN1 substrates used herein, can rapidly and spontaneous undergo dynamic slippage, thereby shifting the location and conformation of the hairpin tip (Xu et al., 2020) (Figure S13). Interrupting CAA units in the CAG tract could dramatically reduce strand slippage and/or destabilize the hairpin, an effect that in preformed hairpins, depended upon the location of the CAA unit (Xu et al., 2020). It is notable that the FAN1 *exo*-nucleolytic pauses localized to the 3′ ends of the repeat slip-outs to the last 3-4 repeat units, upstream of the ssDNA-dsDNA junctions, where significant conformational fluctuations (“DNA breathing”) are likely to occur (Von Hippel et al., 2013; Jose et al., 2009; Pearson, 2002; Slean et al., 2013; Völker et al., 2014). Regarding the fixed/anchored hairpin tip of the non-repeat hairpin: It is not surprising that the worst substrate of FAN1 is a short 5′-overhang (Kratz et al., 2010), which would be the product produced as FAN1 approaches the shortest tract of remaining repeat at the 3′ end.

### FAN1 coding variants in ASD individuals with expanded CGG repeats

FAN1 may regulate the somatic stability of many CGG tract-containing genes, and in this manner could modify the penetrance of disorders, such as ASD, SCZ, FXTAS, FXPOI, or other. Gene-specific CGG repeat expansions have recently been recognized to be extremely common and many are likely to cause clinically related disorders. Recently, multiple new disease-causing CGG repeat expansions loci (Ishiura and Tsuji, 2020; Ishiura et al., 2019; Sone et al., 2019) , as well as more than 1000 expandable CGG tracts (Annear et al., 2021; Garg et al., 2020; Mitra et al., 2021; Trost et al., 2020; York and Lat, 2021), including fragile X at *FMR1,* have been identified (Debacker and Frank Kooy, 2007) . Where a disease link has been made, most instances show partial expression of clinical symptoms which depends upon the CGG expansion size and expression (FXTAS, FXPOI, autism, intellectual disorder, schizophrenia, epilepsy, and more). The expanded *FMR1* CGG repeat experiences high levels of post-zygotic repeat instability, evident as ‘size mosaicism’, ranging from premutation to full mutation repeat expansions within or between tissues of the same individual. Essentially, >80% of individuals with expansions display mosaicism (Jiraanont et al., 2017; Nolin et al., 1994; Zhao and Usdin, 2016). Data from FXS families suggests that multistep *FMR1* CGG mutations involve the gain or loss of several CGG units per event (Lokanga et al., 2013). Thus, if FAN1 regulates all CGG repeat loci like the *FMR1* CGG tract (Zhao and Usdin, 2018), then suggests that the importance of this protein is yet to be fully recognized.

Previously, a range of genetic variants of FAN1 were significantly associated with individuals with ASD and SCZ (Ionita-Laza et al., 2014). Recently, Dumas et al., identified *FAN1* and several other DNA damage response proteins associated with autism and brain diseases, to be amongst the highest positively selected genes during human brain evolution, and thus most highly conserved in the human genome (Dumas et al., 2021). This further highlights the importance of DNA repair genes in maintaining the genome and disease modification. Similarly, copy number variants of FAN1 arising in 15q13.3 microduplication/microdeletion syndrome families, is associated with numerous symptoms of developmental delay (reviewed in (Deshmukh et al., 2021)). Here, we report the identification of rare nonsynonymous variants located within a several key functional regions of FAN1. We also found a significant association between these rare missense variants of FAN1 and increased genome-wide CGG repeat expansions in individuals with ASD. Some of the variants we found have been previously identified in ASD, SCZ, and HD, suggesting a potential shared modifying effect by FAN1, possibly through mediation of repeat instability. The rare variants that we identified in *FAN1* may increase risk of ASD, but the relative risk is probably modest. Given the low frequency of these variants (most are singletons), it is very difficult to estimate relative risks for each individual variant without additional information. Future studies will test the effects of the identified ASD-associated FAN1 variants.

### FAN1-NA coupling to induce CAG repeat contractions

Naphthyridine azaquinolone (NA), which specifically binds to CAG slip-out structures, was recently revealed to induce contractions of expanded repeats *in vivo* in the striatum of HD mice (Nakamori et al., 2020). The ability of NA to induce contractions required transcription across the expanded CTG repeat, displacing the CAG DNA strand. The transcriptionally-displaced CAG DNA strand could form CAG slip-outs, which become bound by NA, that were revealed to be protected from repair, and lead repeat contractions. In the initial study, we demonstrated that the NA-induced CAG contractions required the MutSβ (MSH2-MSH3), a mismatch repair complex known to be required for CAG expansions in HD mice. Notably, NA-bound CAG slip-outs could still be bound by MutSβ (Nakamori et al., 2020), suggesting that MutSβ may act to facilitate slipped-DNA formation, even in the presence of NA. In this initial assessment, a nuclease was suggested to be involved in the NA-induced CAG contractions.

Here we reveal that a fully-functional FAN1 is required for NA’s ability to induce CAG contractions. This finding links FAN1 to slipped-CAG DNAs, as these are targeted by NA, and to the contraction process, in keeping with the suspicion that FAN1 protects against hyper-expansions of CAG repeats through its nuclease activity — removing excess repeats. Our data also reveal that the NA-bound CAG slip-out can still be recognized and bound, and *endo*-nucleolytically cleaved by FAN1. However, NA bound slip-out is protected from FAN1 from *exo*-nucleolytic cleavage (Figure 7E). FAN1 can *endo*- nucleolytically cleave beyond the repeat tracts so as to remove the excess CAG repeat DNA. Thus, inhibiting only *exo*-nucleolytic activity while retaining DNA-binding and *endo*- nucleolytic activity appears to be critical to induce CAG contractions. Thus, screening for selective inhibitors of FAN1’s *exo*-nucleolytic activity, but permit FAN1 to retain its other activities (DNA-binding, *endo*-nucleolytic activity, dimerization, interaction with other proteins, etc.), would lead to CAG contractions. It is possible that it is the retention of FAN1’s DNA binding, dimerization, and *endo*-nuclease activity, while blocking its *exo*-- nucleolytic activity in-action, is what leads to CAG contractions. To this degree, NA is a functional modulator of FAN1’s *exo*-nucleolytic activity on CAG slip-outs, acting specifically at the NA-bound mutant *HTT* allele. This may serve as a guide as to how a FAN1-centered drug screen should be conducted; rather than upregulating FAN1 expression or enhancing both *endo*- and *exo*-nuclease activities, which may subject the whole genome to possible deleterious effects. Together, these results support the suggestion that in HD patient brains FAN1 normally acts by preventing hyper CAG expansion mutations through its nuclease activity (Figure 7).

### HD-disease modifying FAN1 variants

Elucidating the functional impact of the disease modifying FAN1 variants either on CAG instability or another disease-modifying path, is a challenging and complex task (reviewed in (Deshmukh et al., 2021). It is still premature to have a clear view as to the manner by which the disease-associated variants of *FAN1* may hasten- or delay-disease onset (complications of this have recently been reviewed (Deshmukh et al., 2021). Here we have begun to assess the action of FAN1 upon disease-associated repeats and the unusual DNA structures they may assume during repeat instability. The disease-delaying FAN1 variants in the non-coding regions of the *FAN1* gene (15AM2 and rs3512) associate with increased FAN1 expression, leading to delay disease onset by reducing somatic CAG expansions in the brains of affected individuals (Ciosi et al., 2019; Lee et al., 2019). Evidence supports a dose-effect of Fan1 on somatic CAG instability in the brains of HD mice (Loupe et al., 2020). Notably, mice with two, one, or no functional *Fan1* genes (*Fan1*+/+, *Fan1*+/, and *Fan1*-/-) show progressively increased rates of somatic CAG expansions (Loupe et al., 2020). Our finding that the excision rate of excess slip-outs increases with increased concentrations of FAN1, supports excision as the limiting factor in the disease-delaying FAN1 variants. On the other hand, the disease-hastening (coding) FAN1 variants have not revealed strong functional differences. Mild differences between FAN1 forms were observed for DNA binding, FAN1-FAN1 dimerization, and *endo*- or *exo*- nuclease activity (Figure S2, S3, S4 and S7). We showed for multiple DNA substrates that FAN1 binding affinities and *exo*-nuclease rates with very mild insignificant differences in between FAN1 forms. There is no obvious pattern for DNA binding and nuclease activity. Thus, it is unlikely that the varied nuclease rates are a reflection of the binding affinity alone.

Our biochemically controlled DNA-binding experiments using multiple preparations of highly-purified FAN1 from baculovirus-infected insect cells, contrast with the recent report that FAN1^p.R377W^ and FAN1^p.R507H^ showed decreased DNA binding relative to FAN1^p.WT^ (Kim et al., 2020). The discrepancy is likely due to technical differences (Kim et al., 2020). Also, Goold et al were able to detect enriched binding of FAN1 to the expanded *HTT* CAG repeats by ChIP, but did not observe differences in binding between FAN1^p.R507H^ and wildtype FAN1 (Goold et al., 2019). Importantly, DNA binding capacity of FAN1 mutants does not always reflect its nuclease activity: We show that FAN1^p.D960A^, but not FAN1^p.D981A-R982A^, retains slipped-DNA binding capacity comparable to FAN11^p.WT^, while both FAN1^p.D960A^ and FAN1^p.D981A-R982A^ are devoid of nuclease activity. In contrast, the triple mutant FAN1^p.K525E/R526E/K528E^ had near undetectable DNA binding by band-shift, undetectable *endo*-nuclease activity, but retained *exo*-nuclease activity comparable to wildtype FAN1(Zhao et al., 2014). Thus, discordance of FAN1 DNA binding activity from nuclease activity has been reported (Kratz et al., 2010; Liu et al., 2010; MacKay et al., 2010; Zhao et al., 2014).

Are the missense variants of FAN1 associated with hastened-disease onset, affected in any of the FAN1 functions? Our current analyses do not reveal an obvious effect of the FAN1^p.R377W^ and FAN1^p.R507H^ variants on any of the tested functions of FAN1 on CAG/CTG repeat DNAs. Specifically, DNA binding, FAN1-FAN1 dimerization, *endo*- or *exo*-nucleolytic activity (efficiency rates, patterns) are comparable to the wild type FAN1. This is based upon activity assessed upon thirteen different DNA substrates (+/- CAG/CTG), using 4-5 independent highly-purified full-length preparations of each FAN1 form, and 3-5 technical replicates of each test. Future goals include assessing the action of FAN1 and its interacting partners upon slipped-DNAs. FAN1 interacts with the mismatch repair (MMR) proteins, MLH1, PMS2, PMS1, MLH3 (Cannavo et al., 2007), PCNA (Porro et al., 2017), and FANCD2 (Kratz et al., 2010; MacKay et al., 2010; Smogorzewska et al., 2010). FAN1 may interact with the MMR complexes MutLα (MLH1-PMS2), MutLβ (MLH1-PMS2) and MutL*γ* (MLH1-MLH3) via their common MLH1 subunit. The residues of FAN1 that interact with MMR proteins are unknown. Alternately, FAN1 may interact with each of the MutL homologs independent of each other, as recent evidence supports MLH1-independent functions of PMS2 and MLH3 (Rahman et al., 2020). The requirement of the MMR complexes—including MutSα (MSH2-MSH6), MutSβ (MSH2-MSH3), MutLα, MutLβ, and MutL*γ*—for the expansions of CAG/CTG, CGG/CCG, and GAA/TTC repeats has been extensively studied (Schmidt and Pearson, 2016). While it is tempting to speculate the action of FAN1 and its interacting partners upon slipped-DNAs, this is likely to be complex.

### Therapeutic target

It is unclear whether FAN1 may be a safe therapeutic target for repeat expansion diseases. Bona fide autosomal recessive loss of function *FAN1* mutations are definitively linked with karyomegalic interstitial nephritis KIN (OMIM #614817) (Deshmukh et al., 2021; Zhou et al., 2012). Deficiencies of FAN1 lead to mildly increased sensitivities to interstrand crosslinking agents, like mitomycin C, and mildly increased levels of chromatid breaks and radial chromosome, but both were less than enhanced levels associated with Fanconi anemia. Rather than inhibiting or knocking-down FAN1, for repeat expansion disease, increasing the levels and/or the activity of FAN1 seems to be a wanting beneficial path. It was proposed that the FAN1 variants may increase or decrease somatic CAG expansions, which lead to increases or decrease in the rate of HD pathogenesis leading to disease onset (Loupe et al., 2020). Were one to screen for chemical matter that completely ablated all FAN1 activities, would likely identify inhibitors that lead to enhanced CAG expansions. Enhancing FAN1 levels or activities would be expected to reduce levels of somatic CAG expansion. However, over expressing a nuclease could have deleterious effects on genome stability. The *endo*- and *exo*-nuclease activities of FAN1 may have distinct functions, as has been shown for MRE11 and EXO1 (Morafraile et al., 2020; Shibata et al., 2014). Blocking specific FAN1 activities but not others from acting on the repeats, may well lead to effects very different from complete ablation of FAN1, as we have demonstrated.

## Conclusion

A functional role of FAN1 in repeat instability is emerging. Current evidence suggests a role for *FAN1* in suppressing, but not ablating, spontaneous incremental somatic CAG expansions. In the absence of FAN1, either the frequency of expansion events or the magnitude of the expansion is increased. FAN1 protects against hyper-expansions of CAG and CGG repeats in HD and FXS patient cells and in tissues of transgenic mice (Goold et al., 2019; Kim et al., 2020; Loupe et al., 2020; Zhao and Usdin, 2018). Genetic ablation of *Fan1* in HD mice, eliminates *all* of Fan1’s activities, including DNA-binding, dimerization, *endo*-nucleolytic and *exo*-nucleolytic activities. The data presented here suggest that the protective role of FAN1 against somatic hyper-expansions may be through FAN1’s nuclease activity; acting to remove excess repeats. Specifically, the distributive action of FAN1, incurring iterative cycles of incisions with *exo*-nucleolytic pauses in the slipped-out repeats, suggest a manner by which the incremental expansions may be removed. The sustained pauses, may allow for some expansions to escape excision. That this is particularly exacerbated on the transcriptionally displaced repeat strand, supports this action in post-mitotic tissues. The structure specificity of FAN1 digestions of slipped-repeats, and the poor excision of their interrupted forms, supports novel paths by which a nuclease and tract interruptions can offer protection against incremental somatic repeat expansions.

## ACKNOWLEDGMENTS

This work was partially funded by grants from the Canadian Institutes of Health Research (CIHR FRN148910 to C.E.P.; CIHR FDN-388879 to J.Y.M.; CIHR to R.K.C.Y.), the Hereditary Disease Foundation (to A.L.D.). C.E.P. holds a Tier 1 Canada Research Chair in Disease-Associated Genome Instability. J.Y.M holds a Tier 1 Canada Research Chair in DNA Repair and Cancer Therapeutics.

This work was partially supported by the Canadian Institutes of Health Research (FRN388879, J.-Y.M. and FRN148910, C.E.P.), Natural Sciences and Engineering Research Council (RGPIN-2016-08355, C.E.P.), Muscular Dystrophy Canada (C.E.P.), Tribute Communities (C.E.P.), The Petroff Family Fund (C.E.P.), The Kazman Family Fund (C.E.P.), The Marigold Foundation (C.E.P.), the Hereditary Disease Foundation (to A.L.D.). The National Center of Neurology and Psychiatry (29-4, M.N.), a JSPS KAKENHI Grant-in-Aid for Young Scientists (Start-up A, 24890110 and 25713034, M.N.), Scientific Research (B, 16H05321, M.N.) and Specially Promoted Research (26000007, K.N.), J.Y.M holds a Tier 1 Canada Research Chair in DNA Repair and Cancer Therapeutics. C.E.P. holds a Tier 1 Canada Research Chair in Disease-Associated Genome Instability.

## Contributions

ALD performed all *endo*-/*exo*-nuclease and DNA binding experiments, MC and JYM purified all FAN1 proteins and performed FAN1-FAN1 dimerization experiments, MM, SL, GBP, AF, PW performed preliminary experiments. MK, EB, RY performed all bioinformatic experiments. MN performed cell treatment and small-pool PCR repeat length analysis. KN synthesized NA the CAG slip-out ligand. NS generated supplementary Table S3. CEP, ALD, MM, SL, GBP, MK, WB, RY, MC, JYM conceived experiments analyzed data and wrote manuscript. All authors discussed the results and commented on manuscript.

## Materials and Methods

### DNA substrate preparation

Plasmids containing human DM1 genomic (CTG)N•(CAG)N repeats (n=30 or 50) and human nonrepeating sequences flanking the repeat have been previously described (Pearson and Sinden, 1996; Pearson et al., 1997, 1998c, 1998b). (CAG)50•(CAG)30, (CAG)30•(CAG)50 and, (CAG)50•(CAG)50 duplex structures are formed by denaturation and renaturation of plasmids as described (Pearson and Sinden, 1996; Pearson et al., 1997). Each structure is resolved and purified from 4% PAGE, then end labelled by Klenow small polymerase and alpha dNTPs.

All oligo nucleotides are purchased from IDT and are shown in Table S1. Substrates were generated by heating respective oligonucleotides at equimolar concentrations at 95°C, in annealing buffer (10 mM Tris pH 8, 100 mM NaCl and 1 mM EDTA) followed by slow cooling at room temperature. The substrates were checked on 4% PAGE for annealing.

### FAN1 cloning

Wild-type human *FAN1* gene was cloned in a derivative of pFASTBAC1 (Invitrogen) with GST-, FLAG and His_10_ tags. FAN1 variants were introduced in the pFASTBAC1-GST-FAN1-FLAG-His plasmid by site-directed mutagenesis using the Q5 Site-Directed Mutagenesis Kit (New England Biolabs). Confirmed by DNA sequencing. All primers used for site-directed mutagenesis are listed in Table S4.

### FAN1 protein expression and purification

Recombinant wild-type and mutant FAN1 were tagged at the N terminus with GST and at the C terminus with His10 and purified according to a protocol described previously (Maity et al., 2013). Multiple preparations of each FAN1 variant were produced, purity confirmed by SDS-PAGE, detecting only a single electrophoretic species, and no degradation products. Our FAN1 preparations were free of contaminating nucleases, as multiple preparations of the nuclease-dead FAN1^p.D960A^, known to be deficient in *endo*- and *exo-*nucleolytic activities (Smogorzewska et al., 2010; Zhao et al., 2014), were, as expected, consistently nuclease-free. This quality control avoids inappropriate assignment of variable nuclease activity (Bregenhorn and Jiricny, 2014; Shao et al., 2014).

### DNA-binding (electrophoretic mobility shift) assay

Electrophoretic mobility shift assays were performed using 50 nM of alpha dNTPs end filled excess (CAG/CTG), 50L (CAG)50•(CTG)50 and duplex repeat free pUC19 DNA substrates. DNAs were incubated with 0-600 nM (0, 10, 20, 40, 100, 400, and 600 nM) purified FAN1^p.WT^ / FAN1^p.R377W^ / FAN1^p.R507H^ in binding buffer (50 mM Tris HCl pH 8.0, 25 mM NaCl, 10 mM MgCl_2_, 1 mM dithiothreitol, 200 μg/ml BSA), plus 100 ng/reaction poly(dIdC) in 10 μL reactions and incubated at 37°C for 20 minutes. Reaction products were separated on a 4% polyacrylamide gel, dried and exposed for autoradiography. The similar electrophoretic mobility shift assay performed on flap DNA substrates using 5 nM of 3′-end FAM labelled substrate were incubated with 0-600 nM of FAN1 and its variants in binding buffer plus 25 ng/reaction of poly(dIdC) in 10 μL reactions and incubated on ice for 20 minutes. Reaction products were separated on 4% polyacrylamide gel and scanned in Typhoon FLA in fluorescence filter. All quantifications were done in ImageQuant software and represents three experiments.

### Nuclease assays

FAN1 nuclease assays were performed as described in (MacKay et al., 2010) with 100 nM of fluorescently labeled DNA. Reactions were initiated by the addition of protein, incubated at 37°C, for 0-20 minutes then stopped with formamide loading buffer (95% formamide, 10 mM EDTA). Products of were separated using 6% denaturing sequencing gel for 1 hr at 2000 V and detected at fluorescence filter in the Typhoon FLA (GE Healthcare). Rate of cleavage (nucleotide/minutes) is calculated as described in (Subramanian et al., 2003) and statistically assessed as per (Kurita et al., 2009), and plotted on GraphPad Prism (n=3).

### FAN1-FAN1 dimerization (BMH protein crosslinking) assay

BMH (bismaleimidohexane) protein crosslinking assay was carried out as described previously (Zhao et al., 2014).

---

**Figure S1:**
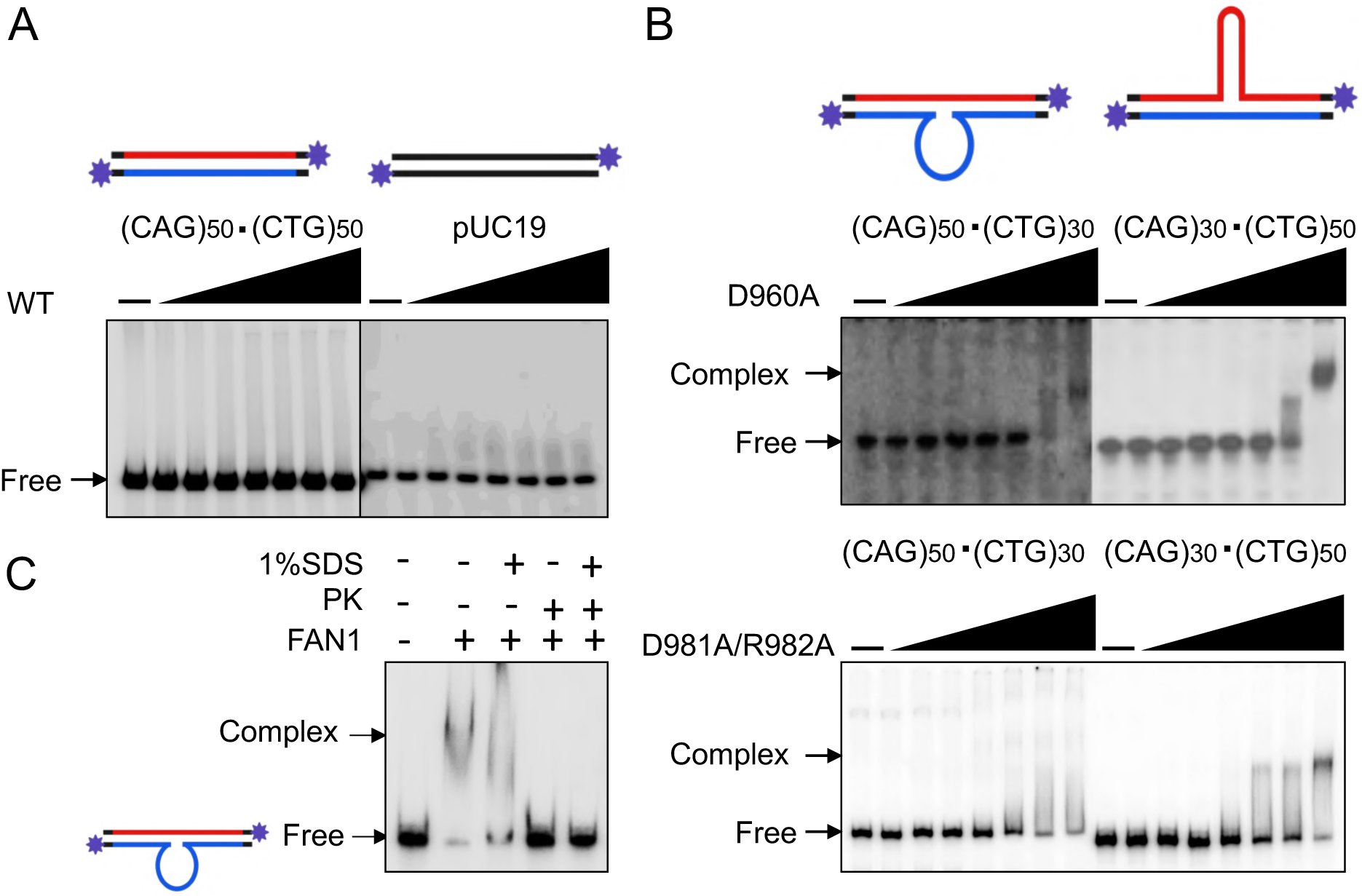
(A) Representative DNA-binding (mobility-shift) assay of FAN1^WT^ on fully-paired repeat strands, (CAG)50•(CTG)50, duplex repeat-free DNA (pUC19). (B) DNA-binding of FAN1 D960A and D981A-R982A on excess CAG/CTG slip-out DNAs: Concentration-dependent DNA-binding is carried out from 10-600 nM of FAN1^p.D960A^ (10, 20, 40, 100, 200, 400 and 600 nM) and FAN1^p.D981A-R982A^ from 200-3200 nM (200, 400, 600, 800, 1200, 1600 and 3200 nM), on 5 nM (3′-end filled with ^32^P-*α*dNTPs, purple star). Samples were resolved on 4% PAGE for 1 hr 30 min at room temperature, gel was dried and exposed to Phosphor Imager and observed under Typhoon-FLA8000. (C) FAN1 DNA-binding is non-covalent; FAN1-DNA complex is treated with SDS or Proteinase K or both and observed loss of shifted DNA complex on native 4% PAGE. (Representative gels of n=3 independent replicates).

**Figure S2:**
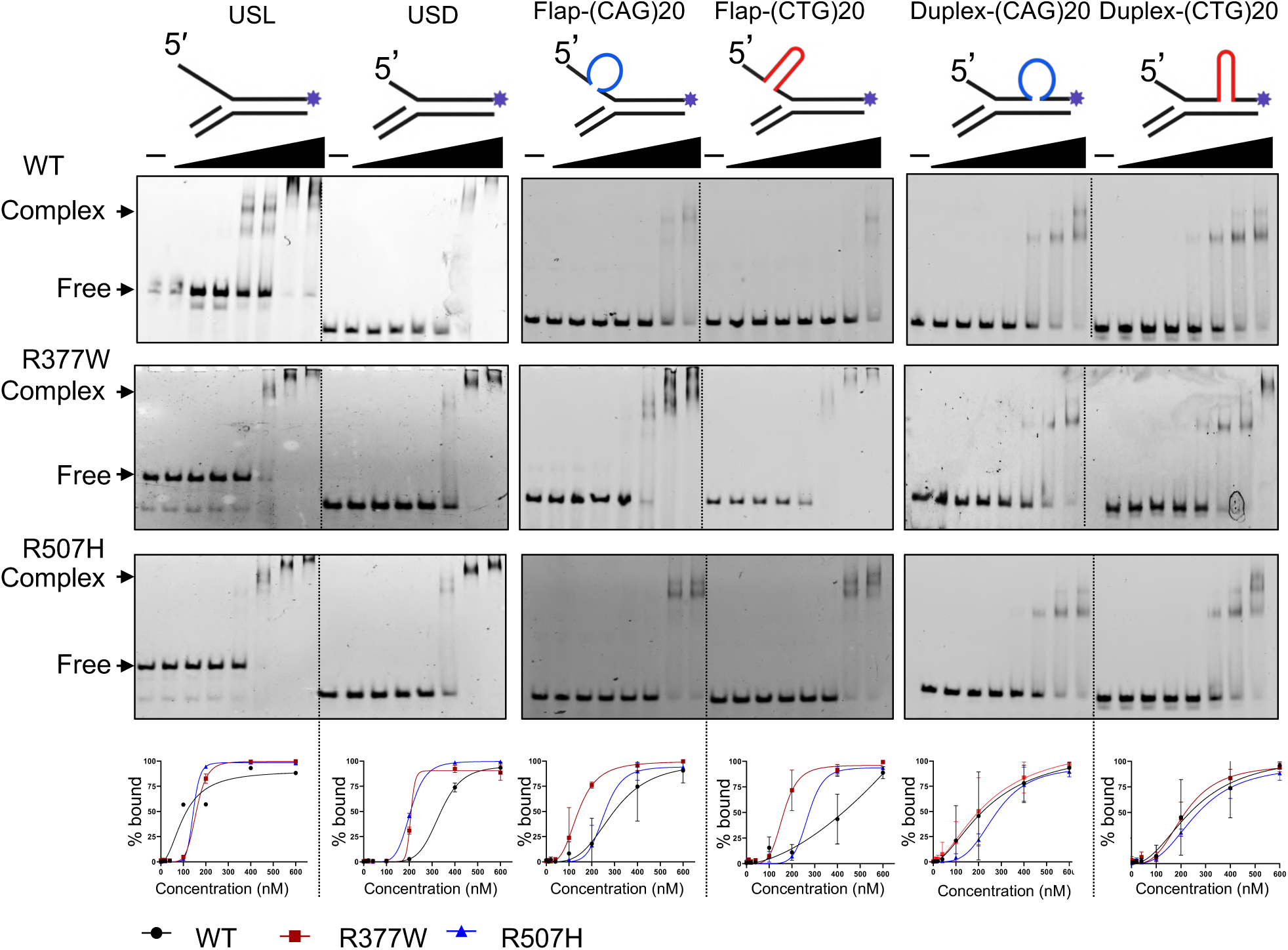
**FAN1 binds flap-bourne and duplex-bourne slip-out DNAs**: Representative DNA mobility shift assay of FAN1 FAN1^p.WT^, FAN1^p.R377W^, and FAN1^p.R507H^ on 3′-FAM labelled unstructured flap substrates (purple star), flap bourne slip-out and duplex bourne slip-out, the schematic of each DNA substrates is on the top of the gel. Concentration-dependent DNA binding is carried out from 10-600 nM of FAN1 (10, 20, 40, 100, 200, 400 and 600 nM) on 5 nM 3′-FAM-labeled DNA substrates. Samples were resolved on 4% PAGE for 3 hrs at 4°C and observed under Typhoon-FLA8000 in fluorescence channel at 100-200V. Data were analyzed on ImageQuant and plotted on GraphPad Prism 8.2 (Representative gels of n=3 independent replicates). Quantified results are plotted below each gel.

**Figure S3:**
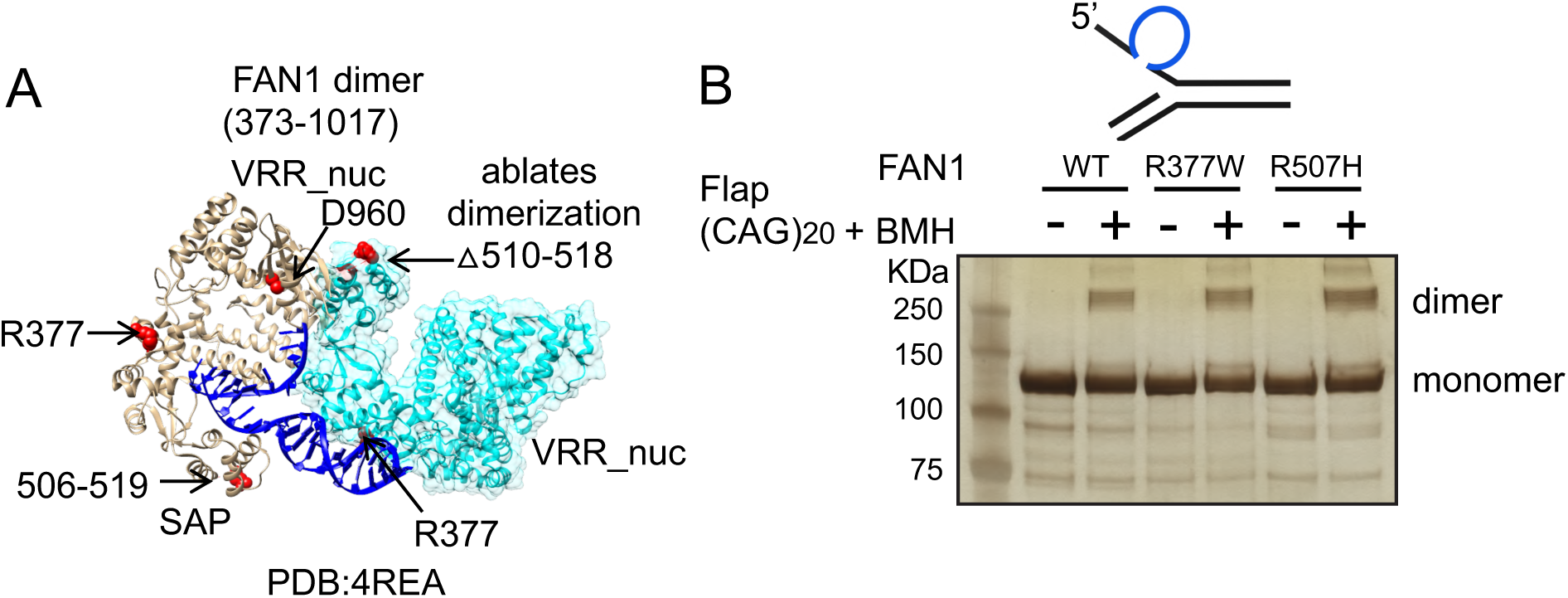
**FAN1 and its variants can dimerize on slipped-DNAs:** (A) Crystal structure of dimeric FAN1 (4REA) drawn by chimera highlighting FAN1 variants by red color (B) FAN1-FAN1 dimerization (BMH cross-linking) assay showing 5’-Flap-(CAG)20 DNA induced dimerization of FAN1^p.WT^, FAN1^p.R507H^ and FAN1^p.R377W^, FAN1 monomer and dimer are indicated in silver-stained gel image.

**Figure S4:**
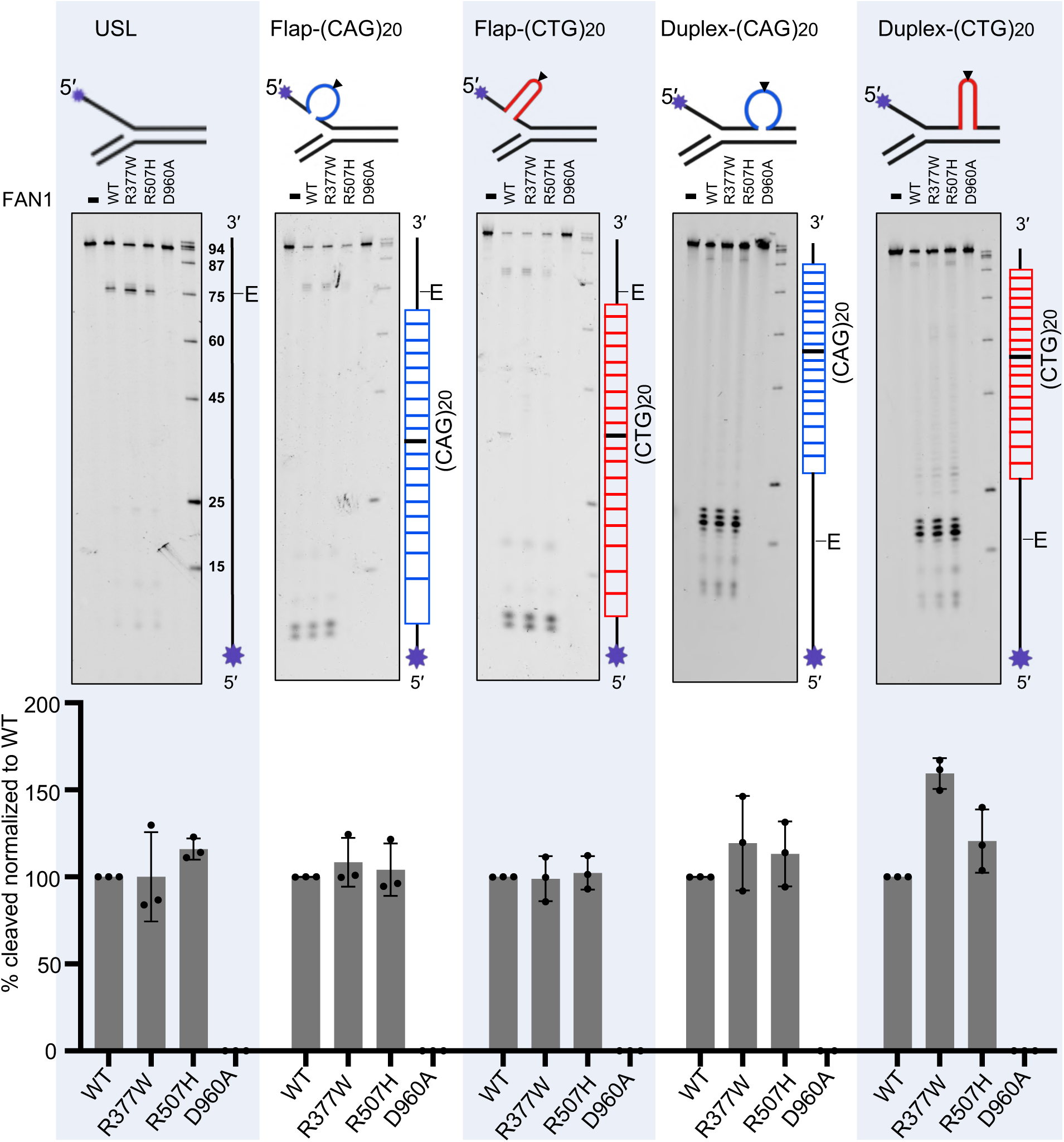
**FAN1 *endo*-nucleolytically cleaves slipped-DNAs similar to canonical repeat-free DNA flaps:** FAN1 and its various forms have *endo*-nuclease activity on 5′ end FAM labelled DNA (label = purple star), which preferentially reveals *endo*-nucleolytic cleavages. 50 nM FAN1^p.WT^ and variants are incubated with 100 nM of DNA (schematic shown above each gel) for 20 minutes and reactions were stopped by denaturing 95% formamide and samples were resolved on 6% denaturing sequencing PAGE at 2000 V for 1 hr. USL (unstructured long flap), flap-bourne Flap-(CAG)20 and Flap-(CTG)20, and duplex-bourne Duplex-(CAG)20, and Duplex-(CTG)20 DNA substrates. All DNAs have 5′- FAM labelled on flap strand (purple star), schematics are shown above each denaturing gel, and labeled strand schematic indicated along the length of the gel, with repeat tract and its center (black line) indicated, “E” denotes elbow region between ds DNA to ssDNA junction. Image obtained on Typhoon fluorescence channels and quantified by ImageQuant. Graphs were plotted by normalizing with the percentage of cleavage for FAN1^p.WT^ by GraphPad Prism 8.2. Representative gels of n=3 independent replicates. Penultimate lane shows absence of nuclease activity with 200 nM FAN1^p.D960A^ for 20 min., indicating that our FAN1 preparations were free of contaminating nucleases, as multiple preparations of the nuclease-dead FAN1^p.D960A^, known to be deficient in *endo*- and *exo-* nucleolytic activities (Smogorzewska et al., 2010; Zhao et al., 2014), were, as expected, consistently nuclease-free. Last lane shows size marker.

**Figure S5:**
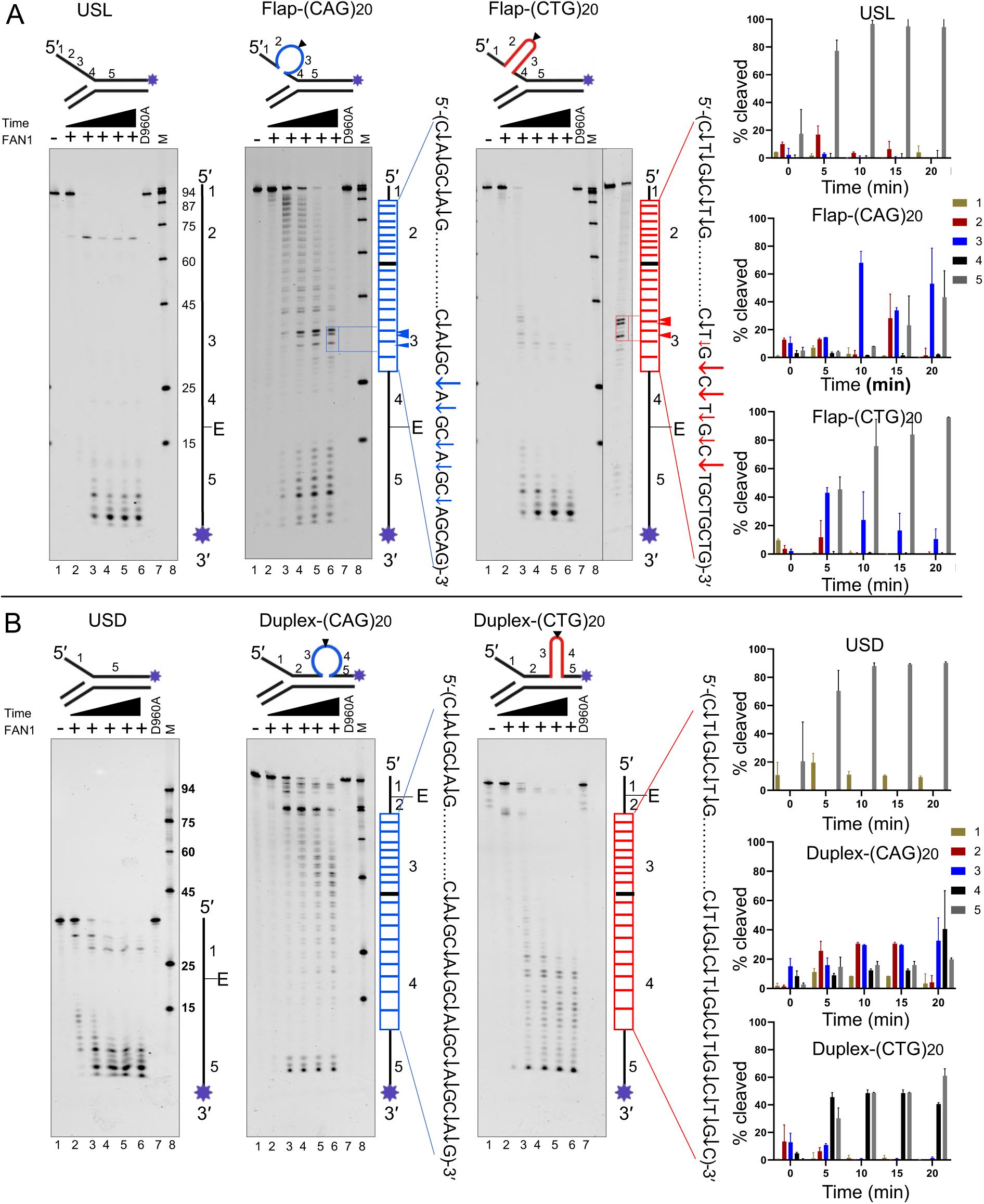
**FAN1 *exo*-nucleolytic cleavage sites depend on slip-out sequence and location (with quantifications):** (A) *Exo*-nucleolytic cleavage patterns were described in Figure 2, quantifications are provided here. USL (unstructured long flap), Flap-(CAG)20, and Flap–(CTG)20 DNA substrates (B) USD, Duplex-(CAG)20 and Duplex-(CTG)20 substrates. All DNAs have 3′-FAM labelled on flap strand (purple star), schematics are shown above each denaturing gel, and labeled strand schematic indicated along the length of the gel, with DNA regions numbered, and repeat tract and its center (black line) indicated, “E” denotes elbow region between ds DNA to ssDNA junction. Following digestions products were denatured and resolved on denaturing sequencing gels. A. Lane 1 indicates, substrate alone and lanes 2-6 show time-dependent digests (0, 5, 10, 15 and 20 min) at 200 nM of human FAN1 and 100 nM DNA substrate (protein: DNA (2:1)). Lane 9 and 10 in rightmost panel of panel A shows substrate alone and nuclease activity of FAN1 at reduced protein concentration (protein: DNA 1:2) for 10 minutes respectively, permitting identification in Lane 10 individual FAN1 cleavage/pause sites on Flap- (CTG)20 DNA substrate. Lane 7 shows absence of nuclease activity with 200 nM FAN1^p.D960A^ for 20 min., indicating that our FAN1 preparations were free of contaminating nucleases, as multiple preparations of the nuclease-dead FAN1^p.D960A^, known to be deficient in *endo*- and *exo-* nucleolytic activities (Smogorzewska et al., 2010; Zhao et al., 2014), were, as expected, consistently nuclease-free. Lane 8 shows size marker. *Exo*-nuclease scissile sites / pause sites were mapped using Maxam-Gilbert chemical DNA sequencing. **For quantification**, the percentage of cleavage (right panel for A and B) was divided in five regions as per schematic and analyzed by ImageQuant, plotted on GraphPad Prism 8.2. Representative gels for n=3 independent replicates.

**Figure S6:**
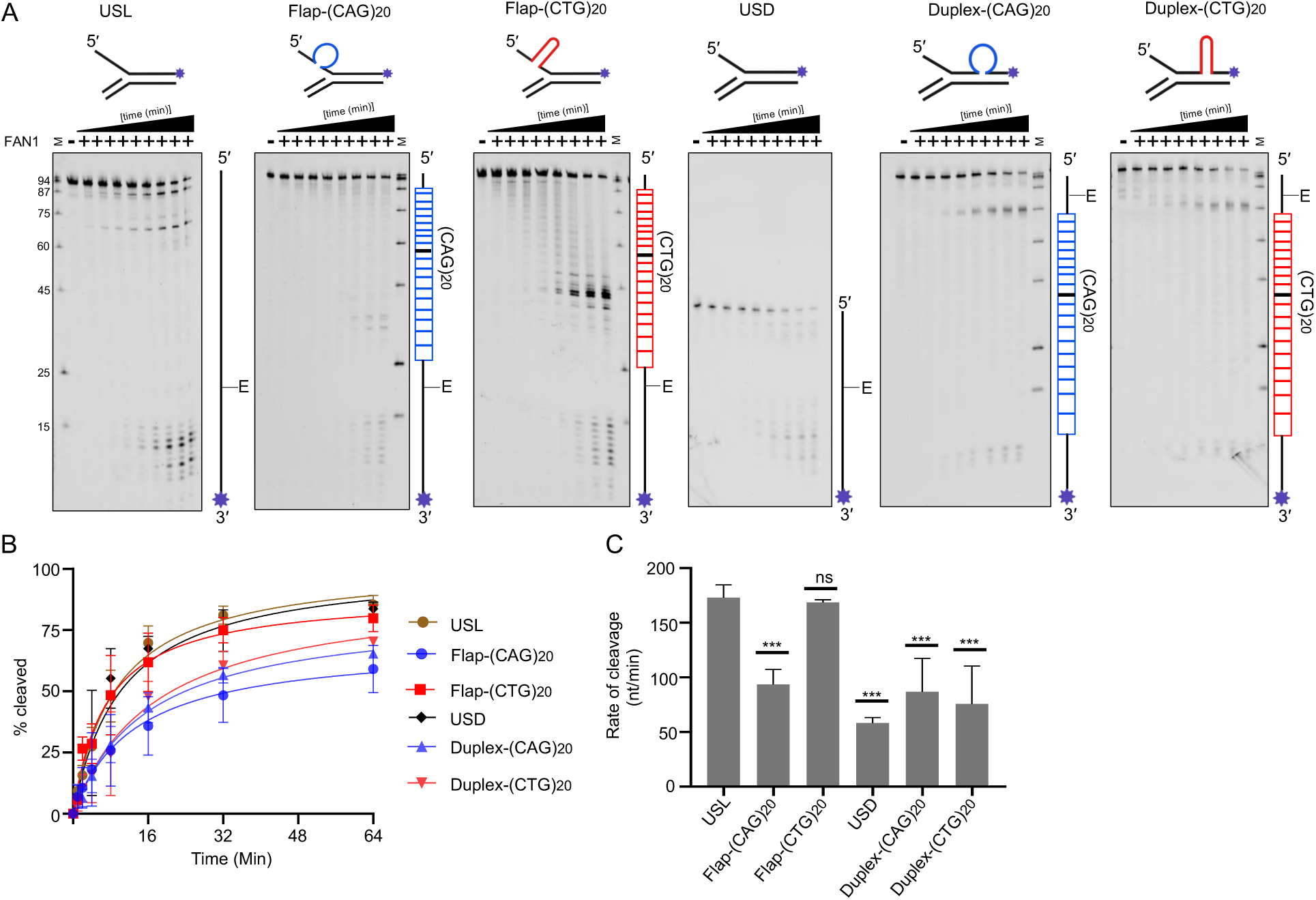
**CAG slip-outs were *exo*-nucleolytically digested significantly slower than CTG slip-outs: A)** All DNAs have 3′-FAM labelled on flap strand (purple star), schematics are shown above each denaturing gel, and labeled strand schematic indicated along the length of the gel, and repeat tract and its center (black line) indicated, “E” denotes elbow region between ds DNA to ssDNA junction. DNAs were incubated with 50 nM of FAN1^p.WT^ with 100 nM of DNA, for 0-128 minutes. Samples were taken at each time interval (0, 2, 4, 8, 16, 32, 64 and 128 minutes), reactions stopped by 95% formamide, samples denatured and resolved on denaturing sequencing PAGE at 2000 V for 1 hr. **B)** Percentage of cleavage is quantified by ImageQuant for each time interval and plotted on GraphPad Prism 8.2. Error bars represent ±SD. Representative gels for n=3 independent replicates. **C)** Rate of cleavage (nucleotide/minutes) is calculated as described in (Subramanian et al., 2003) and statistically assessed as per (Kurita et al., 2009), and plotted on GraphPad Prism (Representative gels for n=3 independent replicates). Error bars represent ±SD. Statistical significance was calculated by Welch’s *t* Test, ***P<0.001, ns (not significant).

**Figure S7.**
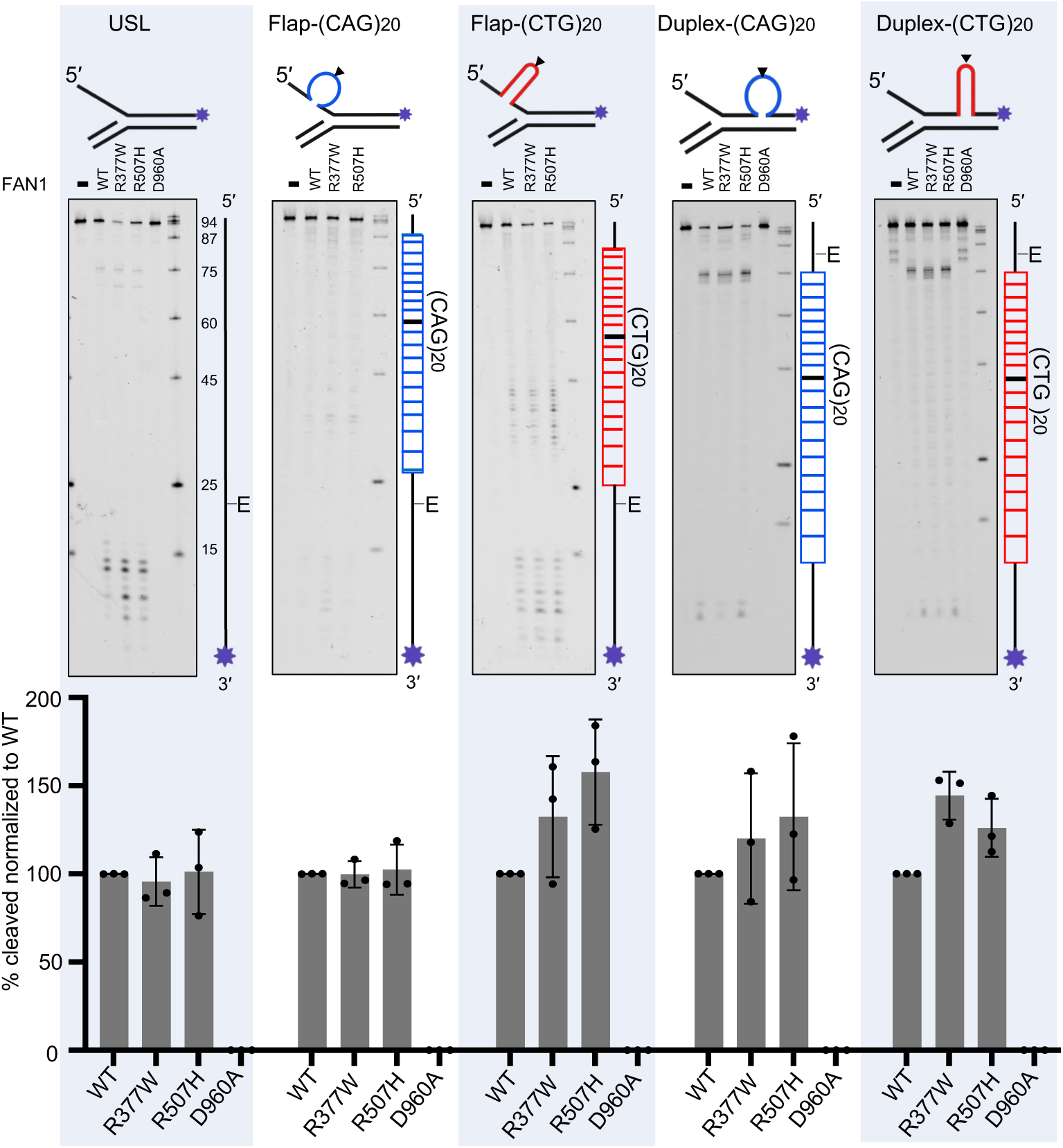
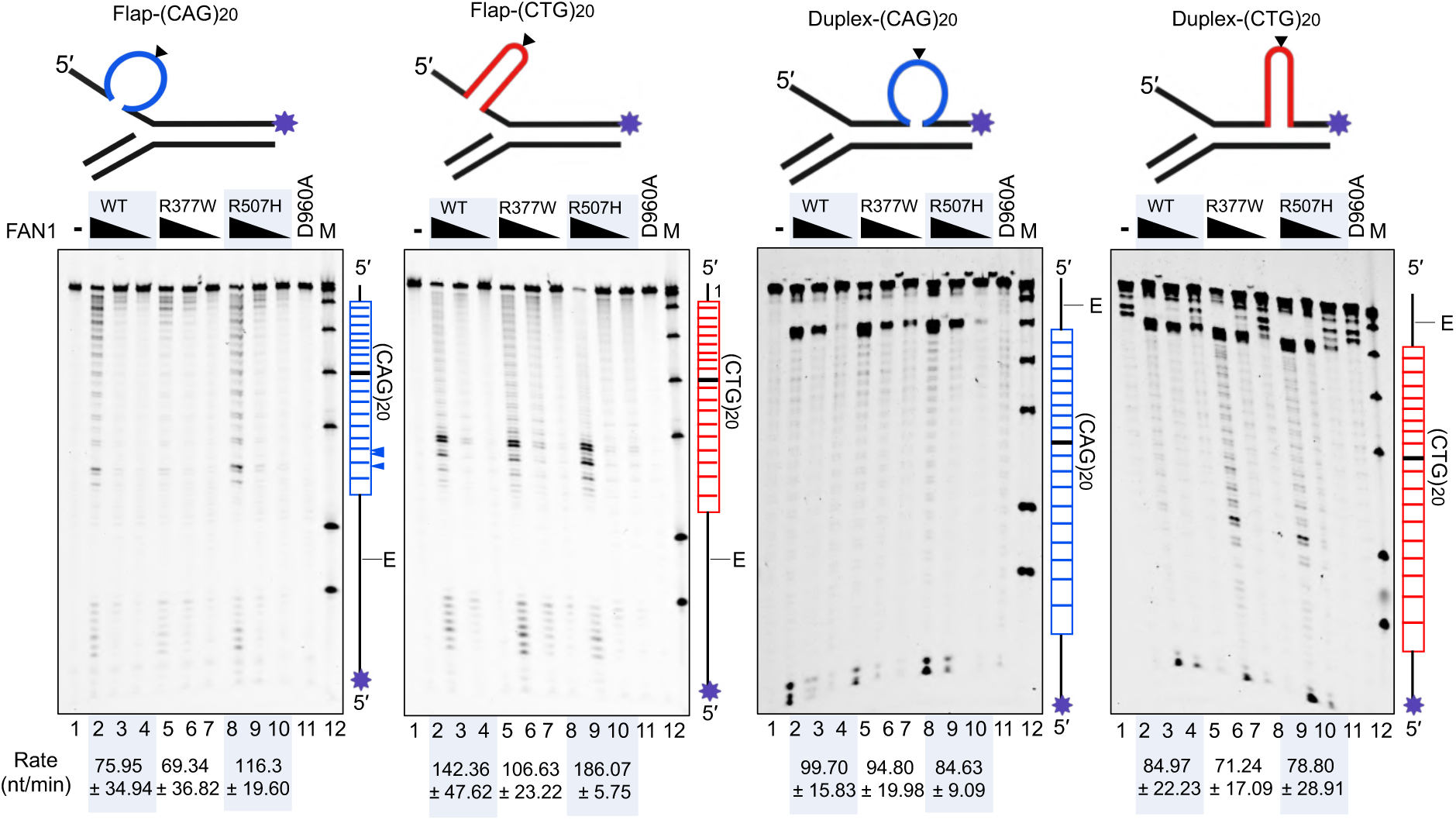
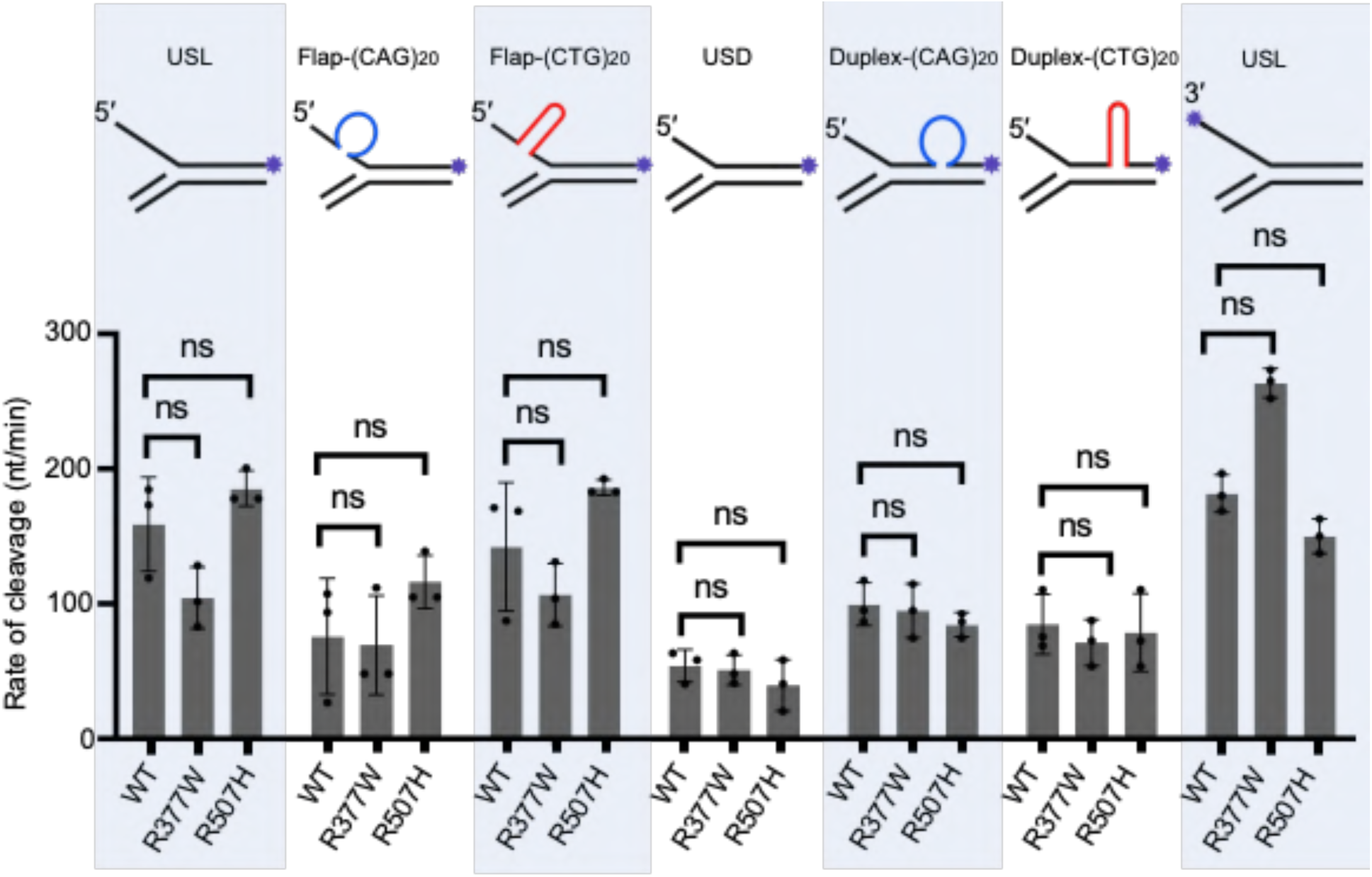
**A: FAN1 variants show similar *exo*-nucleolytic cleavage of slipped-DNAs similar to repeat-free DNA flap**s: FAN1 and its various form are tested for *exo-*nuclease activity on 3′- FAM labelled on flap strand (purple star), schematics are shown above each denaturing gel, and labeled strand schematic indicated along the length of the gel, and repeat tract and its center (black line) indicated, “E” denotes elbow region between ds DNA to ssDNA junction. 50 nM FAN1^p.WT^ and variants are incubated with 100 nM of respective DNA (shown on the top of each gel) for 20 minutes and reactions were stopped by 95% formamide and samples were resolved on 6% denaturation PAGE at 2000 V for 1 hr. Image was taken on Typhoon fluorescence channels and analyzed by ImageQuant. The graphs were plotted by normalizing percentage of cleavage for FAN1^p.WT^ by GraphPad Prism 8.2 (Representative gels for n=3 independent replicates). Penultimate and last lane shows absence of contaminating nuclease activity (FAN1^p.D960A^ for 20 min.) and size marker, respectively. **B**: **FAN1 *exo*-nucleolytically digests slip-outs in a dose-dependent manner:** FAN1 and its various form are tested for dose effect of *exo-*nuclease activity on 3′-FAM labelled on flap strand (purple star), schematics are shown above each denaturing gel, and labeled strand schematic indicated along the length of the gel, and repeat tract and its center (black line) indicated, “E” denotes elbow region between ds DNA to ssDNA junction. Lane 1 shows substrate alone, lanes 2-4 show concentration dependent cleavage (100, 50, and 25 nM) of FAN1^p.WT^ and lane 5-7 show FAN1^p.R377W^ and lane 8-10 shows FAN1^p.R507H^ on 100 nM DNA substrate (Schematic of DNA is shown on top of each gel). Reactions were stopped by adding 95% formamide and samples were resolved on 6% denaturing PAGE at 2000 V for 1hr. Images were taken in Typhoon fluorescence channel and analyzed by ImageQuant (Representative gels for n=3 independent replicates). Quantified digestion rates are indicated. Penultimate and last lane shows absence of contaminating nuclease activity (FAN1^p.D960A^ for 20 min.) and size marker, respectively. **C**: **FAN1 and its forms do not show significant differences in rates of *exo*- nucleolytic cleavage:** FAN1 and its forms were tested for rate of cleavage in time dependent manner. The 3′-FAM labelled DNA substrates (schematic of each DNA substrate, label = purple star, is shown above each graph) is incubated with 50 nM of FAN1^p.WT^ with 100 nM of DNA, for 0-128 minutes. Samples were taken at each interval (0, 2, 4, 8, 16, 32, 64, and 128 minutes) and reactions stopped by 95% formamide, samples were resolved on denaturing PAGE at 2000 V for 1 hr (gel images are not shown). B) Rate of cleavage (nucleotide/minutes) is calculated as described in (Subramanian et al., 2003) and statistically assessed as per (Kurita et al., 2009), and plotted on GraphPad Prism. Experiments were done in triplicate. Error bars represent ±SD. Statistical significance was calculated by Welch’s *t* test, ns (not significant). **Figure S8: Schematic of mapped cleavage sites on slip-outs:** Cleavage sites were mapped by Maxam-Gilbert chemical sequencing and summarized in the schematic

**Figure S8:**
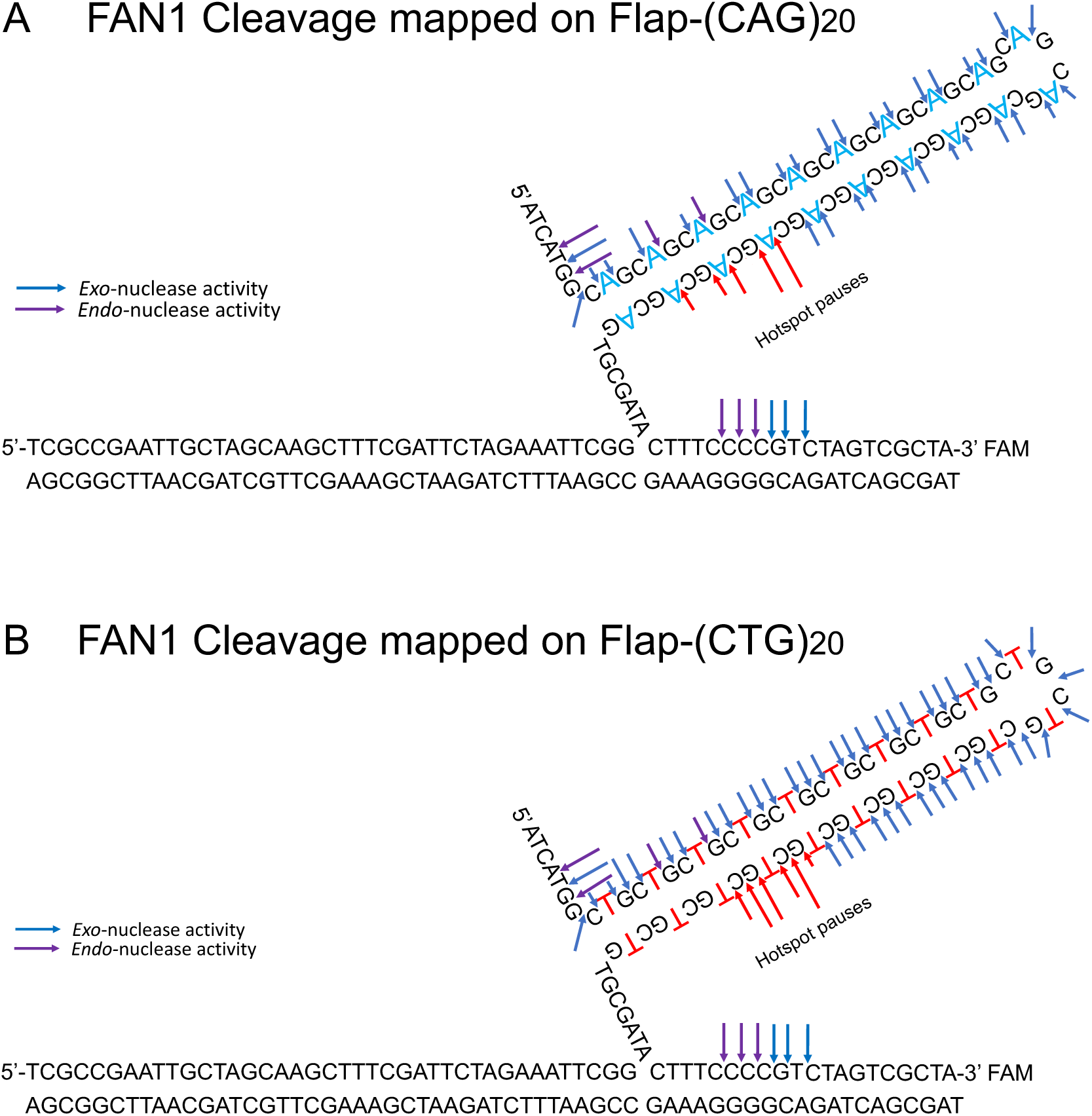
**Schematic of mapped cleavage sites on slip-outs:** Cleavage sites were mapped by Maxam-Gilbert chemical sequencing and summarized in the schematic

**Figure S9:**
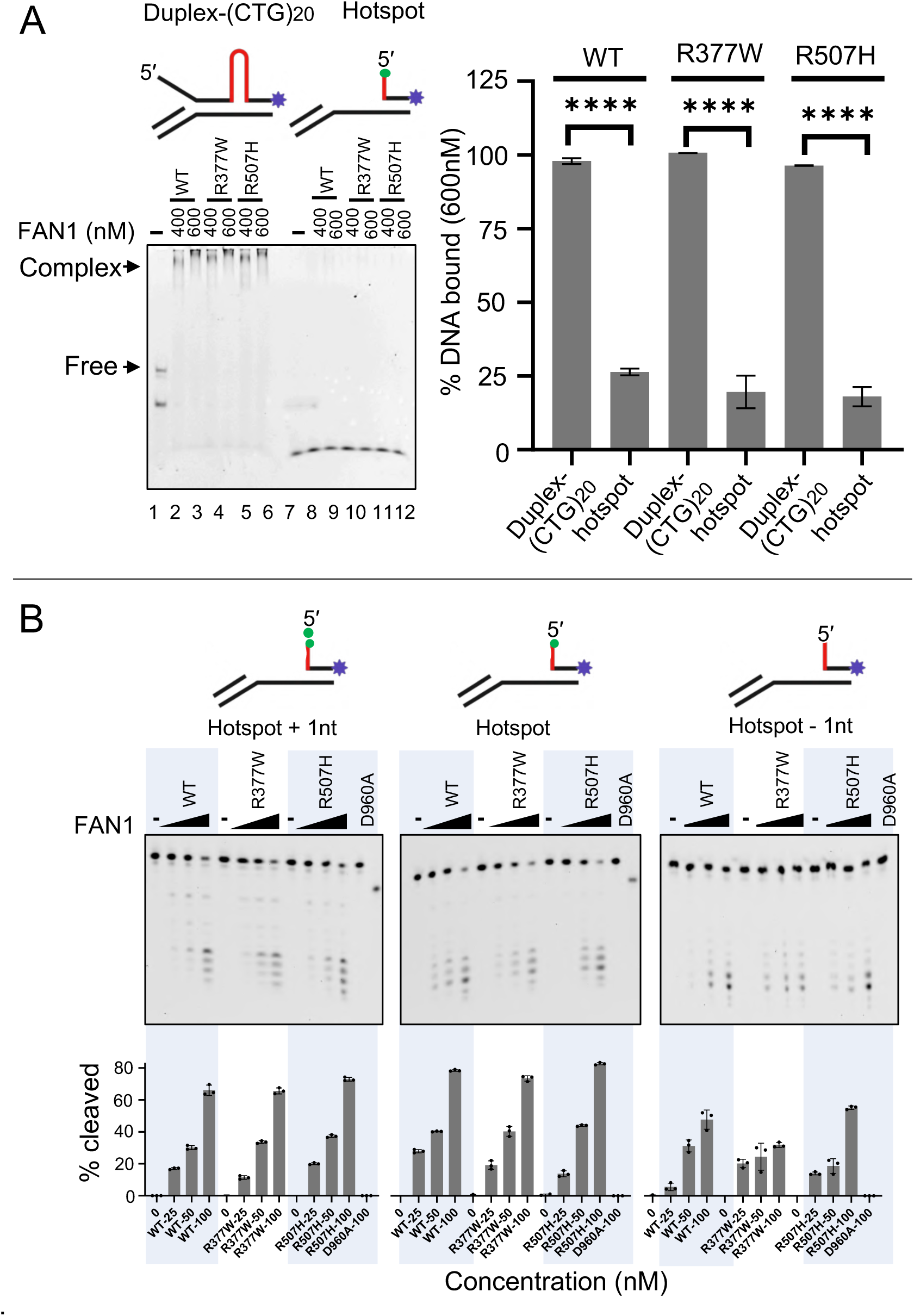
**FAN1p^.WT^ and variants DNA-binding and *exo*-nuclease on CTG hotspot DNA substrates: A)** FAN1^p.WT^ and variants bind very poorly to the terminal “hotspot” DNA digestion product arising from the precursor Duplex-(CTG)20 substrate even at very high protein concentration, while the precursor Duplex-(CTG)20 substrate is an excellent substrate. This “hotspot” substrate has gap of 10 nucleotides and a Flap of 7 nucleotide flap at the indicated stoichiometric concentrations. At left the percentage DNA-binding of FAN1 and its forms were compared with DNA binding at “hotspot” DNA, shows significant reduction (****p>0.0001, Welch’s *t* test). Experiments were done in triplicate. Error bars represent ±SD. **B)** *Exo*-nuclease activity was accessed in FAN1 concentration dependent manner on the “hotspot” product arising from the Duplex-(CTG)20, and Hotspot+1 nucleotide and Hotspot-1 nucleotide substrate. Reactions were stopped, denatured and resolved on denaturing PAGE. Individual bands are quantitated and graphed as in bottom of each gel. Successive incisions changed for Hotspot+1 nucleotide and Hotspot to Hotspot-1 nucleotide, suggest length dependent cleavage and cleaves in duplex region.

**Figure S10:**
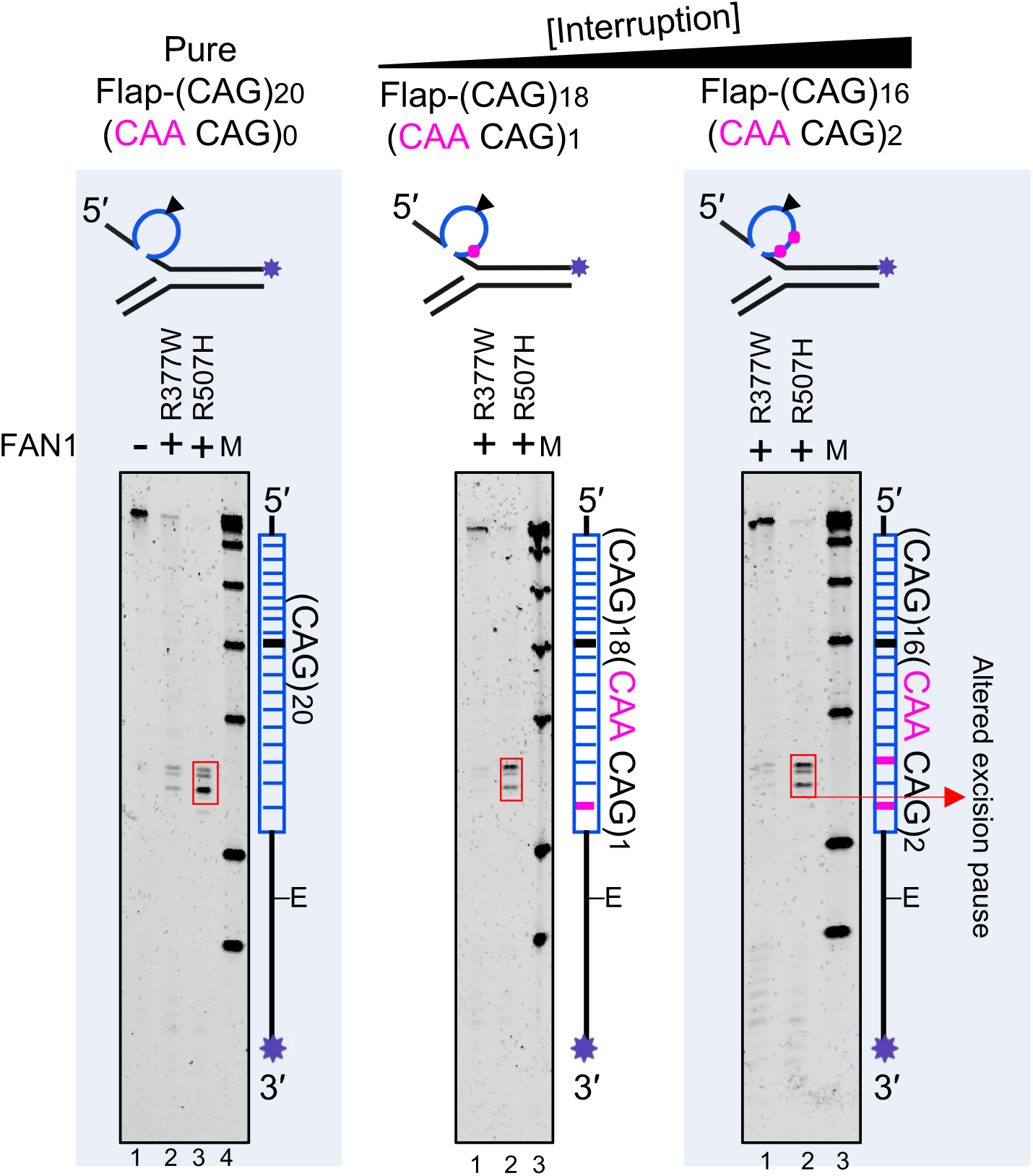
**Interruption changes cleavage site location in hotspot pause:** 200 nM of FAN1 and its forms were incubated with 3′-FAM labelled (flap strand) 100 nM of pure CAG repeat, and CAA-interrupted CAG repeat DNA (schematic is shown on the top of each gel, interruptions are violet dots/bars) for 20 minutes, reactions were stopped by 95% formamide and resolved on 6% denaturing gel and cleavage products were observed by Typhoon fluorescence channel.

**Figure S11:**
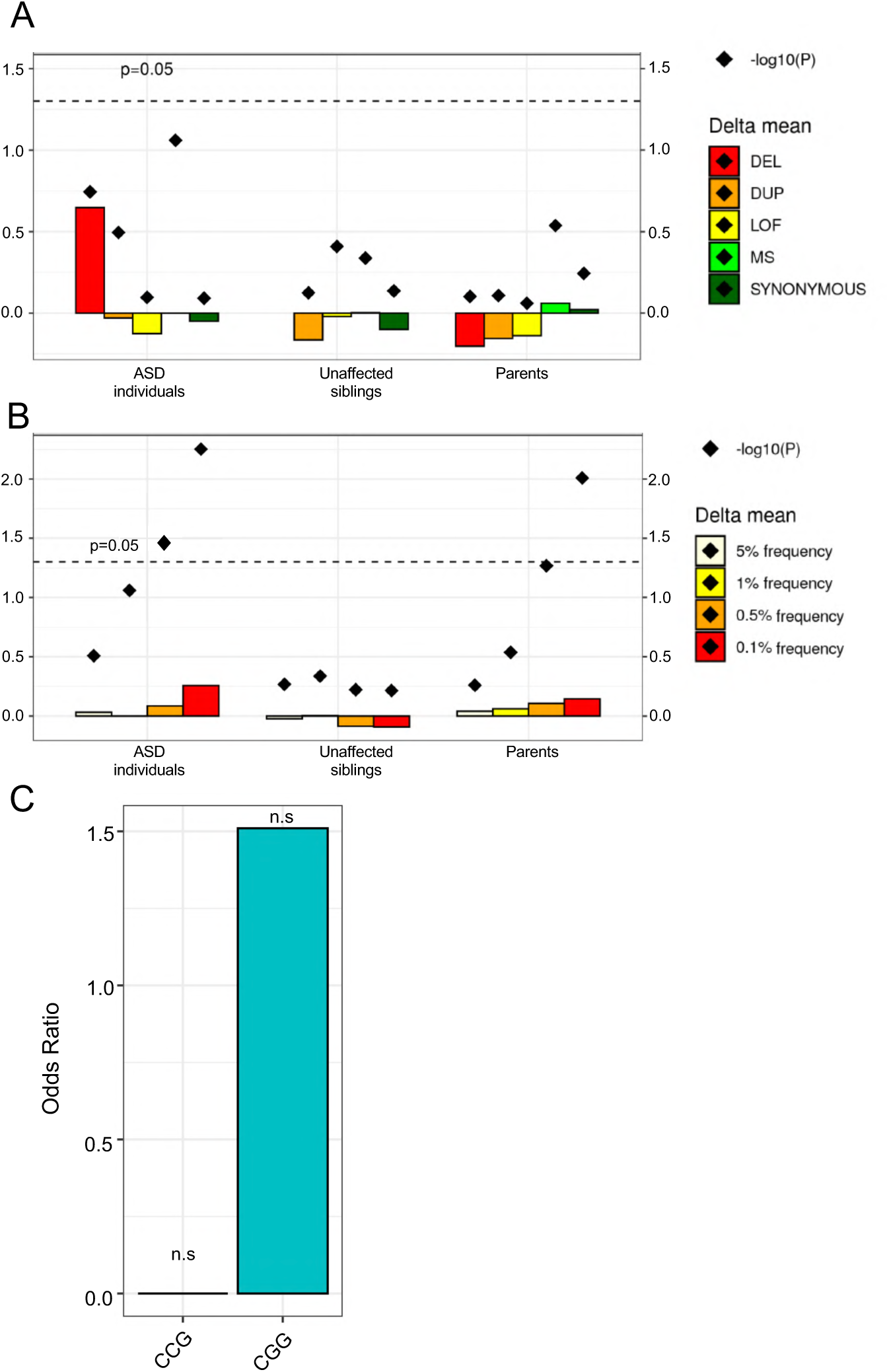
**Tandem repeat count of individuals with genetic variants in *FAN1*. A)** Mean lengths of tandem repeat count in each population (children with ASD, n=4,969; children without ASD, n=1,913; and parents, n=7,945) are compared between individuals with and without genetic variants in *FAN1* using Wilcoxon’s test. Genetic variants investigated include **A)** rare variants (frequency <1%) and **B)** missense variants with various frequency (5%, 1%, 0.5%, 0.1%). Tandem repeat count was based on the sum of in-repeat read counts for CGG (/CCG), CAG (/CTG) or AT as calculated by ExpansionHunter Denovo. Positive value of delta_mean represents longer mean lengths in genetic variant carriers. DEL: copy number loss; DUP: copy number gain; LOF: loss of function variants; MS: missense variants. **C)** Association between the presence of rare (MAF <0.5%) missense FAN1 variants and CCG or CGG repeat expansions. Fisher’s exact test was used to determine the likelihood (odds ratio) of rare (MAF <0.5%) FAN1 missense variants contributing specifically to CCG and CGG repeat expansions, as determined by anchored in-repeat reads from ExpansionHunter Denovo. Non-significant results are denoted by ‘ns’.

**Figure S12:**
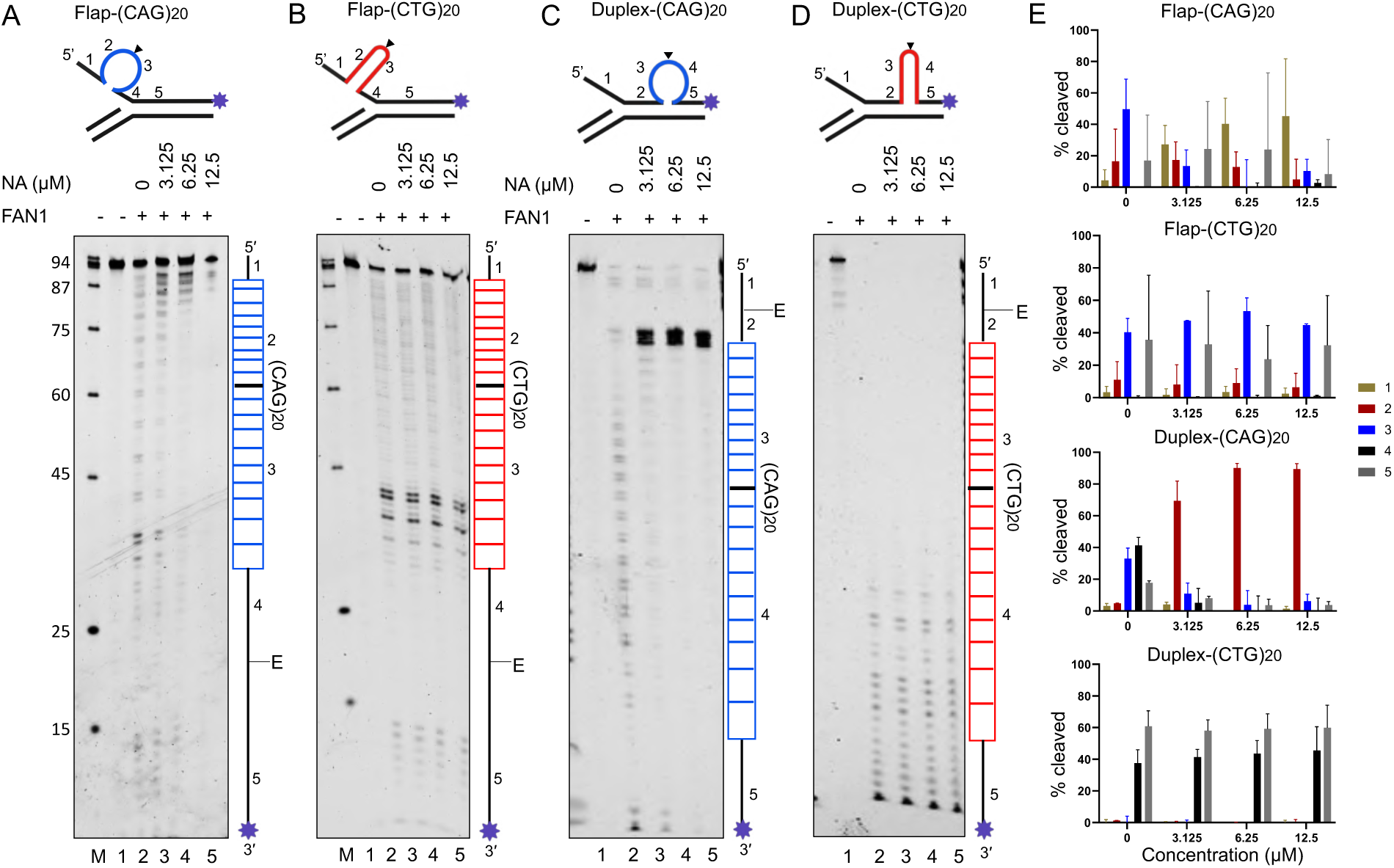
**NA alters FAN1 *exo-*nuclease activity on (CAG)20 slip-outs but not (CTG)20 slip-outs**: Schematics of substrates used are shown on top panel **A)** Flap- (CAG)20 **B)** Flap-(CTG)20 **C)** Duplex-(CAG)20 and **D)** and Duplex-(CTG)20. All DNAs have 3′-FAM labelled on flap strand (purple star), schematics are shown above each denaturing gel, and labeled strand schematic indicated along the length of the gel, with DNA regions numbered, and repeat tract and its center (black line) indicated, “E” denotes elbow region between ds DNA to ssDNA junction. 100 nM of each DNA was incubated with increasing concentrations of NA (3.125 μM, 6.25 μM, and 12.5 μM) and then treated with 100 nM of FAN1. *Exo*-nuclease reactions were stopped by adding 95% formamide EDTA and resolved on 6% denaturing sequencing gel at 2000 V for 1hr. **E)** % of cleavage were analyzed for each numbered region by ImageQuant and plotted by GraphPad Prism. Representative gels of n=3 replicates. Thus, NA does not block FAN1s *exo*-nucleolytic activity per se, as the digestion of the CTG slip-out, which cannot be bound by NA, was unaffected by NA. Experiments were done in triplicate. Error bars represent ±SD.

**Figure S13:**
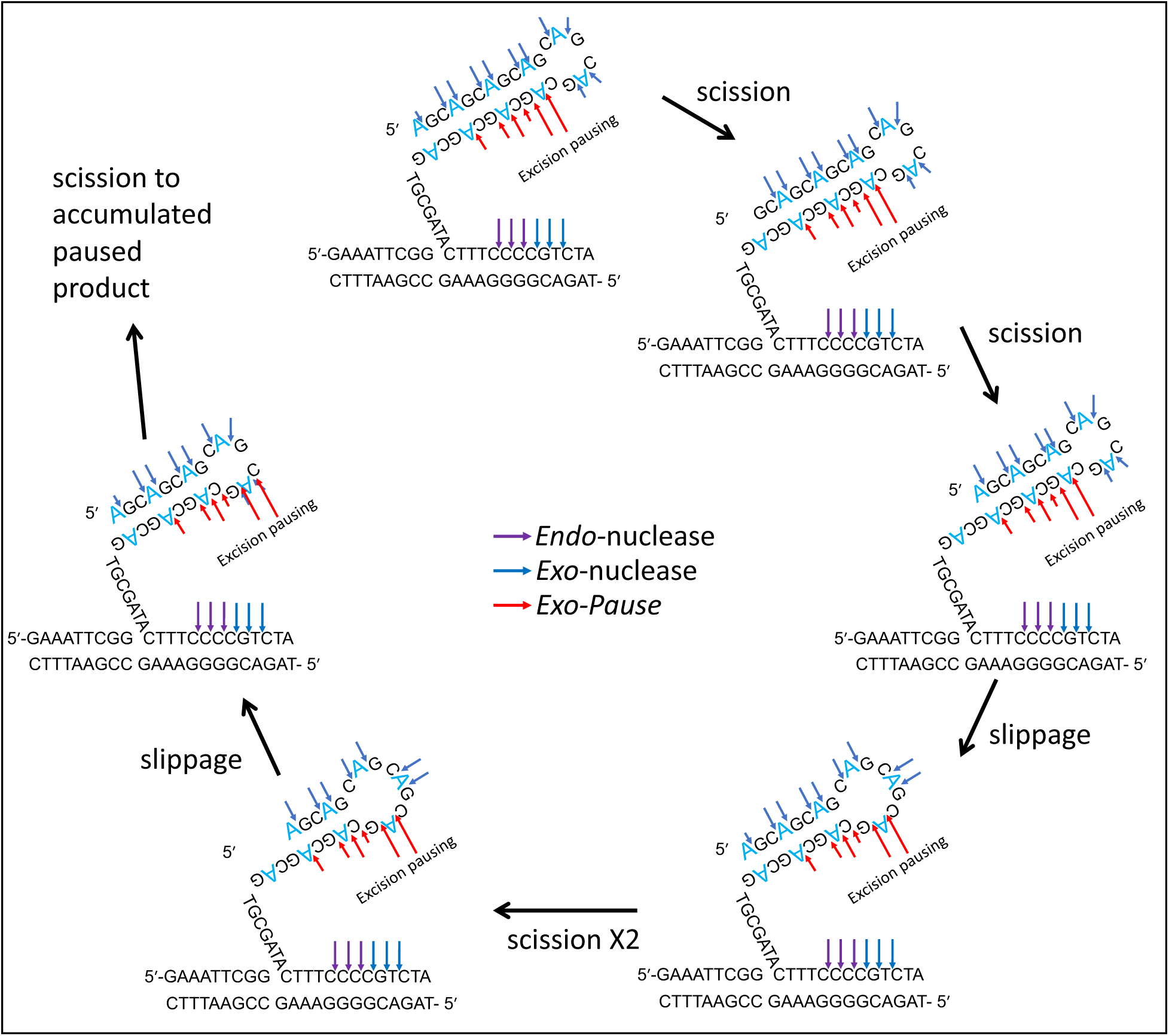
Strand slippage between iterative FAN1 excision/pause steps (see text for details). The length of the red arrows reflects the degree of *exo*-nucleolytic pausing.

**Table S1:**
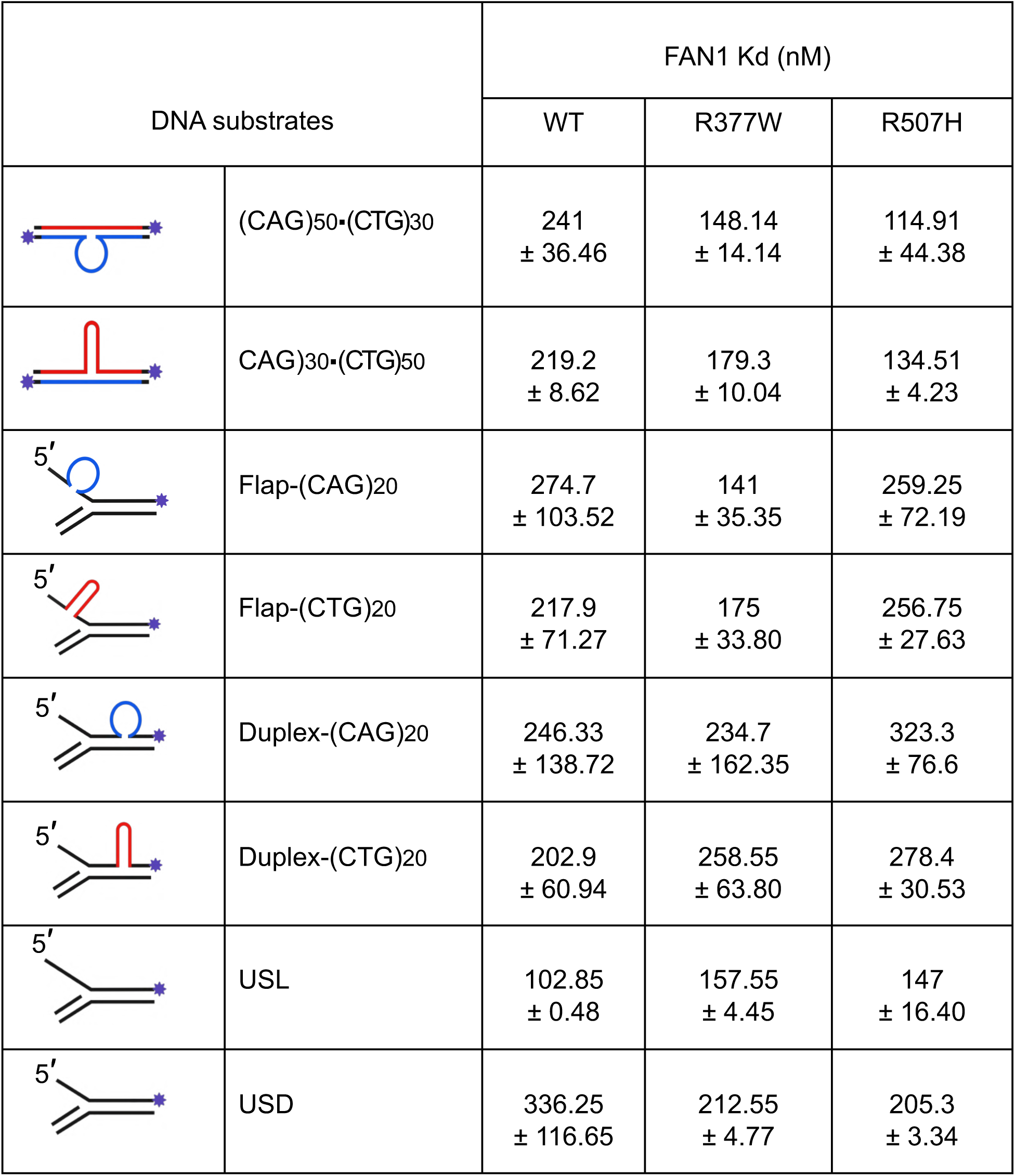
Binding affinity (Kd) summary

| DNA substrates |  | FAN1 Kd (nM) |  |  |
| --- | --- | --- | --- | --- |
|  |  | WT | R377W | R507H |
|  | (CAG)<sub>50</sub>•(CTG)<sub>30</sub> | 241<br>± 36.46 | 148.14<br>± 14.14 | 114.91<br>± 44.38 |
|  | CAG)<sub>30</sub>•(CTG)<sub>50</sub> | 219.2<br>± 8.62 | 179.3<br>± 10.04 | 134.51<br>± 4.23 |
|  | Flap-(CAG)<sub>20</sub> | 274.7<br>± 103.52 | 141<br>± 35.35 | 259.25<br>± 72.19 |
|  | Flap-(CTG)<sub>20</sub> | 217.9<br>± 71.27 | 175<br>± 33.80 | 256.75<br>± 27.63 |
|  | Duplex-(CAG)<sub>20</sub> | 246.33<br>± 138.72 | 234.7<br>± 162.35 | 323.3<br>± 76.6 |
|  | Duplex-(CTG)<sub>20</sub> | 202.9<br>± 60.94 | 258.55<br>± 63.80 | 278.4<br>± 30.53 |
|  | USL | 102.85<br>± 0.48 | 157.55<br>± 4.45 | 147<br>± 16.40 |
|  | USD | 336.25<br>± 116.65 | 212.55<br>± 4.77 | 205.3<br>± 3.34 |

**Table S2:**
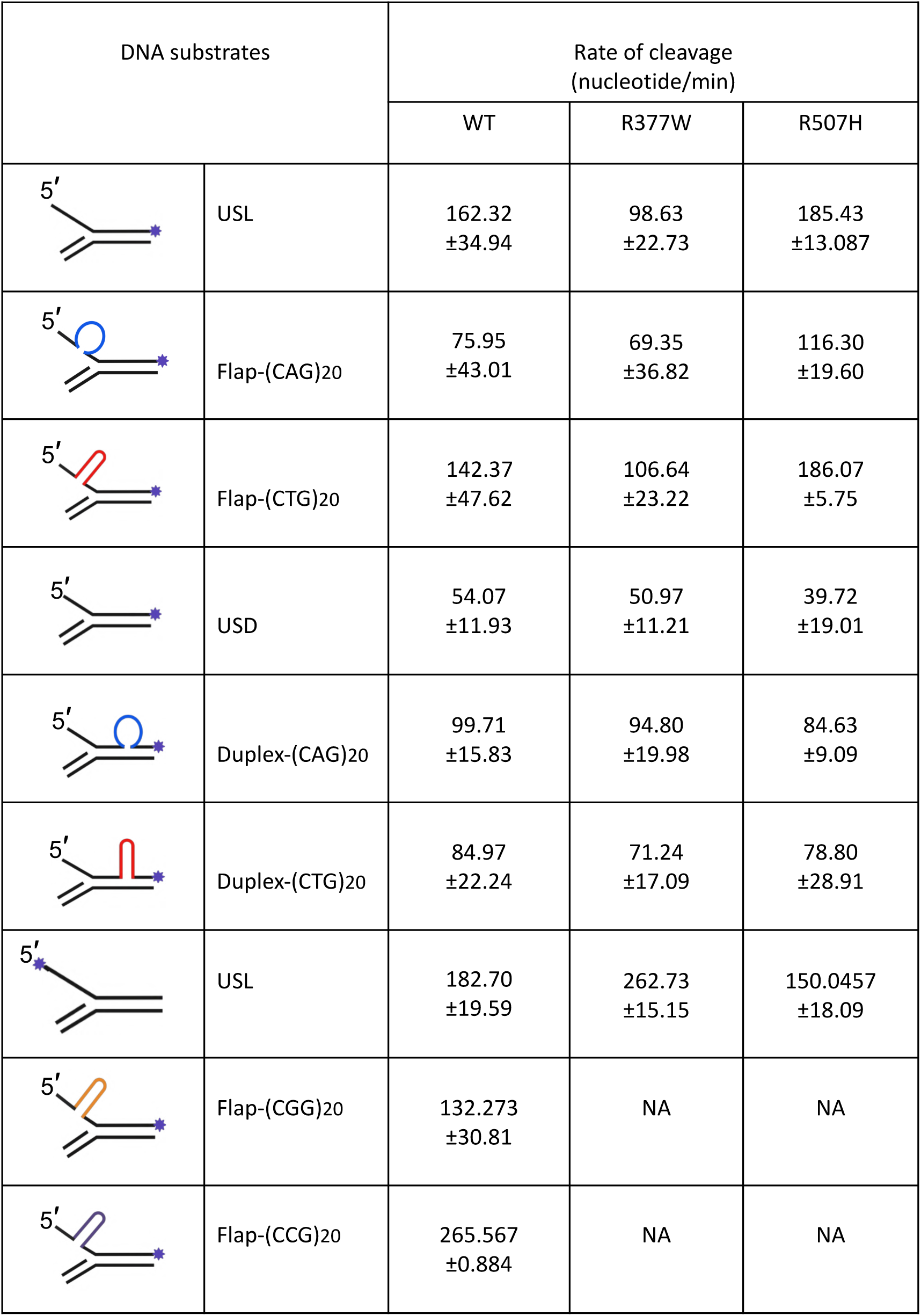
Exo-nucleolytic rate cleavage summary

| DNA substrates |  | Rate of cleavage<br>(nucleotide/min) |  |  |
| --- | --- | --- | --- | --- |
|  |  | WT | R377W | R507H |
|  | USL | 162.32<br>±34.94 | 98.63<br>±22.73 | 185.43<br>±13.087 |
|  | Flap-(CAG)20 | 75.95<br>±43.01 | 69.35<br>±36.82 | 116.30<br>±19.60 |
|  | Flap-(CTG)20 | 142.37<br>±47.62 | 106.64<br>±23.22 | 186.07<br>±5.75 |
|  | USD | 54.07<br>±11.93 | 50.97<br>±11.21 | 39.72<br>±19.01 |
|  | Duplex-(CAG)20 | 99.71<br>±15.83 | 94.80<br>±19.98 | 84.63<br>±9.09 |
|  | Duplex-(CTG)20 | 84.97<br>±22.24 | 71.24<br>±17.09 | 78.80<br>±28.91 |
|  | USL | 182.70<br>±19.59 | 262.73<br>±15.15 | 150.0457<br>±18.09 |
|  | Flap-(CGG)20 | 132.273<br>±30.81 | NA | NA |
|  | Flap-(CCG)20 | 265.567<br>±0.884 | NA | NA |

Note: Please see Table S3 at the end of the draft.

Note: Please see Table S3 at the end of the draft.

**Table S4:**
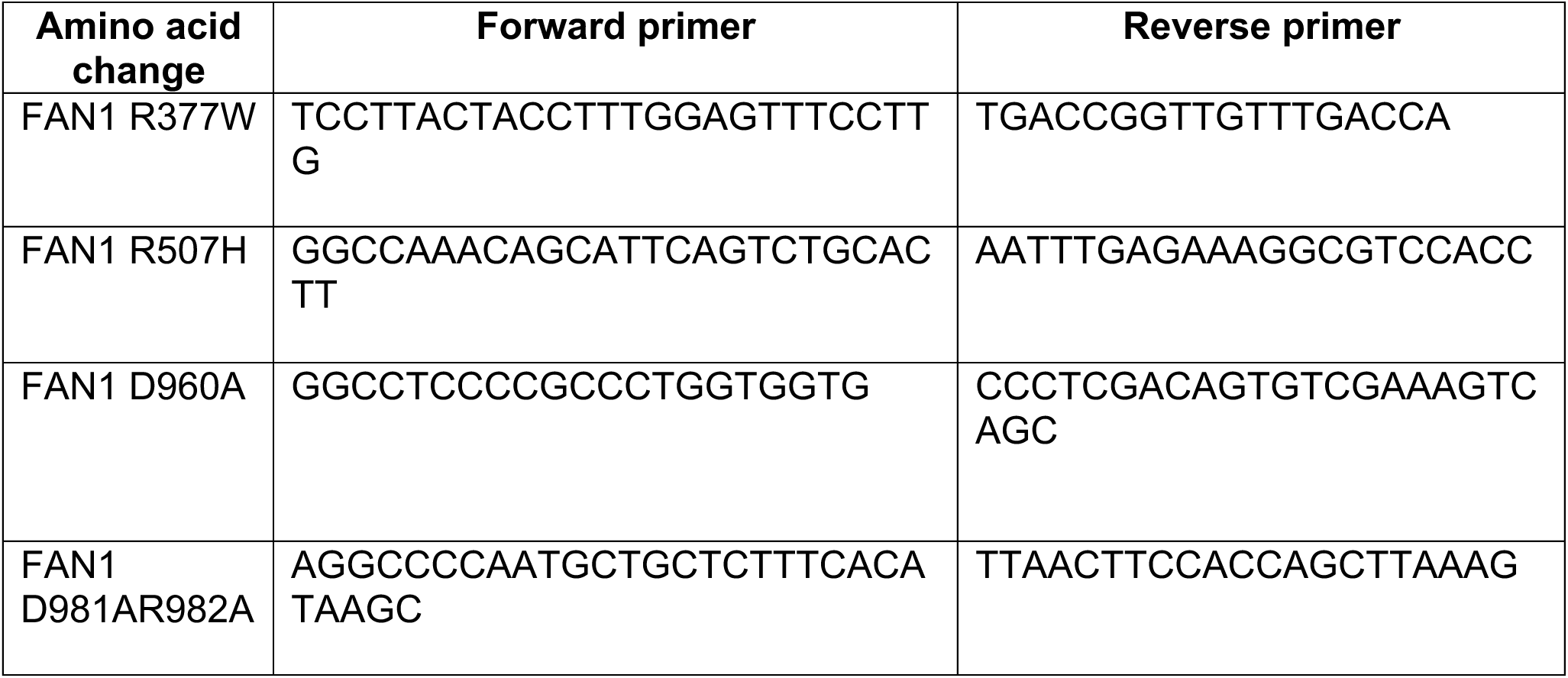
Primers used for cloning

**Table S5:**
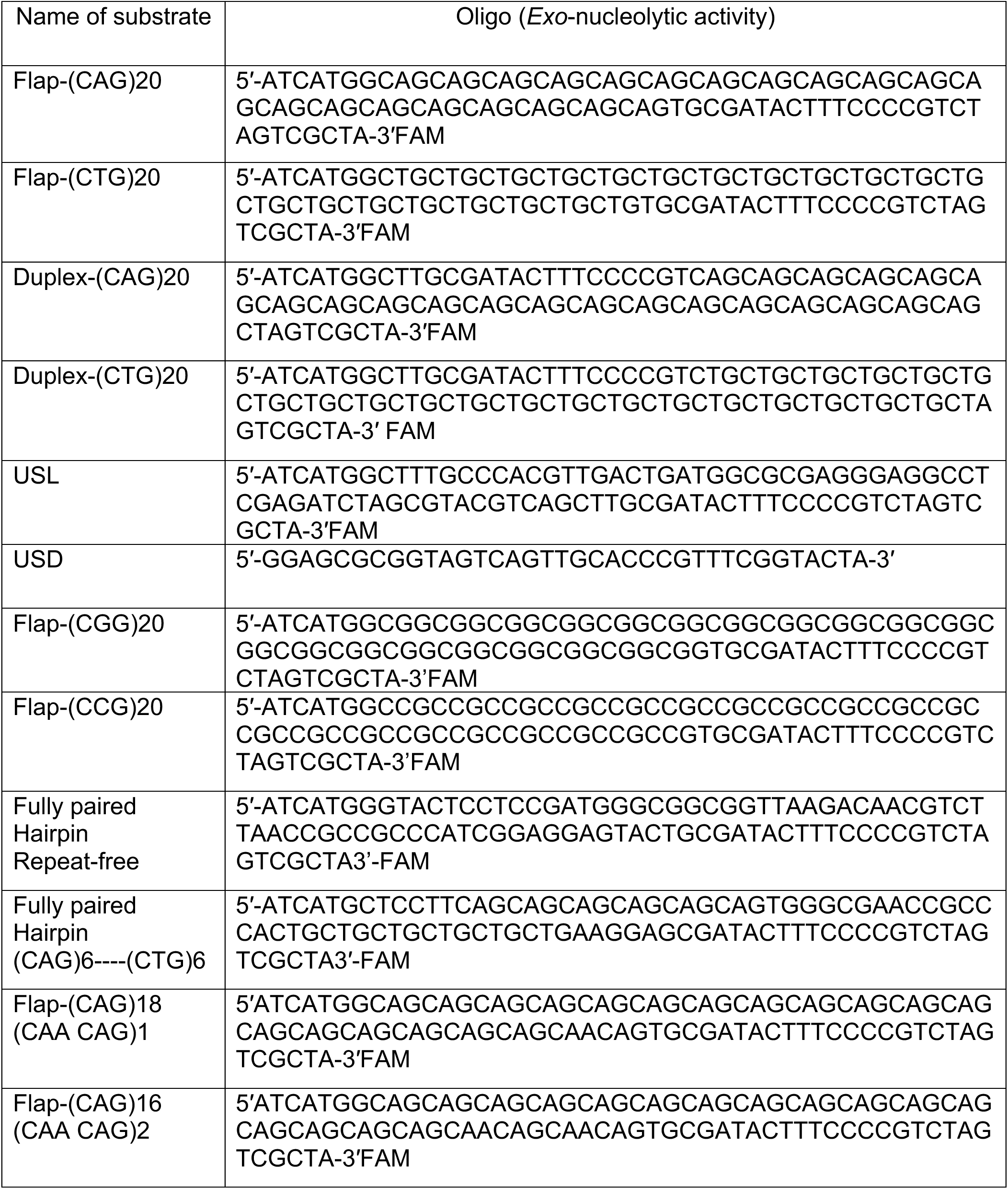
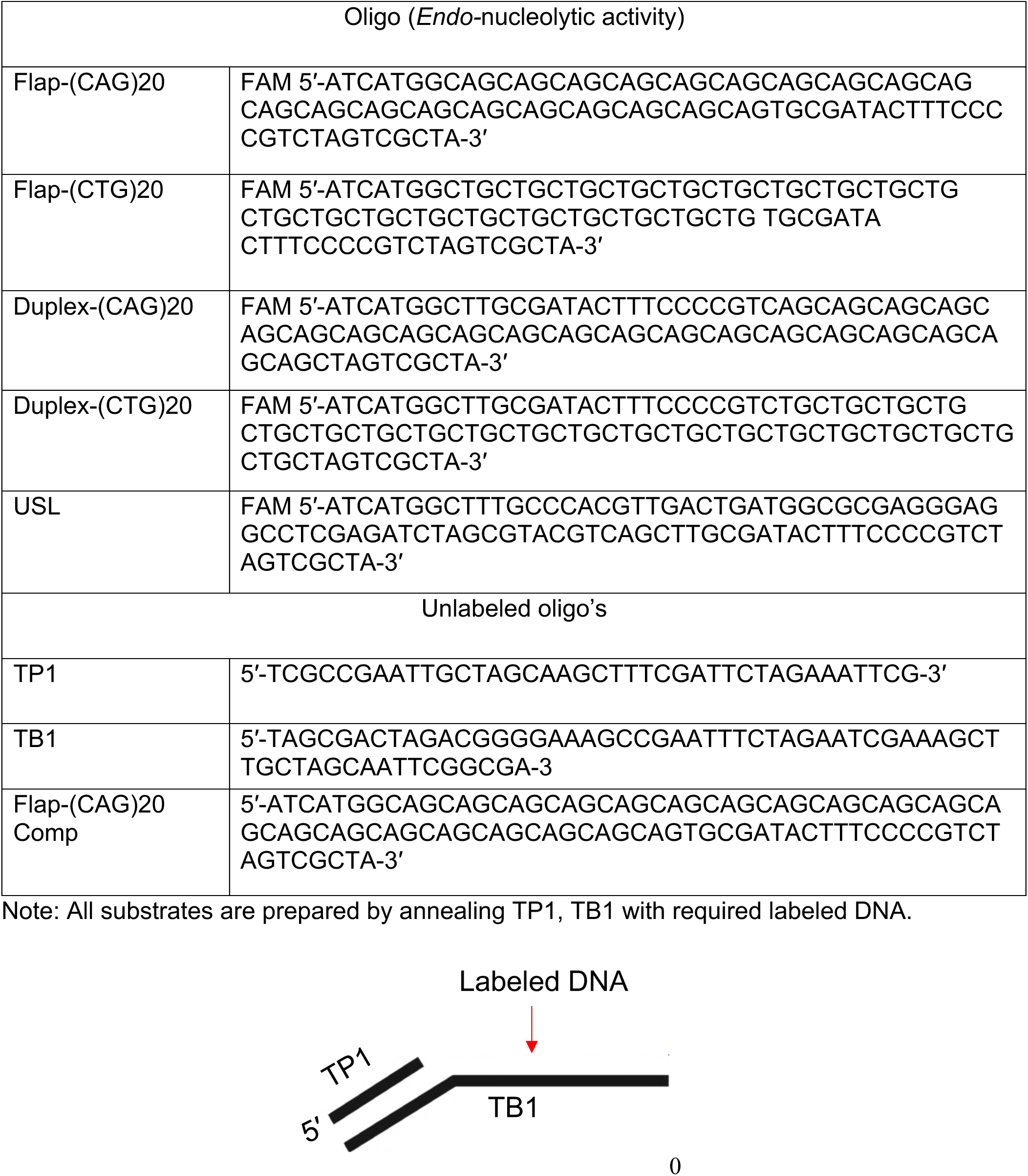
Oligo’s for DNA substrates

**Table S3.**
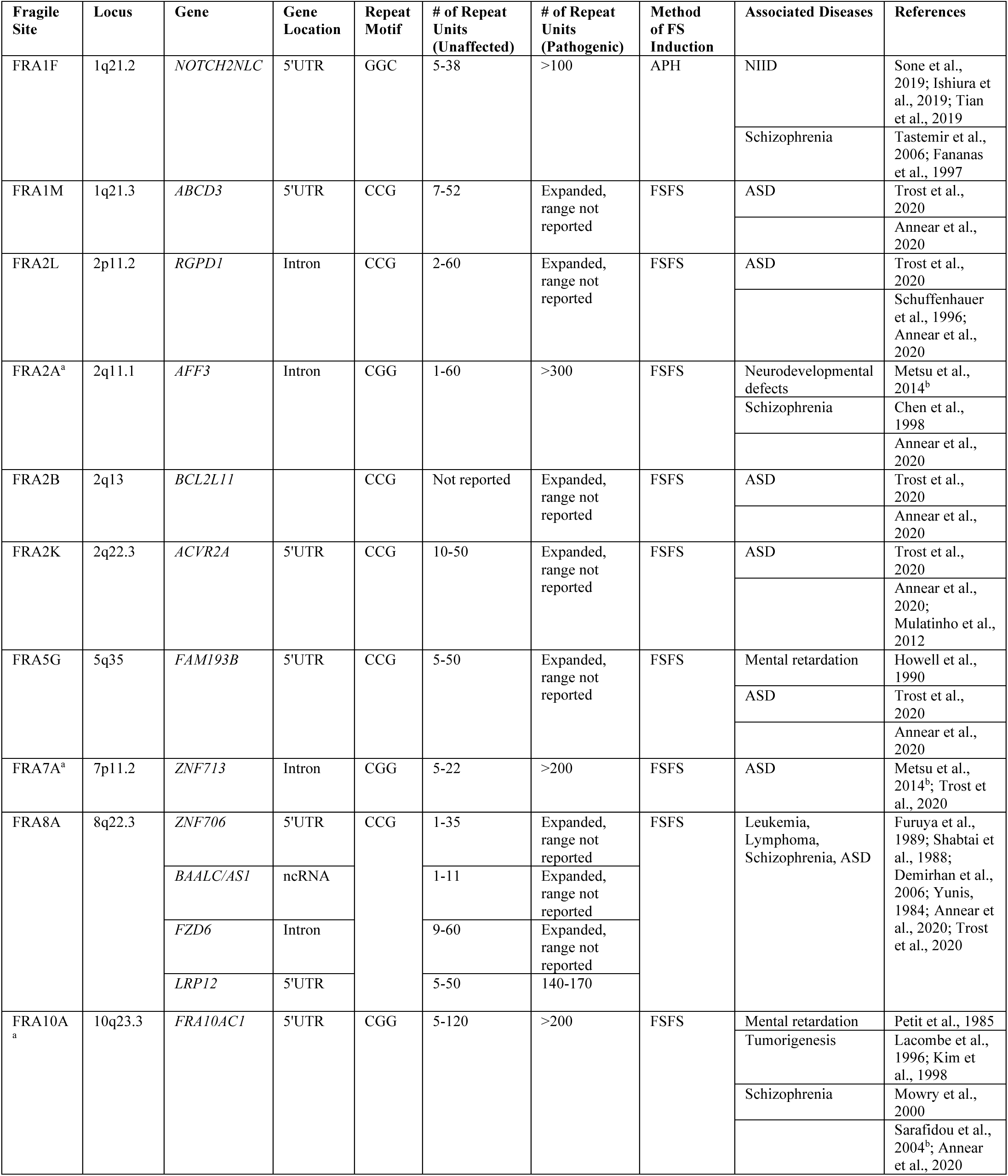
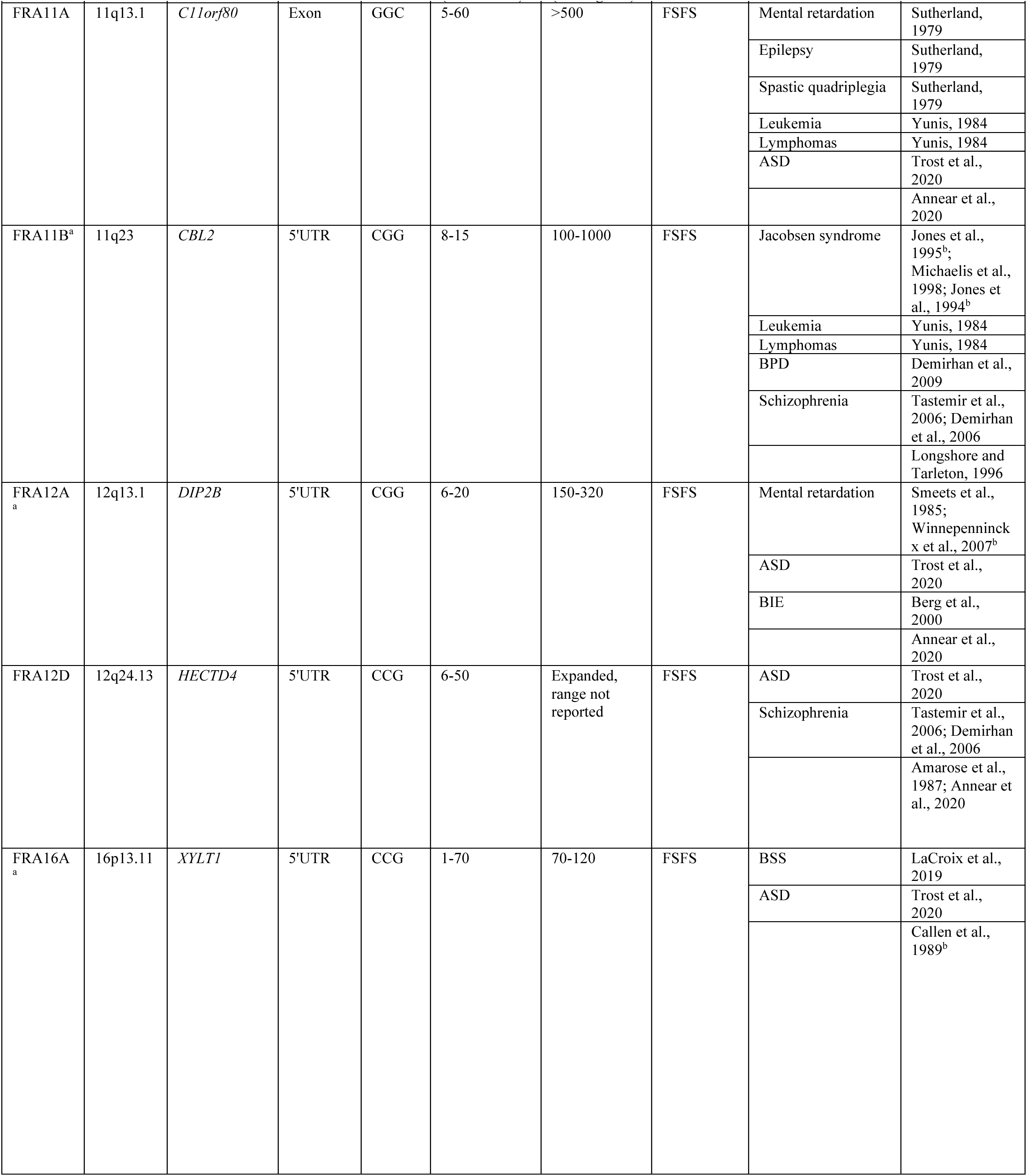
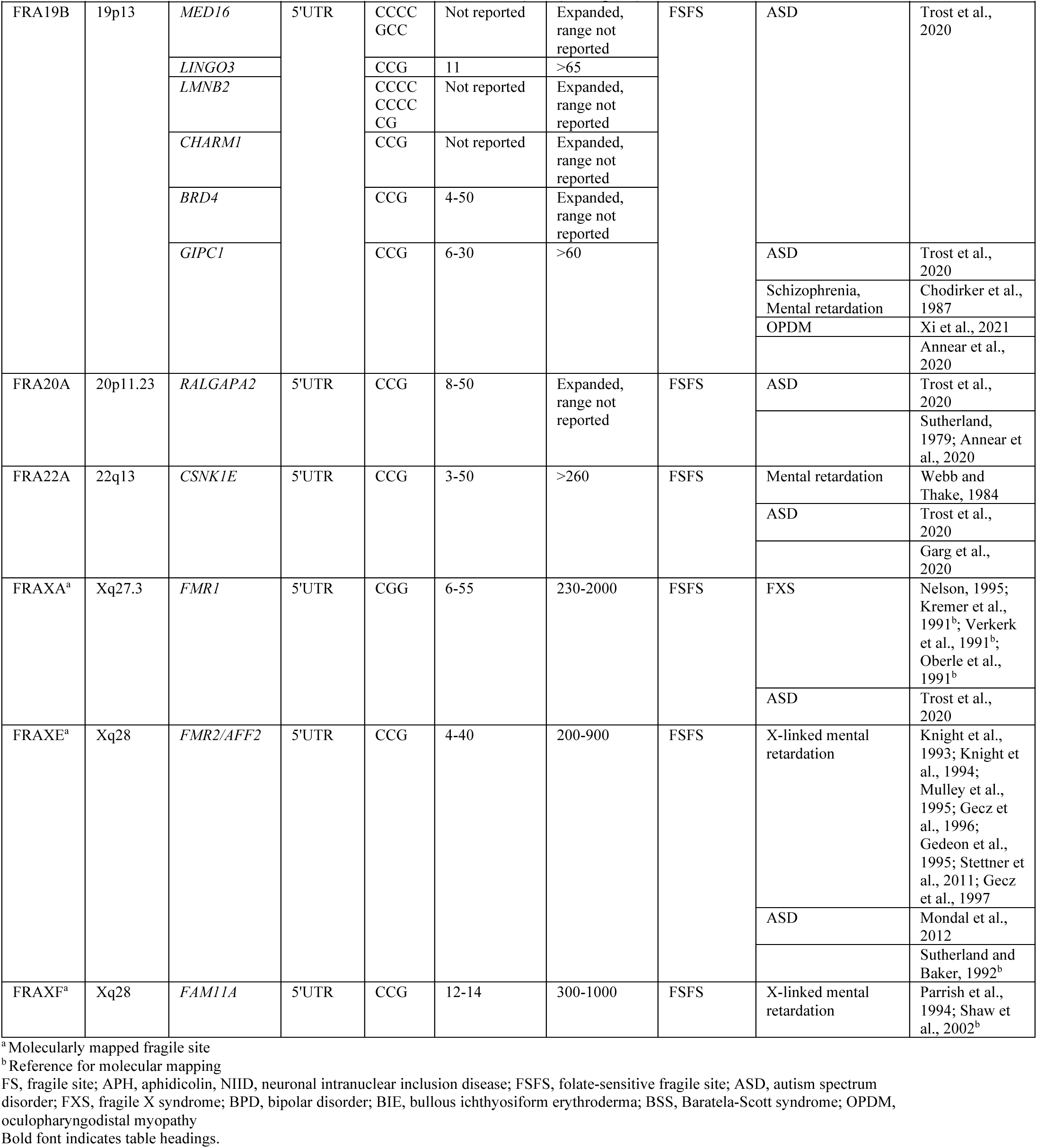
GC-rich expansions overlap with fragile sites and disease-associated loci.

